# Biomaterial preservation of mesenchymal stem/stromal cell fitness improves haematopoietic support in a post-leukaemic bone marrow niche model

**DOI:** 10.64898/2026.07.30.741738

**Authors:** Ioannis A. Tsigkos, Yusuf Ayten, Penelope M. Tsimbouri, Massimo Vassalli, Manuel Salmeron-Sanchez, Matthew J. Dalby

## Abstract

Mesenchymal stem/stromal cells (MSCs) are promising regenerative partners to haematopoietic stem cell (HSC) transplantation used for treatment of blood cancers. However, prolonged ex vivo expansion induces senescence and phenotypic drift that may compromise therapeutic efficacy. Here, we engineered a poly(ethyl acrylate)-laminin interface to preserve MSC fitness during expansion and investigated whether this biomaterial-controlled state influences function in a bioengineered post-leukaemic bone marrow niche model. Culture on the polymer-laminin interface enhanced MSC expansion while reducing senescence, DNA damage, and oxidative stress-associated phenotypes. Using a polyethylene glycol (PEG) hydrogel-based three-dimensional model of acute myeloid leukaemia (AML), we show that leukaemic remodelling induces inflammatory and stromal changes that are only partially reversed by chemotherapy. Introduction of biomaterial-expanded MSCs promoted recovery of the damaged niche, attenuated disease-associated stromal signalling, and enhanced HSC retention, haematopoietic progenitor output, and long-term multilineage potential. Together, these findings establish a direct link between biomaterial-guided MSC manufacturing and subsequent haematopoietic support, while providing a humanised platform for evaluating regenerative strategies following AML treatment.

**Graphical abstract:** 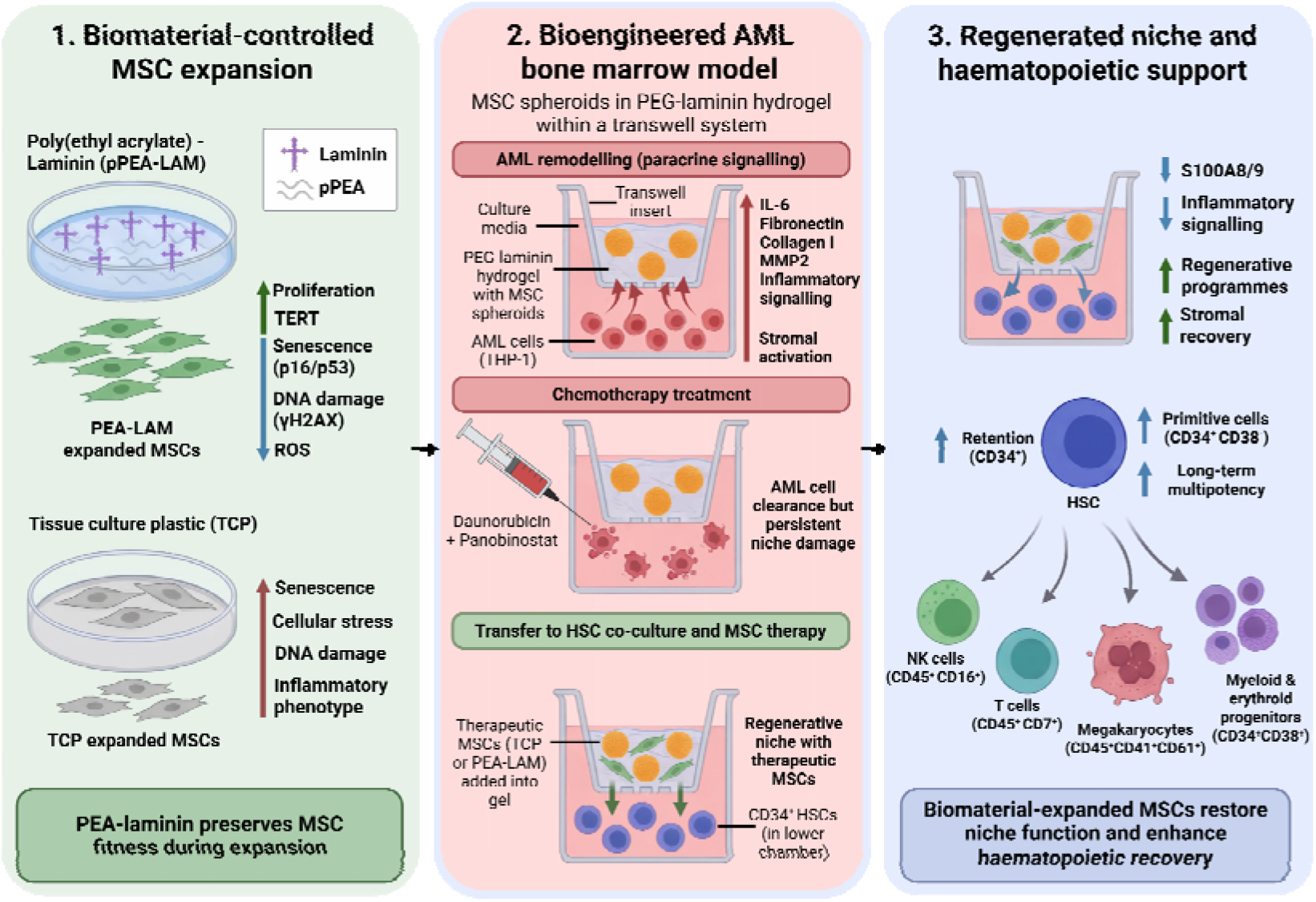

Created in BioRender. Tsigkos, I. (2027) https://BioRender.com/3wo7e1y

**Highlights:**

- Polymer-laminin interfaces maintain mesenchymal stromal/stem cell fitness and limit culture-induced senescence
- Engineered leukaemic niches reproduce disease-associated stromal remodelling and inflammation
- Biomaterial-expanded mesenchymal stromal/stem cells restore regenerative functions in post-treatment bone marrow
- MSC fitness preservation improves haematopoietic stem cell retention and multilineage haematopoiesis

**The bigger picture:** Acute myeloid leukaemia (AML) develops within the bone marrow, where cancer cells actively remodel the surrounding tissue to create a microenvironment that supports disease progression and protects leukaemic cells from therapy. Although chemotherapy can eliminate much of the tumour burden, it also leaves behind a damaged bone marrow niche that may provide suboptimal support for the haematopoietic stem cells (HSCs) required to rebuild the blood system following treatment or transplantation. Strategies that can restore healthy bone marrow function after therapy therefore represent an important but underexplored opportunity to improve patient recovery

Here, we investigate whether biomaterials can be used to improve the regenerative potential of mesenchymal stem/stromal cells (MSCs), a cell type that plays a key role in maintaining the bone marrow microenvironment. We show that expanding MSCs on a poly(ethyl acrylate)-laminin interface preserves a healthier, less senescent cell state during prolonged culture. We then evaluate these cells in a humanised three-dimensional model that reproduces AML-associated niche remodelling, chemotherapy treatment, and subsequent regeneration. Biomaterial-expanded MSCs promoted recovery of the damaged stromal environment and improved support of HSC survival, retention, and multilineage blood-cell production

These findings demonstrate that the conditions used to manufacture regenerative cell therapies can directly influence their performance after administration. More broadly, this work highlights how biomaterial-guided cell manufacturing can be combined with human microphysiological disease models (new approach methodologies, NAMs) to evaluate regenerative therapies in clinically relevant settings. Such approaches may help accelerate development of therapies that restore tissue function after cancer treatment while reducing dependence on animal models

## Introduction

Leukaemia refers to a group of haematological malignancies that affect the blood and bone marrow (BM). Leukaemia can be classified as either myeloid or lymphoid depending on the cell from which the disease originated^1^. Acute myeloid leukaemia (AML) is linked to ageing and is a particularly aggressive blood cancer, characterised by poor survival outcomes^2^. 7+3 chemotherapy (days 1-3 daunorubicin + days 1-7 cytarabine) has remained as the first-line treatment for more than 40 years^3^. Notably, disease recurrence occurs in approximately 60% of patients treated with chemotherapy alone^4^. Use of haematopoietic stem cell (HSC) transplant (HSCT) post-chemotherapy provides the only therapeutic option^4^ due to the graft vs leukaemia effect, where donor-derived T cells and natural killer cells eliminate residual AML cells^5^.

A key step following HSCT is the engraftment of transplanted HSCs, commonly defined by recovery of the absolute neutrophil count^6^. Primary graft failure is a life-threatening complication, for which a second HSCT may be used as a rescue procedure, although long-term success remains limited^7,8^. In a recent cohort of AML patients with primary graft failure, one-year overall survival was 55.5%^7^. Despite high engraftment rates, disease relapse remains a major challenge following HSCT and contributes substantially to treatment failure^7,8^. Therefore, preventing graft rejection^9^ and maximising graft vs leukaemia effect^5^ are key to improving patient outcomes.

During health, HSCs are supported by stromal cells, such as mesenchymal stromal (or stem) cells (MSCs) in the BM^10^. MSCs contribute extracellular matrix to the BM stem cell niche and secrete paracrine factors that support HSC maintenance and function^10^. When cancer evolves within the BM, the cancerous HSCs are also supported by the MSCs, and these stromal cells remodel the BM niche to support diseased cells rather than healthy cells, forming a tumour microenvironment (TME)^11^. This remodelling provides metabolic support to AML cells and helps them to evade treatment^12^. As with failed engraftment, relapse post-chemotherapy and HSCT is very serious as the cells are more likely to be refractory to treatment and, therefore, the outlook for the patient is very dismal.

Here, we propose that MSCs, used in conjunction with HSCs, as regenerative partners, can help BM niche remodelling to support health and improve HSC support following transplantation. MSCs, widely known for their regenerative potential, also have the ability to modulate the immune system to help with engraftment^13,14^. However, for cell therapies, they are massively expanded *in vitro*. This expansion leads to changes in phenotype^15^. Many early clinical trials with MSCs show efficacy; however, as the cells are expanded more to meet the demands of later-stage trials, this efficacy drops off and can be attributed to a regenerative secretome becoming a more negative, inflammatory, secretome^16,17^. Here, we use a biomaterials-mechanobiological approach to controlling MSC phenotype during lab expansion. This is based on adoption of poly(ethylacrylate) (PEA) that is known to spontaneously reveal cryptic cell adhesion and growth factor binding sites as extracellular matrix (ECM) proteins such as fibronectin (FN) and laminin (LAM) adsorb to the polymer^18–20^. Post PEA-extracellular matrix expansion, we introduce our culture-enhanced MSCs vs MSCs that have undergone standard lab expansion alongside HSC grafts in our AML niches, to mimic HSCT post chemotherapy.

Despite the clinical importance of engraftment, there are limited physiologically relevant *in vitro* models that allow investigation of HSC support and retention within the leukaemic BM niche^14^. Most current studies rely heavily on animal models, which do not fully recapitulate human disease and are associated with ethical and translational limitations. Therefore, developing a bioengineered humanised model of HSC support following transplantation is essential to better understand how the leukaemic niche influences transplant success. In this study, we develop a simple bioengineered model of AML to test our hypothesis.

## Results

We will start by demonstrating that polymer interfaces can be used to enhance MSC growth before introducing our bioengineered model of AML. We will then discuss the introduction of the enhanced MSCs into the model, as partner cells to HSCs, to study effects on HSC support and retention.

### Polymer-extracellular matrix interfaces for MSC growth

Our first goal is to demonstrate the ability to grow high-quality MSCs, without phenotypical drift or senescence that is typical of prolonged culture on tissue culture plastic (TCP)^21^. Mechanobiological research using nanoscale topography has shown that MSC phenotype is tightly linked to adhesion and intracellular tension, with slightly lowered intracellular tension (from that on tissue culture plastic) retaining the MSC phenotype^13,22^. A problem with the use of precise nanotopographies is the lack of ability to pattern large areas; this is a limitation if our ambition is to underpin the research on scalable technologies. Polyethylacrylate (PEA) is a polymer that can be plasma polymerised (pPEA) on large and complex surfaces^23^. As extracellular matrix (ECM) proteins, such as FN and LAM, are adsorbed onto PEA, they spontaneously reveal cryptic ^19^ cell adhesion and growth factor binding sites^20^. Use of FN with PEA has been associated with increased cell adhesion and intracellular tension^20^. However, recent work shows that while LAM increases cell adhesion, it shields cells from intracellular tension^24^.

We coated TCP with pPEA and used X-ray photoelectron spectroscopy (XPS) to confirm successful PEA coating. There were slight shifts in the binding energies compared to the EA monomer, likely due to crosslinking and functional group loss during polymerisation^23^. pPEA-coated samples showed distinct peaks for C-C (284 eV), C=O (531 eV), and C-O-C (533 eV), while the TCP polystyrene control displayed a peak at 292 eV (π-π aromatic interactions), which was absent in PEA, confirming effective surface coating (Figure 1A). Fluorescent analysis showed good adsorption of both FN and LAM to both uncoated TCP control and pPEA surfaces; both were reduced in adsorption from pPEA to TCP (**Figure S1A**, B). Micro bicinchoninic acid assay (BCA) revealed no significant difference in total ECM adsorption between pPEA and TCP (Figure 1B, C).

**Figure 1.**
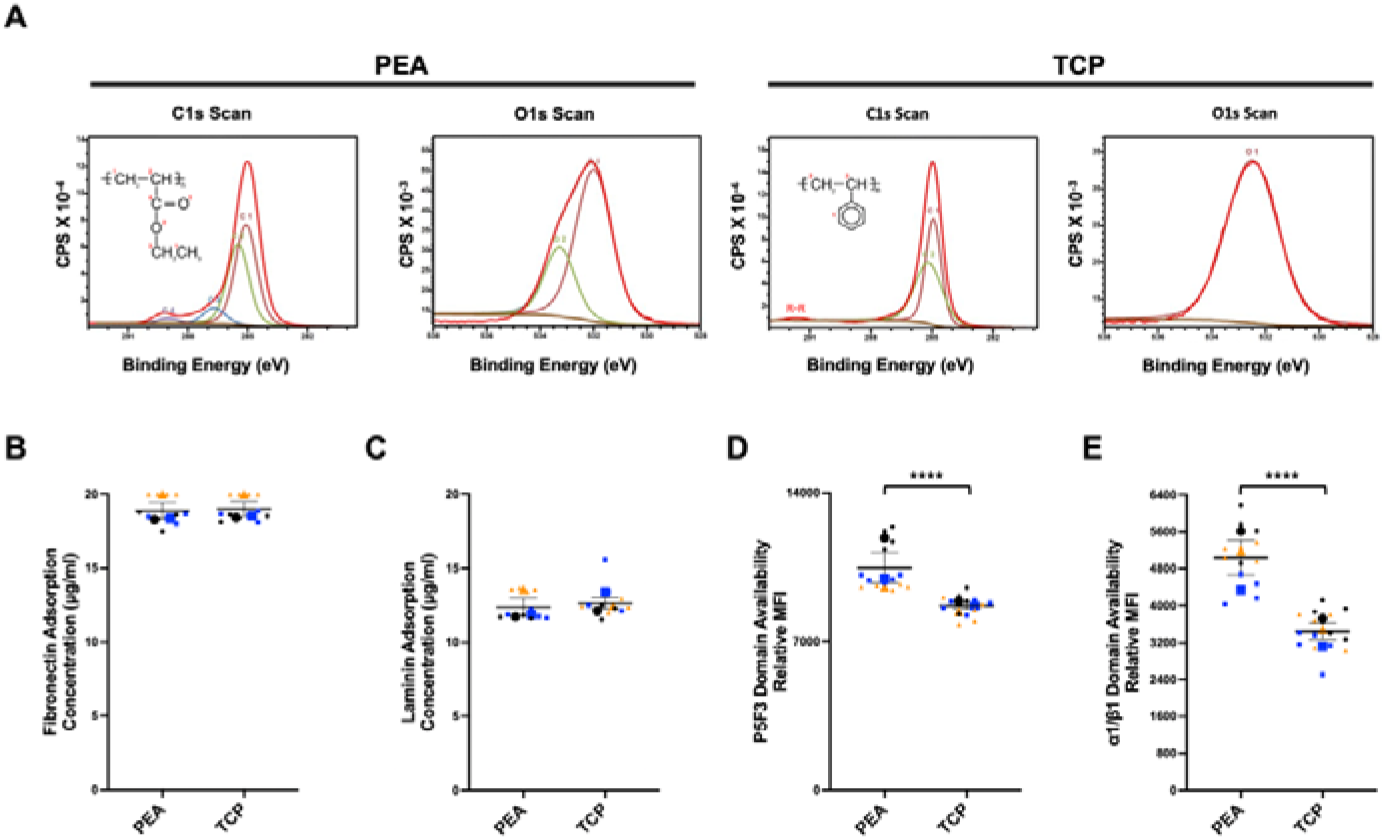
Characterisation of PEA-coated surfaces. Chemical composition of PEA-coated and TCP control surfaces (**A**). BCA assay measuring total adsorbed FN (**B**) and LAM (**C**) protein contents, expressed as µg/mL. Quantification of fluorescence intensity of P5F3 (for FN) (**D**) and α1β1 (for LAM) (**E**) availability. The larger different-coloured symbols on super plots represent the mean of each condition for each experiment. Smaller symbols represent the technical replicates of every experiment, accordingly. Statistical analysis was performed via Student’s t-test. *: p ≤ 0.05, **: p ≤ 0.01, ***: p ≤ 0.001, ****: p ≤ 0.0001. Data is presented as (mean ± SEM) from 3 biological replicates (N=3).

Functional domain availability, assessed via immunofluorescent analysis, showed significantly higher availability of the promiscuous heparin growth factor binding domain for pPEA coated with FN (P5F3) and LAM (α1/β1-HEP) (Figure 1D, E), indicating enhanced functional domain presentation independent of total protein adsorption (Figure 1B, C, **Figure S1A**, B), tying in with previous work on PEA-FN^14^. Contact angle measurements were used to help confirm successful coatings (**Figure S1C**, D).

### MSCs’ growth is improved on ECM interfaces

For a successful cell therapy, many MSCs need to be grown without unwanted senescence, which stops growth and generates a negative, inflammatory, secretome (senescence-associated secretory phenotype, SASP) ^16,17^. Observation of cell numbers after 21 days of culture showed that MSCs grew significantly faster on ECM-coated pPEA compared to TCP (Figure 2A, **Figure S2A**).

**Figure 2.**
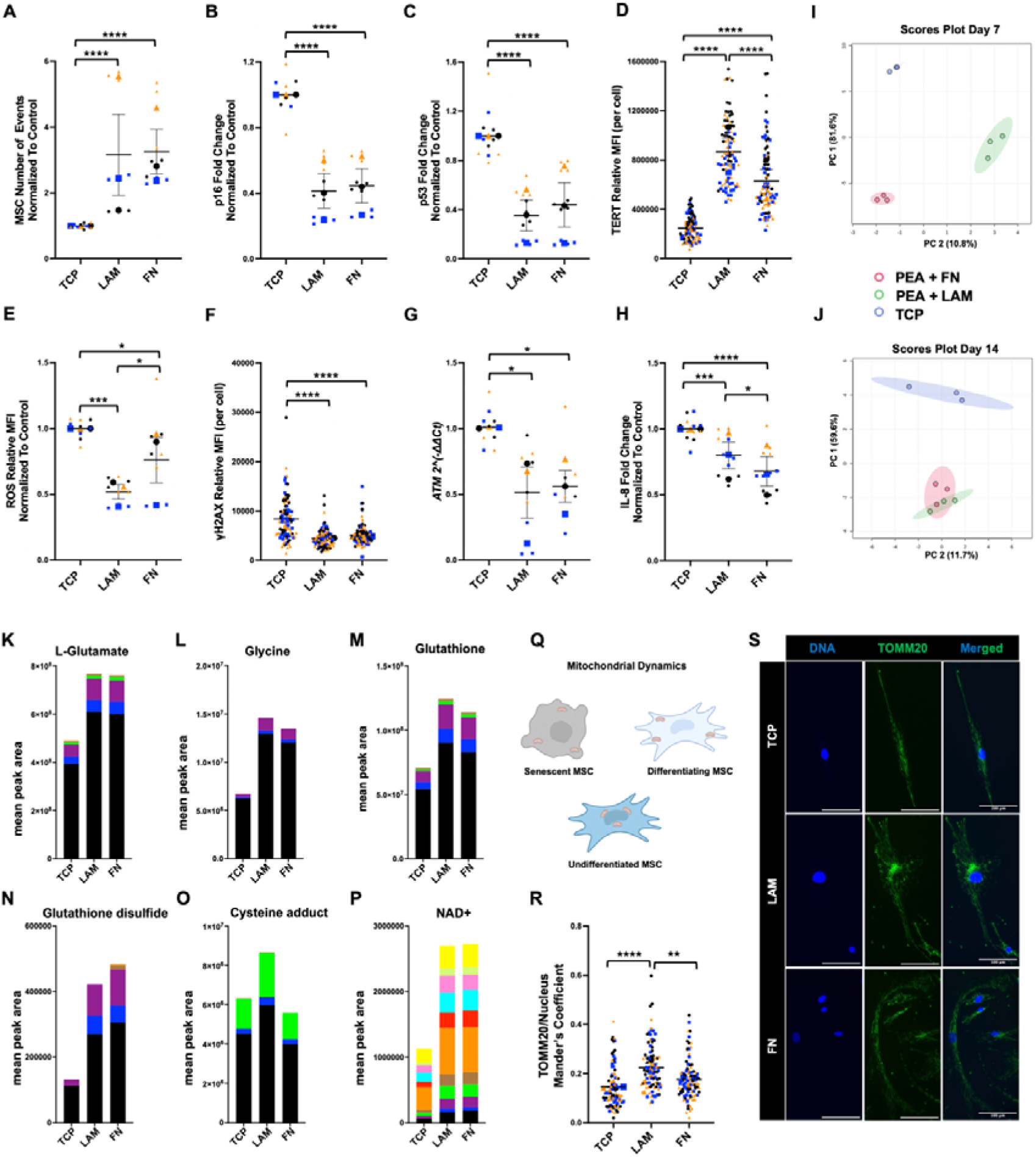
Investigating the effects of LAM, FN, and TCP on MSC senescence, metabolism and mitochondrial localisation. Quantification of the number of MSCs by flow cytometry (**A**). Protein expression of p16 (**B**) and p53 (**C**). Measurement of nuclear TERT MFI (**D**). Quantification of ROS median fluorescence intensity (MFI) (**E**). Measurement of γH2AX MFI (**F**). Gene expression of ATM (**G**). Protein expression of IL-8 (**H**). PCA plots visualising metabolomic clustering among TCP, pPEA-LAM and p-PEA-FN conditions, with PC1 and PC2 explaining 81.6% and 10.8% of the total variance, respectively (day 7), (**I**). PCA plots visualising metabolomic clustering among TCP, pPEA-LAM and p-PEA-FN conditions, with PC1 and PC2 explaining 59.6%, and 11.7%, of the total variance, respectively (day 14), (**J**). Representative stable isotope tracing of selected metabolites at day 14 (**K**-**P**). Corresponding isotopologue distributions following ^13^C -glucose labelling are comprehensively presented in (Figure S4). Schematic representation of mitochondrial dynamics in MSCs at different states (Created in BioRender. Tsigkos, I. (2027) https://BioRender.com/ohwpd9e) (**Q**). Measurement of TOMM20 localisation at day 3 (**R**). MSC staining for TOMM20. Blue and green stains represent the DNA, and protein of interest, respectively. Scale Bar = 100 µm. (**S**). The larger different-coloured symbols on super plots represent the mean of each condition for each experiment. Smaller symbols represent the technical replicates of every experiment, accordingly. Representative images out of 3 biological replicates. Statistical analysis was performed via One-way ANOVA with Tukey’s multiple comparisons. *: p ≤ 0.05, **: p ≤ 0.01, ***: p ≤ 0.001, ****: p ≤ 0.0001. Data is presented as (mean ± SEM) from 3 biological replicates (N=3).

Protein expression of senescence-associated growth inhibitors p16 and p53 was reduced on ECM-coated pPEA compared to TCP (Figure 2B-C, Representative images in **Figure S2B**) after 21 days of culture. We can see that this effect is integrin-related, as if we inhibit integrin subunits α6 (part of the laminin receptor) ^25^ and α5 (part of the fibronectin receptor)^26^ on pPEA-LAM or pPEA-FN, increased expression of p53 is observed, indicating a return to senescent phenotype (Figure S3). As cells senesce, telomeres shorten unless maintained by telomerase (TERT)^27^. TERT expression was significantly higher on the ECM-coated pPEA compared to TCP, but also significantly higher on pPEA-LAM than pPEA-FN (Figure 2D, Representative images in **Figure S2C**). Reactive oxygen species (ROS) are related to cell stress that can drive senescence; ROS was seen to be reduced on the ECM-coated pPEA compared to TCP (Figure 2E, Gating strategy in **Figure S2D**).

DNA damage is another driver of senescence^28^. Therefore, γH2AX, an early marker of DNA damage (Figure 2F, Representative images in **Figure S2E**), and ataxia-telangiectasia mutated (ATM)-kinase (Figure 2G) and ataxia telangiectasia and Rad3-related protein (ATR)-kinase (**Figure S2F**) as early-stage DNA damage initiators^29^ were investigated. Expression of γH2AX, ATM-kinase and ATR-kinase was reduced for ECM-coated pPEA compared to TCP (non-significant for ATR-kinase).

Expression of pro-inflammatory cytokines is associated with senescence^30^. IL-8 was seen to be reduced on PEA-adsorbed ECM proteins compared to TCP after 21 days of culture (Figure 2H).

Next, use of untargeted metabolomics showed that MSCs on TCP grouped very differently from MSCs on ECM proteins coated on pPEA at day 7 (Figure 2I and **Figure S4A**) and 14 of culture (Figure 2J and **Figure S4B**); we note that clear differences were also apparent between pPEA-LAM and pPEA-FN. ^13^C glucose labelling was next used to look at metabolism changes after 14 days of culture. ECM-coated pPEA conditions showed increased synthesis of L-glutamate and glycine, key precursors for glutathione (GSH) production, as well as elevated synthesis of GSH itself and its oxidised form, glutathione disulfide (GSSG) (Figure 2K-N, **Figure S4C** to see all control groups)^31^. The GSH pathway is a key antioxidant pathway used by cells to reduce cellular stress generated by ROS^32^.

Increased formation of cysteine adducts was also seen on ECM-coated pPEA conditions compared to TCP (Figure 2O, **Figure S4C**). These are temporary storage forms of cysteine activated in response to oxidative stress, again suggesting an enhanced redox-buffering system with the use of the ECM interface^33^. Lastly, increased NAD⁺ levels in MSCs cultured on ECM-coated pPEA compared with TCP (Figure 2P, **Figure S4C**). As NAD⁺ supports mitochondrial maintenance and DNA repair, its elevation suggests enhanced protection against mitochondrial ageing and genomic damage^34^.

The drivers of metabolism, the mitochondria, are known to relocate depending on whether the MSC is undifferentiated, differentiating or senescent (Figure 2Q). Peripheral mitochondrial localisation indicates an OXPHOS-active phenotype or senescence, while perinuclear clustering is associated with quiescence^35^, often seen in young or undifferentiated MSCs. After 3 days in culture, mitochondrial distribution was seen to be more perinuclear in MSCs cultured on pPEA-LAM compared to those on pPEA-FN or TCP (Figure 2R, S, Figure S5). By 21 days of culture, both ECM-coated PEA substrates supported MSCs with more perinuclear mitochondria than TCP (**Figure S6D**, Representative images in **Figure S6B**-C).

Together, the data show that with ECM-coated pPEA, more cells are grown in the same time compared to TCP, with lower stress and senescence and with greater cytoprotective advantage. These changes are important if our aim is to produce more, higher-quality MSCs for cellular therapies.

### Hypoxia further enhances MSC phenotype but with reduced growth and viability

Having shown that the PEA-ECM interfaces offer better growth and phenotype than TCP, we next considered hypoxia. The BM is a hypoxic environment with spatially heterogeneous oxygen tension^36^ and, therefore, we compared MSC phenotype on PEA-FN/LAM substrates under normoxia (21% O₂) and hypoxia (5% O₂). The architecture of the BM stem cell niche is complex, with MSCs and HSCs forming a part of each other’s niche^10^. In general, close to the endosteal bone lining, surface, periarteriolar MSCs (nestin^bright^/NG2⁺) are linked to quiescent HSCs, whereas perisinusoidal MSCs (nestin^dim^/Leptin receptor (LepR)⁺) provide key secretory support via stem cell factor (SCF, promotes stem cell survival, proliferation, differentiation, and migration) and stromal cell-derived factor-1 (CXCL12, a chemokine involved in stem cell migration) to differentiating HSCs^37–39^. Other cells, such as CXCL12-abundant reticular cells (CAR cells), express CXCL12 at the sinusoids, and osteoblasts contribute thrombopoietin (TPO, involved in HSC growth and differentiation) and angiopoietin (Angpt-1, promotes HSC maintenance) at the endosteal surface^37–39^ (see schematic in Figure 3A).

**Figure 3.**
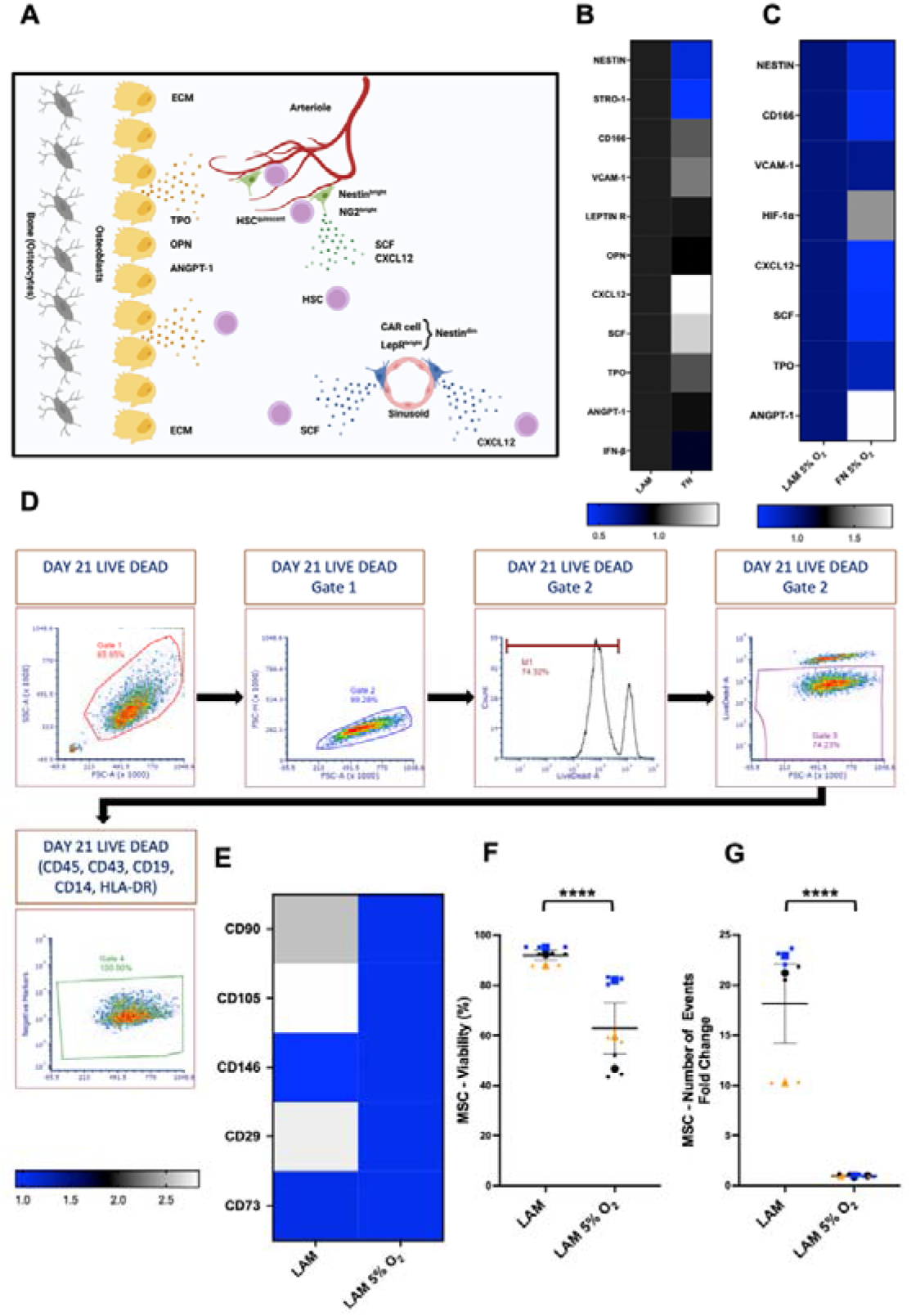
Expression of stromal and stemness markers on MSCs under normoxia or hypoxia. Graphical representation of the stromal cell localisation, marker expression and secretory profile, in the BM niche (Created in BioRender. Tsigkos, I. (2027) https://BioRender.com/90bkba4). (**A**). Protein expression of stromal markers on pPEA-LAM and pPEA-FN under normoxic conditions (**B**). Protein expression of stromal markers on pPEA-LAM and pPEA-FN under hypoxic conditions (**C**). Gating strategy for flow cytometry. After single-cell selection, the dead cells are excluded (dead cell curve peak 10^5^) from the consequent evaluation of marker expression (**D**). Expression of CD73, CD29, CD146, CD105, and CD90 (**E**). MSC viability (**F**) and number of viable cell-associated events following gating (**G**). Data is presented as heatmaps from 3 biological replicates. Coloured scale bar: white indicates an upregulation while blue a downregulation, (N=3). The different-coloured symbols on super plots represent the mean of each condition for each experiment. Smaller symbols represent the technical replicates of every experiment, accordingly. Statistical analysis was performed via Student’s t-test. *: p ≤ 0.05, **: p ≤ 0.01, ***: p ≤ 0.001, ****: p ≤ 0.0001. Data is presented as (mean ± SEM) from 3 biological replicates (N=3).

Under normoxia, MSCs on PEA-LAM showed increased nestin and Stro-1 expression, resembling periarteriolar stroma (HSC support phenotype)^38,40,41^, while PEA-FN supported MSCs with an increased CXCL12 and SCF phenotype, consistent with perisinusoidal features (HSC differentiation phenotype)^10^ (Figure 3B). PEA-FN also showed moderate increases in CD166 (ALCAM, an MSC marker), VCAM-1 (vascular cell adhesion molecule, used by HSCs to interact with MSCs), and TPO (Figure 3B).

Under hypoxia, PEA-LAM further upregulated nestin, CD166, VCAM-1, CXCL12, SCF, and TPO, while PEA-FN showed higher HIF-1α (hypoxia-inducible factor) and Angpt-1 (Figure 3C). These data suggest that PEA-LAM with hypoxia promoted an overall superior stromal support and phenotype, while PEA-FN with hypoxia perhaps produces a more HSC quiescence-supporting phenotype.

As PEA-LAM gave the most arteriolar phenotype in normoxia, and supported further niche markers in hypoxia, we next compared PEA-LAM in normoxia and hypoxia against ISCT (International Society for Cell and Gene Therapy)^42^ MSC markers (Figure 3D-E, Histograms in Figure S7). ISCT marker analysis showed higher CD90, CD105, and CD29 for MSCs on PEA-LAM under normoxia, while CD73 was unchanged, and CD146 was slightly reduced. Normoxia also massively enhanced viability and proliferation (Figure 3F-G).

In summary, MSCs grew significantly faster (Figure 2A) on PEA-ECM matrices, with lowered senescence. PEA-LAM, in particular, retained high TERT levels (Figure 2D), more perinuclear mitochondria (Figure 2R, S, Figure S6) and expressed the highest levels of periarteriolar/endosteal niche markers that are used to maintain HSCs (Figure 3B). Hypoxia further enhanced expression of niche factors, but at massive detriment to MSC viability and, critically, growth (Figure 3E-F). In designing a cell therapy, this detriment to growth and viability would be a key consideration for a GMP cell manufacturer as they need to preserve all viability, phenotype and growth for successful manufacture^17^. Therefore, we selected not to pursue hypoxic conditions and to select PEA-LAM as our optimal condition for MSC growth.

### Development and characterisation of the AML niche

Next, we wanted to develop a 3D *in vitro* leukaemic BM model in which to test our culture enhanced MSCs. Polyethylene glycol (PEG) hydrogels were selected to mimic the hydrogel character of the BM, as the use of maleimide functionalisation can allow Michael’s type addition gelation and incorporation of functional groups, such as ECM proteins and degradable crosslinks^43^. 10% PEG-maleimide (PEG-MAL) was crosslinked with PEG-dithiol (PEG-SH) and the matrix metalloprotease (MMP) degradable VPM (GCRD**<u>VPM</u>**SMRGGDRCG) peptide (75:25 PEG-SH:VPM molar ratio) to have a gel with BM-like stiffness of ∼7.5 kPa, the physiological range of BM being ∼2-35 kPa^44^ (Figure 4A-C), from which we can recover cells^43^. For enhanced biocompatibility, we incorporated 0.1 mg/mL PEGylated LAM into the hydrogels, giving a stiffness of ∼6.5 kPa (Figure 4C).

**Figure 4.**
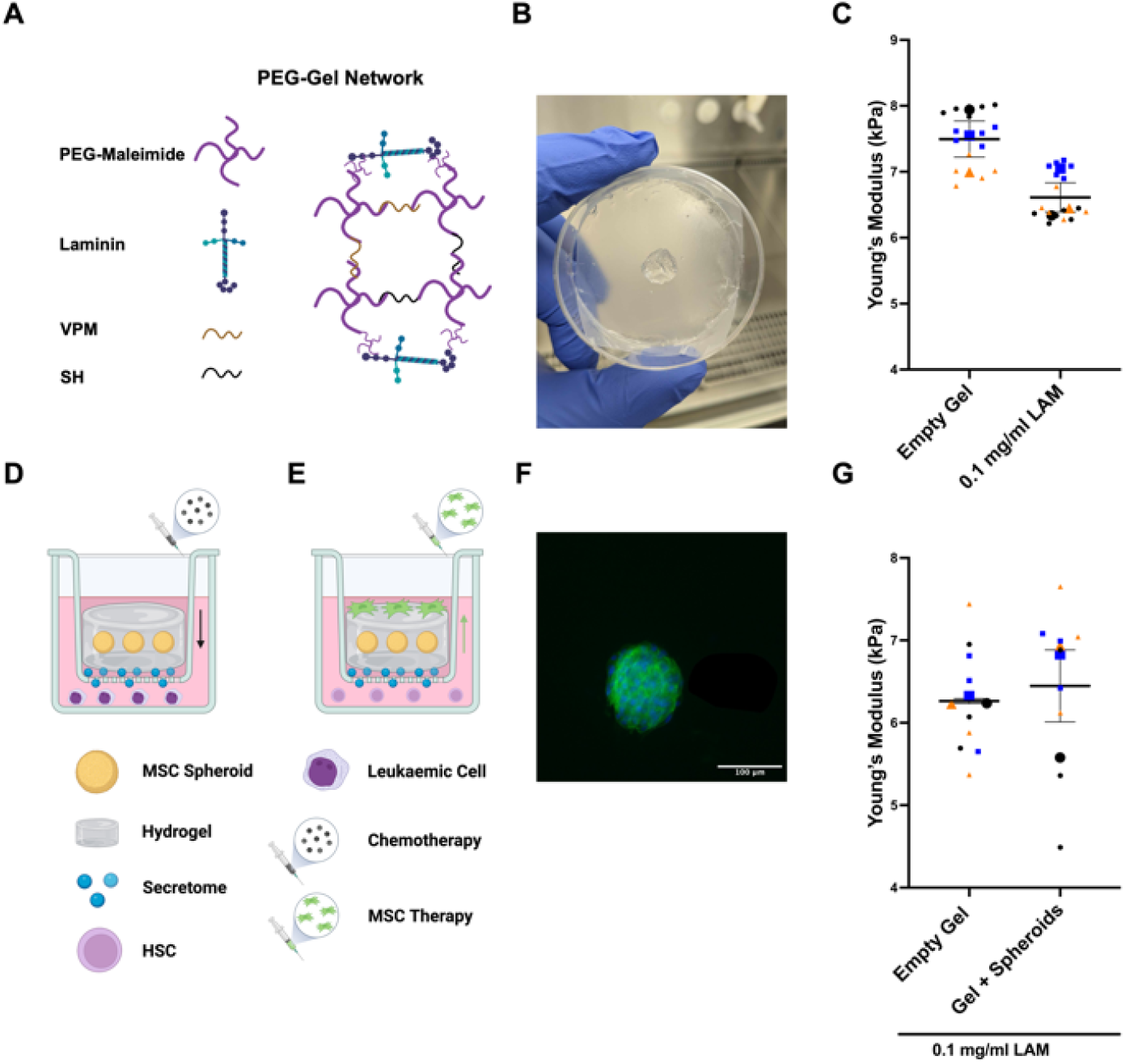
Biophysical characterisation of PEG-MAL hydrogel functionalisation and mechanics. Schematic representation of PEG-maleimide gel networks upon PEGylation with LAM (Created in BioRender. Tsigkos, I. (2027) https://BioRender.com/5gkf4q9) (**A**). Macroscopic image of polymerised PEG-maleimide hydrogel (**B**). Measurement of gel stiffness after LAM-PEGylation (**C**). Schematic representation of the transwell-based AML niche (**D**) and subsequent regenerative treatment workflow (Created in BioRender. Tsigkos, I. (2027) https://BioRender.com/k7d1yz1). (**E**). Representative MSC spheroid generated following 24 h culture in AggreWell™400 plates and subsequently incorporated within hydrogels. Green and blue stains represent the F-actin and DNA, respectively. Scale Bar = 100 µm. (**F**). Measurement of gel stiffness after incorporation of spheroids (**G**).

Our design requires the hydrogel to contain healthy stromal cells, which can then be exposed to AML cells to remodel the BM environment and where chemotherapeutics can be added (Figure 4D). Once remodelled and treated, we need a design where cell therapies (culture-enhanced MSCs along with HSCs to test haematopoietic support) can be added (Figure 4E). We chose to incorporate MSC spheroids (Figure 4F) (∼100 cells per spheroid created in AggreWell^TM^400 24-well plates) into the gels (∼100 spheroids per 100 µL of gel) within a transwell system; this allowed us to: (1) place the niches into environments containing cancerous or healthy haematopoietic cells where secretome was shared; (2) add chemotherapeutics / our culture enhanced MSCs/HSCs; and (3) easily separate the MSCs from the leukaemic or haematopoietic cells for analysis, where desirable (e.g. RNAseq or flow cytometry) to study stromal response alone or HSC progeny (Figure 4D, E). We note that addition of the spheroids did not alter the stiffness of the PEG-LAM gels (Figure 4G).

When the BM models were introduced to control or THP-1, a cell line derived from acute myeloid leukaemia (AML) cultures, in wells (Figure 5A), spheroid viability remained high (Figure 5A, B). However, with THP-1 cell exposure, the spheroids exhibited morphological changes consistent with activation and disaggregation (Figure 5B-G, **Figure S8A**). ELISA showed significantly increased secretion of IL-6, FN, and pro-collagen Iα1 (Figure 5H-J) in co-culture compared to MSC monoculture. Zymography for active MMP2 and 9 showed increased active MMP2 in the coculture (Figure 5K-L).

**Figure 5.**
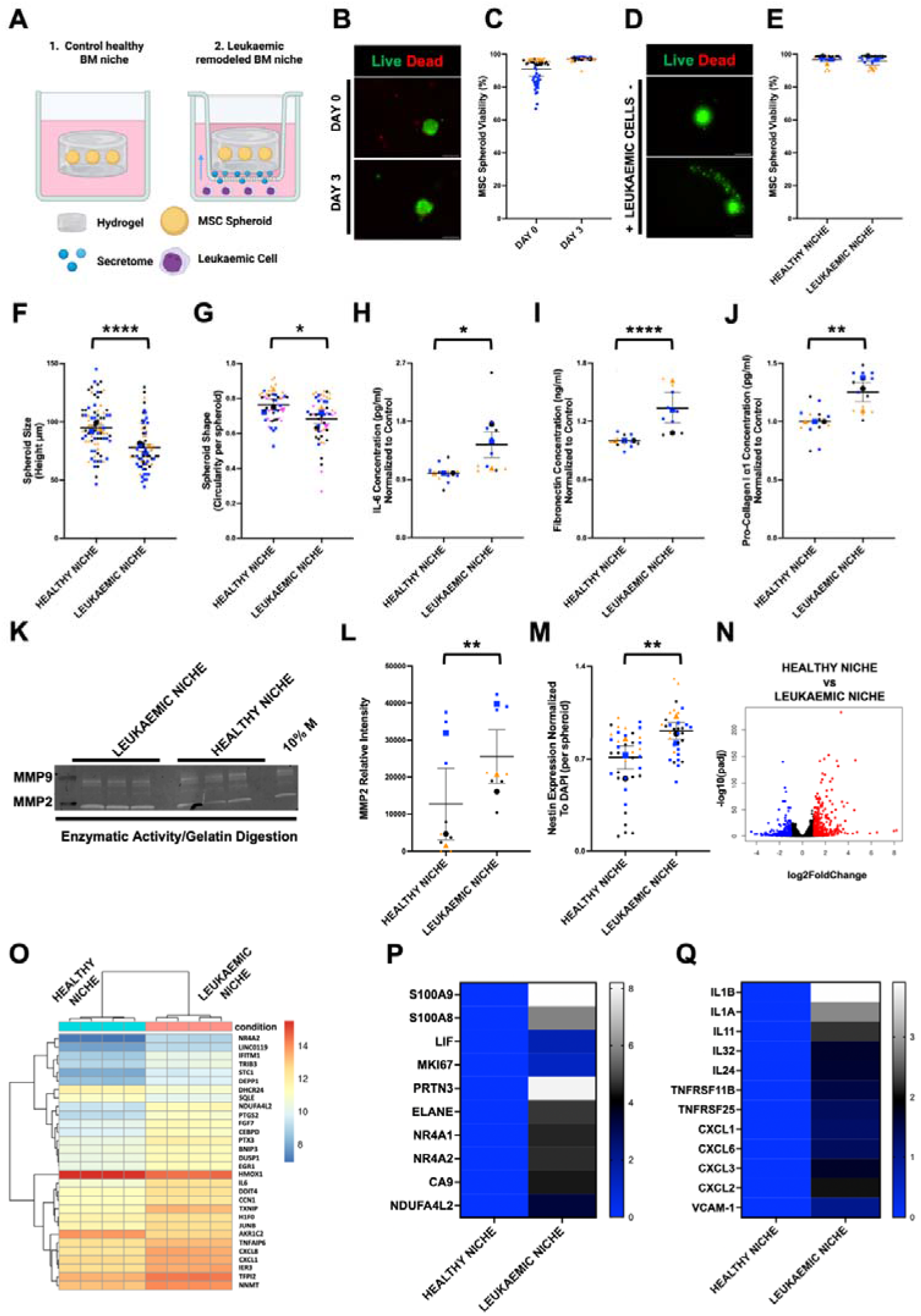
Development and characterisation of the leukaemic BM niche. Schematic representation of leukaemic niche development (Created in BioRender. Tsigkos, I. (2027) https://BioRender.com/ec77exr). (**A**). Live-dead staining of spheroids for day 0 and day 3. Scale Bar = 100 µm (**B**). Quantification of viability for day 0 and day 3 (**C**). Live-dead staining of spheroids for day 6 after THP-1 introduction. Scale Bar = 100 µm (**D**). Quantification of viability for day 6 after THP-1 introduction (**E**). Measurement of spheroid size (**F**) and shape (**G**). Measurement of IL-6 (**H**), FN (**I**), and pro-collagen Iα1 (**J**) secretion via ELISA. Greyscale image of zymogram gel (**K**). Quantification of MMP proteolytic cleavage (**L**). Quantification of spheroids stained for nestin (MFI) (**M**). The global transcriptional change across the groups compared was visualised by a volcano plot. Each data point in the scatter plot represents a gene. The log_2_ fold change of each gene is represented on the x-axis and the -log_10_ of its adjusted p-value is on the y-axis. Genes with an adjusted p-value less than 0.05 and a log_2_ fold change greater than 1 are indicated by red dots. These represent up-regulated genes. Genes with an adjusted p-value less than 0.05 and a log_2_ fold change less than -1 are indicated by blue dots. These represent down-regulated genes (**N**). Bi-clustering heatmap illustrates the expression of the top 30 most differentially expressed genes, sorted by their adjusted p-value by plotting their log_2_ transformed expression values in samples. Red and blue indicate an upregulation or downregulation, respectively (**O**). Heatmaps illustrate the expression of selected genes, by plotting their log_2_ fold change values. White and blue indicate an upregulation or downregulation, respectively (**P**-**Q**). Data is presented from 4 technical replicates (n=4). The different coloured symbols on graphs represent the mean of every condition for every experiment. Smaller symbols represent the technical replicates of every experiment, accordingly. Representative images out of 3 technical replicates. Statistical analysis was performed via Student’s t-test or One-way ANOVA with Tukey’s multiple comparisons. *: p ≤ 0.05, **: p ≤ 0.01, ***: p ≤ 0.001, ****: p ≤ 0.0001. Data is presented as (mean ± SEM) from 3 biological replicates, (**B**-**M**) (N=3). Analysis was performed on four technical replicates derived from one biological sample, (**N**-**Q**), (n=4).

Further, in the BM, MSCs have been shown to increase nestin expression in response to AML developing, with the cancer cells killing nestin-negative MSCs and relying on nestin-positive cells for remodelling^12^. Quantification of nestin in the spheroids showed a significant increase with the addition of THP-1 cells to the culture (Figure 5M, **Figure S8B**).

Next, RNA-seq was performed to compare MSC spheroids within the PEG-LAM gels (healthy niche, no THP-1s) to MSC spheroids within the PEG-LAM gels in culture with THP-1 cells (leukaemic niche). Volcano plotting demonstrated large changes between the healthy and leukaemic niches (Figure 5N). Gene ontology (GO) analysis showed a number of increased differences in areas of inflammation, proliferation, migration (and chemotaxis) and ECM organisation, among others, for the MSCs in the leukaemic niche compared to those in the healthy niche (Figure S9). Genes (please see **table S1** for a list of genes we name, their full name and their function) associated with AML survival and chemoresistance (e.g., *CXCL1, CXCL8, IL-6, TNFAIP6*) ^45^ and genes involved in stress (*IER3, TXNIP, DDIT4, ATF4, JUNB, NR4A1/2*) ^46^ were significantly upregulated (Figure 5O). Focussed analysis revealed further up-regulations of genes involved in immune response (*S100A8, S100A9, PRTN3, ELANE*)^47^, cell proliferation (*MKI67*), hypoxia response (*CA9, NDUFA4L2*)^48^, stem cell development (*LIF, NR4A1, NR4A2*)^49^ (Figure 5P) and immune response and inflammation (*IL-1B, IL-1A, IL-11, IL-32, IL-24, TNFRSF11B, TNFSF25, CXCL1, CXCL6, CXCL3, CXCL2, VCAM-1*)^50,51^ (Figure 5Q).

Taken together, these data demonstrate spheroid activation and MSC remodelling under the control of the AML cells.

### Treating the niche with chemotherapy

Now that we have constructed a cancer remodelled niche (MSC spheroids in gels introduced to THP-1 cells), to complete the model, we also need to mimic successful treatment of the BM cancer niche. To achieve this, we needed an effective dose of chemotherapeutics with which to form the treated AML niche in which HSC support could be evaluated (Figure 6A). First, THP-1 monocultures were treated with daunorubicin with/without panobinostat. Lower drug doses (25-50 nM) showed cooperation and efficacy, and we note that high concentrations (up to 1 μM) did not add much more drug efficacy (**Figure S10A**). Dosing again on day 4 with fresh 200 nM drug doses also did not add significantly to drug efficacy (**Figure S10B**). Similarly, adding cytarabine (another front-line AML treatment) to the regimen for 2 or 4 days also failed to further enhance drug efficacy (**Figure S10D**-E). Please note that DMSO controls are shown in **Figure S10C**&F. As even the lowest concentrations tested were above the IC₅₀ for THP-1 cells, suggesting effective treatment, we opted to use 50 nM daunorubicin + 50 nM panobinostat (DP) for 2 days as our optimal dose that would provide minimal potential MSC toxicity.

**Figure 6.**
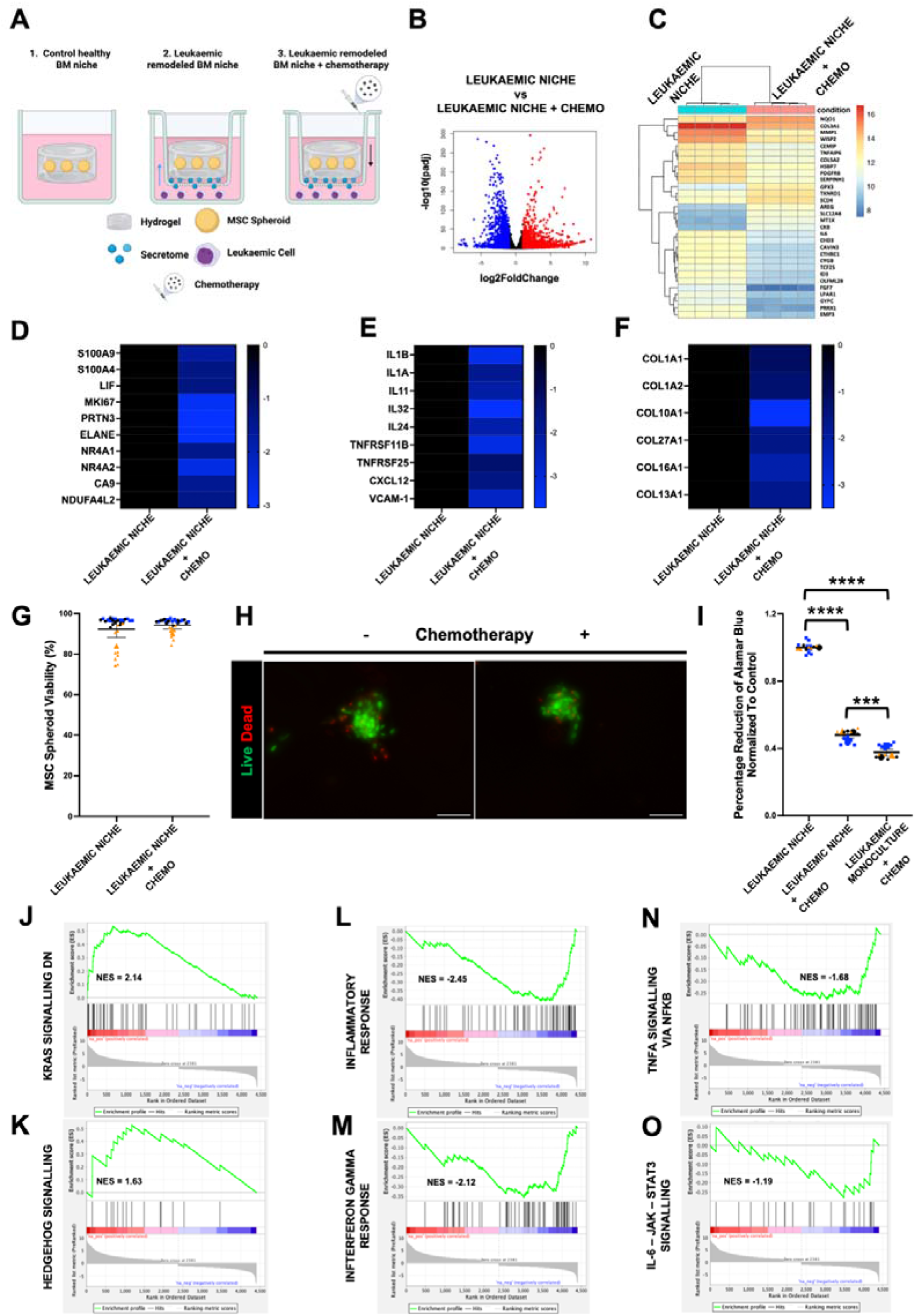
Treatment of the leukaemic BM niche with chemotherapy. Schematic representation of leukaemic niche treatment with chemotherapy and transcriptomic characterisation (Created in BioRender. Tsigkos, I. (2027) https://BioRender.com/b0y7k93). (**A**). The global transcriptional change across the groups compared was visualised by a volcano plot. Each data point in the scatter plot represents a gene. The log_2_ fold change of each gene is represented on the x-axis and the -log_10_ of its adjusted p-value is on the y-axis. Genes with an adjusted p-value less than 0.05 and a log_2_ fold change greater than 1 are indicated by red dots. These represent up-regulated genes. Genes with an adjusted p-value less than 0.05 and a log_2_ fold change less than -1 are indicated by blue dots. These represent down-regulated genes (**B**). Bi-clustering heatmap illustrates the expression of the top 30 most differentially expressed genes, sorted by their adjusted p-value by plotting their log_2_ transformed expression values in samples. Red and blue indicate an upregulation or downregulation, respectively (**C**). Heatmaps illustrate the expression of selected genes, by plotting their log_2_ fold change values. White and blue indicate an upregulation or downregulation, respectively (**D**-**F**). Data is presented from 4 technical replicates (n=4). Quantification of cell viability for DAY 8 after THP-1 introduction and chemotherapy treatment (**G**). Live-dead staining of spheroids. Scale Bar = 100 µm (**H**). Measurement of THP-1 metabolic activity with AlamarBlue (**I**). GSEA was performed on transcriptomic data comparing MSCs from leukaemic conditions to those from leukaemia□+□chemotherapy treatment. The enrichment plots highlight hallmark gene sets that are differentially regulated in response to chemotherapy. Each plot displays the running enrichment score (green line), positions of gene set members in the ranked gene list (black bars), and the ranking metric (grey). Red and blue shading indicate positive or negative correlations, respectively. Identified enrichment of KRAS signalling (downregulation-associated genes) (**J**) and hedgehog signalling (**K**) pathways. Identified negative enrichment of gene sets associated with inflammatory response (**L**) and interferon-γ response (**M**), TNF-α signalling via NF-κB (**N**), and IL6-JAK-STAT3 (**O**), post-chemotherapy. Data is presented from 4 technical replicates (n=4). The different coloured symbols on graphs represent the mean of every condition for every experiment. Smaller symbols represent the technical replicates of every experiment, accordingly. Representative images out of 3 technical replicates. Statistical analysis was performed via Student’s t-test or One-way ANOVA with Tukey’s multiple comparisons. *: p ≤ 0.05, **: p ≤ 0.01, ***: p ≤ 0.001, ****: p ≤ 0.0001. Data is presented as (mean ± SEM) from 3 biological replicates, (**G**-**I**), (N=3). Analysis was performed on four technical replicates derived from one biological sample, (**C**-**F** & **J**-**O**), (n=4).

For the MSCs in the model, we wanted to first check, ahead of adding chemotherapy, whether MSC culture as spheroids was associated with reduced growth-related activity (these therapies preferentially target rapidly dividing cells). Therefore, CellTiter-Glo 3D assays were performed to assess cellular ATP levels. MSCs were seeded either as monolayers or into spheroids at equal initial cell numbers. MSCs grown in monolayers exhibited significantly higher ATP-associated luminescence than MSCs cultured as spheroids (**Figure S10G**), with spheroids maintaining relatively stable luminescence between days 2 and 8 (**Figure S10H**). These findings allow us to progress checking MSC phenotype in response to THP-1 addition and survival in response to the chemotherapies.

Following DP treatment of the leukaemic niche, MSC gene expression patterns were surveyed by RNA-seq in order to observe stromal response to treatment. Transcriptional changes showed changes in both up- and down-regulated genes (Figure 6B). Patterns of up-regulated transcripts included antioxidant, cell defence-related genes (*MT1X, GPX3, TXNRD1, NQO1*)^52^ (Figure 6C, Please see **table S2** for a list of genes, their full name and their function). Patterns of down-regulated transcripts included those involved in proliferation and cell stress (*WISP2, SERPINH1, HSPB7*)^53^, chemoresistance (*IL-6, TNFAIP6, PDGFRB*)^54^, and ECM remodelling and membrane organisation (*COL3A1, COL5A2, MMP1, GYPC, EMP3, CAVIN3, OLFML2B, CTHRC1, CEMIP*)^55^ (Figure 6C). Gene groupings related to cancer-associated fibroblasts, proliferation, cytoprotection, and tumour microenvironment (Figure 6D, S100A9, *S100A4, LIF, MKI67, PRTN3, ELANE, NR4A1, NR4A2, CA9, NDUFA4L2*)^56^, tumour-stroma communication and bone remodelling (Figure 6E, *IL-1B, IL-1A, IL-11, IL-32, IL-24, TNFRSF11B, TNFRSF25, CXCL12, VCAM-1*)^57^ and ECM components linked to desmoplasia, the pathological hardening of cancer tissue, were suppressed (Figure 6F, *COL1A1, COL1A2, COL10A1, COL27A1, COL16A1, COL13A1*)^58^ (please see **table S2** for list of genes, their full name and their function). MSC spheroid viability following DP treatment was assessed, and remained high, indicating that the chemotherapy regimen did not compromise MSC survival within the niche model (Figure 6G-H).

In contrast, DP treatment significantly reduced THP-1 metabolic activity within the co-culture, demonstrating effective targeting of leukaemic cells while preserving the post-treatment stromal compartment (Figure 6I). However, THP-1 cells co-cultured with MSC spheroids retained significantly higher metabolic activity compared with THP-1 monocultures following treatment (Figure 6I). This suggests that MSCs provide a protective niche that partially preserves leukaemic cell viability under chemotherapy, consistent with microenvironment-mediated chemoresistance^59^. Moreover, gene set enrichment analysis (GSEA - a method to see if genes belonging to specific biological pathways change relative to control) was leveraged to give further information on the microenvironment. KRAS (Kirsten rat sarcoma virus) signalling DN and Hedgehog hallmark gene sets were positively enriched (Figure 6J–K, Figure S11), suggesting a transcriptional signature consistent with reduced KRAS signalling alongside altered Hedgehog signalling following chemotherapy, in a leukaemic microenvironment where reciprocal MSC-AML paracrine interactions can contribute to chemoresistance^60^. However, hallmarks of pathways associated with proliferation, inflammation and regeneration (TNF-α signalling via NF-κB, IL6-JAK-STAT3, inflammatory response, interferon response)^61^ all exhibited negative enrichment (Figure 6L-O, Figure S11).

Taken together, the data illustrate that the MSCs do act to protect the THP-1 cells, through signalling and increased viability; offering niche protection. However, the chemotherapy is more broadly impacting the MSCs to reduce the preparation of the tumour microenvironment to support the THP-1 cells.

### RNA-seq characterisation of treated AML niches with addition of therapeutic MSCs

With our aim of demonstrating that high-quality MSCs can improve HSC support, we added MSCs that were pre-cultured on TCP (TCP-MSCs, as is standard) or on PEA-LAM (LAM-MSCs, to maintain naïve phenotype) into the niches via injection into the hydrogel matrix at a density of 50,000 cells per gel. This was performed post-DP chemotherapy and with the spheroid-containing hydrogels removed to wells without THP-1s, to represent successful treatment (Figure 7A). We note that therapeutic MSCs were expanded on either TCP or PEA-LAM for 21 days at an original seeding density of 2000 cells/cm^2^.

**Figure 7.**
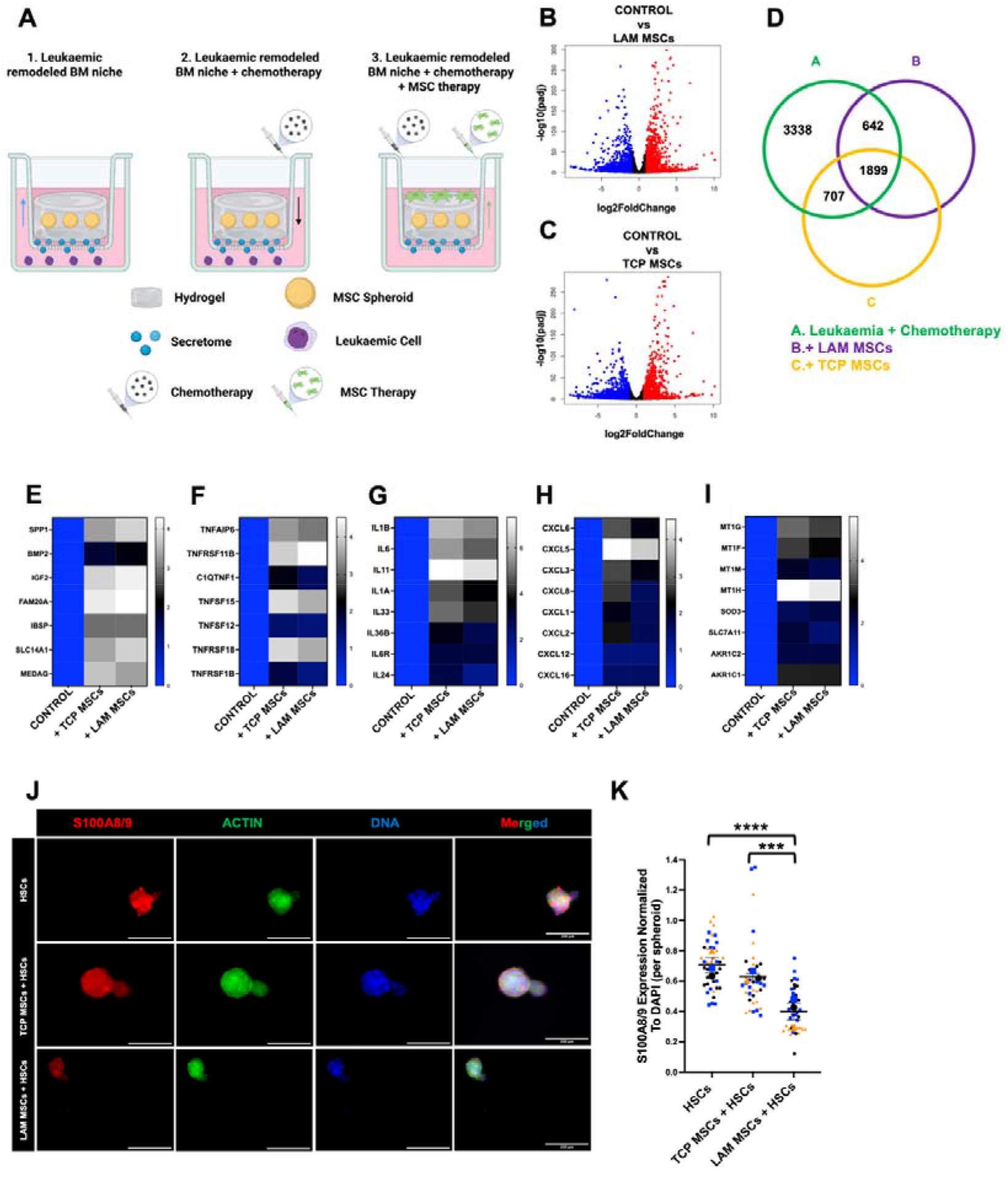
Characterisation of the chemotherapy-treated leukaemic BM niche upon its regeneration with therapeutic MSCs. Schematic representation of experimental design (Created in BioRender. Tsigkos, I. (2027) https://BioRender.com/gyy3szm) (**A**). The global transcriptional change across the groups compared was visualised by a volcano plot. Each data point in the scatter plot represents a gene. The log_2_ fold change of each gene is represented on the x-axis and the -log_10_ of its adjusted p-value is on the y-axis. Genes with an adjusted p-value less than 0.05 and a log_2_ fold change greater than 1 are indicated by red dots. These represent up-regulated genes. Genes with an adjusted p-value less than 0.05 and a log_2_ fold change less than -1 are indicated by blue dots. These represent down-regulated genes (**B**-**C**). Venn diagram illustrating shared and condition-specific significant differentially expressed genes across leukaemia + chemotherapy, leukaemia + chemotherapy + LAM-MSC, and leukaemia + chemotherapy + TCP-MSC conditions, with each DEG set generated relative to the leukaemic niche. Green circle: leukaemia + chemotherapy; purple circle: leukaemia + chemotherapy + LAM-MSCs; orange circle: leukaemia + chemotherapy + TCP-MSCs (**D**). Heatmaps illustrate the expression of selected genes, by plotting their log_2_ fold change values. White and blue indicate an upregulation or downregulation, respectively (**E**-**I**). Data is presented from 4 technical replicates (n=4). Spheroid staining for S100A8/9 (**J**). Quantification of S100A8/9 (**K**). The different coloured symbols on super plots represent the mean of every condition for every experiment. Smaller symbols represent the technical replicates of every experiment, accordingly. Green, blue, and red stains represent the F-actin, DNA, and protein of interest, respectively. Scale Bar = 200 µm. Representative images out of 3 biological replicates. Statistical analysis was performed via One-way ANOVA with Tukey’s multiple comparisons. *: p ≤ 0.05, **: p ≤ 0.01, ***: p ≤ 0.001, ****: p ≤ 0.0001. Data is presented as (mean ± SEM) from 3 biological replicates, (**J**-**K**), (N=3). Analysis was performed on four technical replicates derived from one biological sample, (**B**-**I**), (n=4).

RNA-seq 5 days after MSC addition showed large differences in gene regulation following regeneration of the DP-treated AML niches with LAM-MSCs or TCP-MSCs (Figure 7B-C). To distinguish transcriptional responses associated with chemotherapy from those associated with subsequent MSC-mediated regeneration, significant differentially expressed gene sets from the leukaemia + chemotherapy, leukaemia + chemotherapy + LAM-MSC, and leukaemia + chemotherapy + TCP-MSC conditions were compared. This analysis revealed both shared and condition-specific transcriptional responses across the three post-leukaemic conditions (Figure 7D).

Focussing on key areas shows that a more pro-regenerative environment is created with the addition of the MSCs, especially with the LAM-MSCs (Figure 7E, *SPP1, BMP2, IGF2, FAM20A, IBSP, SLC12A1, MEDAG*)^62–64^. For all the other focus areas, when therapeutic MSCs are added, there is a positive response compared to not adding MSCs, but the response is more attenuated with the addition of LAM-MSCs compared to TCP-MSCs. The highlighted pathways are: the TNF superfamily, involved in cell communication, regulating immune responses, stromal activation, differentiation, and inflammation (Figure 7F, *TNFAIP6, TNFRSF11B, C1QTNF1, TNFSF15, TNFSF12, TNFRSF18, TNFRSF1B*)^65^; genes involved in disease progression (Figure 7G, *IL-1B, IL-6, IL-11, IL-1A, IL-33, IL-36B, IL-6R, IL24*)^66–68^; genes related to pro-inflammatory chemokine signals used by MSCs to recruit immune cells (Figure 7H, *CXCL6, CXCL5, CXCL3, CXCL8, CXCL1, CXCL2, CXCL12, CXCL16*)^69^, and; genes related to cell survival in adverse environments (Figure 7I, *MT1G, MT1F, MT1M, MT1H, SOD3, SLC7A11, AKR1C2, AKR1C1*)^70^. (Please see table S3 for a list of genes, their full name and their function).

Taken together, these changes illustrate that the addition of therapeutic MSCs to the treated AML niches has broad effects in the areas of regeneration and immune response. This is sensible as MSCs are known for both their regenerative and immunomodulatory potentials. It was seen that the LAM-MSCs appeared to be more pro-regenerative and, while activating immunomodulatory pathways, did so to a lesser extent than the TCP-MSCs.

Next, the response of the MSCs to HSCT in the treated niches was studied. To achieve this, the treated AML niches were removed, using the transwell system, into a fresh well containing CD34^+ve^ cells isolated from BM; this procedure was used so that we could easily isolate the CD34^+ve^ cells for flow and colony analysis as described in the next section, along with either no MSC, control (TCP-MSC) or therapeutic (LAM-MSC) co-therapies (after 21 days of culture) that were injected into the hydrogel; the niches were then cultured for a further 5 days before analysis.

Looking at the MSC spheroids, the addition of the MSCs, especially of the LAM-MSCs, reduced expression of S100A8/9, associated with stromal cancer-associated fibroblasts^71^ (Figure 7J-K). While expression of S100A8/9 is not normally attributed to MSCs, they do express it under stress conditions such as chemotherapy, and its expression in MSCs is linked to AML progression^72^. This further suggests a regenerative effect on the resident MSC population.

Analysing the whole MSC population, GSEA analysis revealed that in the absence of MSC regeneration, adipogenesis and peroxisome pathways were modestly upregulated, indicating a shift toward an adipogenic and stress-adaptive stromal phenotype linked to reduced haematopoietic support. Regeneration with MSCs suppressed adipogenic signalling and reduced peroxisome pathway activity, suggesting restoration of stromal identity and improved redox balance (Figure S12). Lastly, heatmap and PCA analyses (**Figure S13A**-B) demonstrated distinct transcriptomic signatures between control, leukaemic, chemotherapy-treated, and regenerated niches. PCA confirmed strong clustering differences across experimental groups, clearly separating leukaemic from control and therapy-regenerated niches, and distinguishing LAM-MSCs and TCP-MSCs therapies at the gene expression level.

### Characterisation of HSC fate in treated leukaemic niches with MSC co-therapy

The experimental workflow for HSCT is outlined in Figure 8A. AML niches were DP-treated and moved to fresh transwells without THP-1s in order to represent remission. To represent HSC transplantation, CD34^+ve^ HSCs from human BM (50,000 HSCs/mL per niche) were added to the medium in the lower chamber of the transwell, outside the insert containing the hydrogel, to study HSC retention and fate in the remission niches with ease of access to these cells for flow and colony analysis. However, to see if HSC support can be improved by incorporation of pre-conditioned, therapeutic MSCs, TCP-MSCs or LAM-MSCs were introduced, as above, at a concentration of 50,000 cells/mL into the spheroid containing hydrogels. The HSCs were then cultured for 5 days before flow analysis (gating shown in Figure S14).

**Figure 8.**
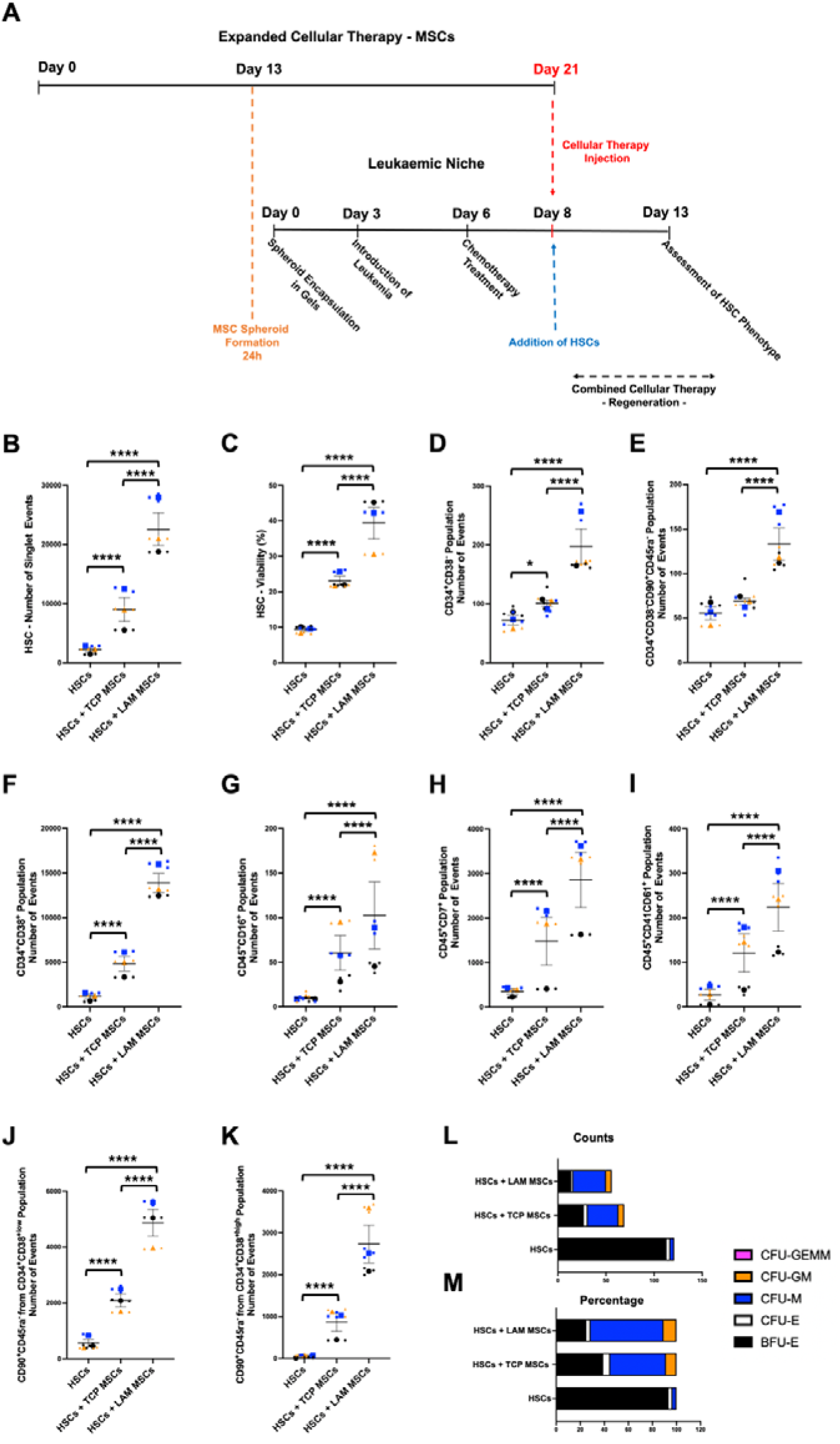
Application of combined cellular therapy in our niche model. Schematic of experimental design timeline (**A**). Quantification of total cell numbers (**B**) and viability (**C**) of HSCs. Quantification of total cell numbers of CD34^+^CD38^-^ (**D**) and CD34^+^CD38^-^ CD90^+^CD45ra^-^ (**E**) HSCs. Quantification of total cell numbers of CD34^+^CD38^+^ (**F**), CD45^+^CD16^+^ (**G**), CD45^+^CD7^+^ (**H**), and CD45^+^CD41CD61^+^ (**I**) cells. Quantification of total cell numbers of CD90^+^CD45ra^-^ cells from the CD34^+^CD38^+low^ (**J**) and the CD34^+^CD38^+high^ (**K**) populations. Quantification of total colony counts by subtype, including BFU-E, CFU-E, CFU-M, CFU-GM, and CFU-GEMM (**L**). Percentage distribution of colony subtypes (**M**). The different coloured symbols on super plots represent the mean of every condition for every experiment. Smaller symbols represent the technical replicates of every experiment, accordingly. Statistical analysis was performed via One-way ANOVA with Tukey’s multiple comparisons. *: p ≤ 0.05, **: p ≤ 0.01, ***: p ≤ 0.001, ****: p ≤ 0.0001. Data is presented as (mean ± SEM) from 3 biological replicates, (N=3).

Data showed significantly more CD34^+ve^ haematopoietic cells (Figure 8B), which were more viable (Figure 8C, **Figure S15A**), after exposure to the treated niches with MSCs added, especially the LAM-MSCs. The same followed when identifying multipotent haematopoietic progenitor cells (CD34^+ve^, CD38^-ve^, Figure 8D, **Figure S15B**)^14,73^ and stem cells (CD34^+ve^, CD38^-ve^, CD90⁺^ve^, CD45ra^-ve^, Figure 8E, **Figure S15C**)^14,73^. It is notable that for these crucial populations, while HSCs cultured with TCP-MSCs showed a trend of more maintained HSCs, only HSCs cultured with LAM-MSCs gave significant retention.

Looking for HSC progeny, flow analysis indicated increased expansion of committed HSC oligopotent progenitors (CD34⁺^ve^, CD38⁺^ve^)^73,74^ and natural killer (CD45⁺^ve^, CD16⁺^ve^)^75^, T-cell (CD45⁺^ve^, CD7⁺^ve^)^76^ and megakaryocytes (CD45⁺^ve^, CD41⁺^ve^, CD61⁺^ve^)^77^ for HSCs introduced to treated niches with MSCs (Figure 8F-I) respectively, (**Figure S15B**, **Figure S16A**-C). LAM-MSCs consistently supported the largest number of HSC progeny, while TCP-MSCs showed a lesser but still significant effect. Further stratification of CD34⁺^ve^, CD38⁺^ve^ cells into low (less committed) and high (more committed) subsets revealed that the CD34⁺CD38⁺low population was enriched in CD90⁺CD45ra^-^ cells, especially in LAM co-cultures (Figure 8J-K, **Figure S17A**-B). This points to the presence of intermediate differentiation states^78^ and suggests that MSCs (especially LAM-MSCs) support expansion without depleting progenitor or HSC pools.

We also used long-term culture and colony assays (long-term culture-initiating cell assays) to look at stem cells and differentiation potential. After 5 days of culture in the treated niches with/without therapeutic MSCs, haematopoietic cells were put into long-term culture (5 weeks) to remove non-stem cells; the resulting HSCs were then allowed to differentiate in a colony formation assay (Figure S18). While at first look, it appears more colonies were observed with HSCs alone, these colonies suffered massive lineage bias, producing almost exclusively erythroid burst-forming units (BFU-E, very early erythroid commitment) (Figure 8L-M). With either LAM-MSCs or TCP-MSCs, however, a good spread of phenotype was seen, demonstrating greater multipotency of the remaining stem cells (Figure 8L-M).

Taken together, the data show increased haematopoietic survival and retention in the treated niches with the addition of therapeutic MSCs. If the MSCs are cultured on PEA-LAM rather than TCP prior to administration, they support greater retention of HSCs and their progenitors, which have greater multipotentiality, and also a more diverse blood cell environment.

## Discussion

We developed a model of the BM where the addition of AML-derived cells (THP-1) induces IL-6, pro-collagen Iα1, fibronectin and MMP-2 expression from spheroid-cultivated MSCs within the model (Figure 5F-M), signatures of remodelling/fibrotic environments that have been linked to blood malignancy^79^. These changes go hand in hand with changes in MSC gene expression profiles attributed to immune response, cell stress and AML survival^47,50,51^ (Figure 5N-Q). Further, the morphology of the MSC spheroids within the BM model was seen to adopt a more activated, disseminated phenotype (Figure 5D). These data illustrate the MSCs remodelling the niche in response to the THP-1 cells; an attribute that we wanted to mimic in our model.

When chemotherapy is applied to the niches, the MSC gene expression patterns show a switch from remodelling to protecting (Figure 6). Genes associated with ECM remodelling, cancer-associated fibroblasts and tumour microenvironment are down-regulated^54,56,58^ in the MSC population. At the same time, THP-1 survival is increased when cultured with the MSCs (Figure 6), demonstrating niche protection^59^, another key feature of mimicking the BM cancer microenvironment.

Achieving these features in our model allows us to then look at HSC support and retention with and without the addition of therapeutic MSCs. As discussed, MSC therapies require a large-scale expansion of the cells *in vitro*^17^. However, as they grow, they suffer phenotypical drift away from their most primitive phenotype and can senesce, along with a shift from a positive, immunomodulatory secretory phenotype to a negative, inflammatory, secretory phenotype^13,15^. Therefore, for optimal cell therapy, the phenotype of the cultured MSCs is of central importance.

In order to achieve an optimal MSC phenotype, we employed PEA, a polymer that can organise ECM proteins (Figure 1). While a reduction in MSC intracellular tension has been linked to preserved multipotency^22^ and immunomodulatory phenotype^13^, this can be challenging to achieve without compromising cell adhesion; for MSCs, well-adhered cells are fundamental for cell growth. LAM has been associated with allowing high levels of cell attachment through integrin ligation, but reducing intracellular tension by reduced association with the microfilament cytoskeleton^24^. Therefore, PEA-LAM was employed here to generate well-adhered MSCs with lowered senescence phenotype (seen with reduction in p16, p53 and IL-8 expression, Figure 2B, C, H) and lowered culture-associated DNA damage (seen by increased TERT expression and reduction in damage-associated factors γH2AX and ATM^15,28^, Figure 2D, F, G). This change in phenotype compared to standard culture was associated with increased production of L-glutamate, glycine, GSH and GSSG; components of one of the cells’ main antioxidant defence pathways^31^ (Figure 2K-P). Indeed, data linked these observations with lowered ROS levels in the MSCs (Figure 2E), ROS being a key driver of cell ageing^80^. Taken together, our BM cancer/treatment model and our MSC with enhanced phenotype (fitness) allow us to next study therapeutic MSCs, which may improve HSC support and retention in a human cell-containing, disease remodelling-associated, post-treatment, 3D microphysiological system.

Addition of MSCs, pre-cultured on either control TCP or pPEA-LAM, into the chemotherapy-treated leukaemic remodelled niches, results in a transcriptional signature broadly related to tissue remodelling with active stromal-immune crosstalk and metabolic adaptation^81^ (Figure 7E-I). This is sensible given that the therapeutic MSCs were interacting with MSCs that had been cultured with diseased immune cells, had radically changed metabolism, and had remodelled the niche for cancer (Figure 5) before exposure to chemotherapeutic drugs (Figure 6). As HSCs are added with/without MSCs, reduced expression of S100A8/9, associated with stromal support of cancer cells^72^, is seen for MSCs in the model spheroids (Figure 7J, K); this is a highly significant reduction when LAM-MSCs are added compared to the addition of no MSCs or TCP-MSCs.

Looking at the HSC population added to the post-leukaemic niche, addition of therapeutic MSCs led to higher numbers of HSC-derived cells in the model, and this included higher numbers of primitive cells, progenitor cells and mature progeny (Figure 8B-K). Looking more specifically at long-term HSC function using LTC-IC, it appears that longer-term cells (which should be the most primitive) removed from the HSC support experiment without therapeutic MSCs resulted in cells with very little differentiation potential (they all form BFU-E). This is clearly undesirable as the aim is for HSCs to engraft and then to be able to repopulate the whole blood system. However, with therapeutic MSCs, especially LAM-MSCs, broader multilineage colony-forming potential in the long-term cells was observed (Figure 8L, M). This result suggests poor niche support for HSCs cultured without therapeutic MSCs. Indeed, HSCs require CXCL12, SCF, TGF□β, Notch, and angiocrine signals as well as a low ROS, low inflammation environment to maintain multipotentiality; damaged niches may allow survival, but not function, and stressed niches result in preferential myeloid bias^37^. The therapeutic MSCs, and critically, the LAM-MSCs, support maintenance of multipotent HSCs, which would be more therapeutically useful.

This study has limitations. The model uses THP-1 cells as a tractable representation of AML and therefore does not reproduce the full biological heterogeneity of patient disease or every cellular and vascular component of the human bone marrow niche. In addition, the RNA-sequencing and stable-isotope tracing analyses used technical replicates derived from a single biological sample, limiting assessment of biological variability. Future studies should validate the findings using primary AML material and additional biological replicates and incorporate further niche components where appropriate.

## Methods

### Substrate Preparation

Tissue culture treated plates (TCP) were immobilised in a plasma chamber vertically to the flow, while coverslips were placed on a flat surface horizontally. To completely remove any residual organic matter, the samples were first treated with air plasma for 5 minutes at 50 W of radio frequency (RF) incident power. Consequently, for the polymerisation of the PEA monomer on the surfaces (pPEA), RF was set at 50 W for 15 minutes. Before cell culture use, the PEA-coated samples were sterilised for 30 minutes under UV light. Regarding the adsorption of FN and LAM from the PEA surfaces, human FN and murine LAM (stocks 1 mg/mL) were diluted in PBS to a working concentration of 20 μg/mL. FN or LAM were added on top of the PEA-coated samples for 1 hour, ensuring even coverage of the surfaces. After the incubation, surfaces were washed 2 times and kept in PBS until cell seeding. For every experiment, the corresponding TCP without PEA coating were used as a control.

### X-ray photoelectron spectroscopy

X-ray photoelectron spectroscopy (XPS) was employed to determine the surface chemical composition of the samples. All XPS spectra were acquired at the Harwell XPS facility using a K-alpha XPS system (Thermo Fisher) equipped with a monochromatic Al K-alpha source.Each sample was analysed three times, with a maximum beam size of 400 μm × 800 μm. The instrument parameters were set as follows: X-ray energy: 1486.68 eV, voltage: 12 kV, current: 3 mA, power: 36 W. Spectral analysis and curve fitting were conducted using CasaXPS software to ensure accurate data interpretation.

### Contact Angle Measurements

Contact angle measurements were performed to evaluate the wettability of Nunc™ Thermanox™ Coverslips (Thermo Fisher, #174950) under the following conditions: TCP, PEA-coated controls, as well as PEA-coated surfaces covered with LAM or FN. Static water contact angles were measured using an Easy Drop goniometer (Kruss DSA20E). A fixed volume (5 µL) of deionised water was dispensed onto the surface using an automated syringe. Images of the droplet profile were captured, and the contact angle was determined using the DSA1 software. Surface wettability was assessed by comparing contact angles, with lower angles indicating greater hydrophilicity while higher angles greater hydrophobicity. All measurements were conducted at room temperature (RT).

### Micro BCA Assay

The Micro BCA^TM^ Protein Assay Kit (Thermo Fisher, #23235) was used to quantify the amount of FN and LAM adsorbed onto TCP and PEA substrates, following the manufacturer’s instructions. To facilitate protein adsorption, 20 μg/mL of FN or LAM was added to the substrate surfaces and incubated for 1 hour at RT. After incubation, the supernatant was carefully removed and collected in Protein LoBind Tubes (Eppendorf) for further analysis. All samples, including standards, were loaded in triplicate onto a 96-well plate, followed by the addition of the Micro BCA working reagent. The plate was incubated for 30 minutes at RT to allow colour development. Absorbance was measured at 562 nm using a Multiskan FC microplate reader.

### LAM PEGylation

Murine laminin-111 (LAM-111) (Thermo Fisher, 1 mg/mL, #23017015) was PEGylated using PEG-MAL-SVA (5 kDa, Laysan Bio) to covalently modify the protein with PEG groups. PEGylation was performed based on the molecular weight of LAM-111 (850 kDa) at a 1:4 molar ratio of LAM to PEG-MAL-SVA. LAM-111 was initially supplied in 50 mM Tris-HCl (pH 7.4), 0.15 M NaCl at 1 mg/mL. Prior to PEGylation, the required volume was dialysed into sodium bicarbonate buffer (NaHCO₃, pH 8.5), as PEGylation occurs optimally at this pH. PEG-MAL-SVA (1 mg/mL in NaHCO₃) was then added at the calculated volume to achieve the desired molar ratio and allowed to react for 2 hours at RT under mixing. Following PEGylation, the solution was dialysed to remove unreacted PEG and acrylates, while exchanging the buffer to PBS for 1 hour at 4 °C. The PEGylated LAM-111 was stored at -20 °C for future use.

### Gel preparation

After the overnight development of MSCs to spheroids, harvesting was followed with PBS supplemented with 2% FBS to maximise the yield of collected spheroids. The PEG -4 maleimide (PEG-4MAL) (20K, Laysan Bio) and the crosslinker PEG-Dithiol (PEG-SH) (2K, Creative PEGworks), were sterilised with UV-light for 30 minutes followed by dilution in PBS to 500 mg/mL and 200 mg/mL, respectively. The degradable crosslinker VPM (GenScript) was diluted in PBS to 200 mg/mL and mixed with PEG-SH at a molar ratio of 1:3. The PEG-4MAL and crosslinker solutions were mixed at a molar ratio of 1:1 (maleimide: thiol) to be fully crosslinked. 100 spheroids were incorporated in 100 μL of PEG-gel (1 spheroid/1 gel μL). The final concentration of PEG is 10% w/v. After the addition of the crosslinker, samples were incubated for at least 30 minutes at 37 °C to allow gelation.

### Bulk rheological characterisation

Rheological measurements were conducted using a stress-controlled rheometer (MCR302, Anton Paar) with a parallel plate geometry (15 mm upper plate diameter) at 23 °C. To maintain sample hydration, PBS was added to the exposed edges of the specimens. During gel preparation, a 12 mm hydrophobic glass coverslip (treated with RainX solution, RainX) was placed on top of the gels to ensure uniform thickness. Before all experiments, a frequency sweep was performed to identify the range in which the shear storage modulus (G’) remained independent of angular frequency (ω), allowing the shear modulus (G) to be determined as (*G* = lim_ω→0_ *G′ (*ω)). Strain sweeps were conducted within the linear viscoelastic (LVE) regime over a strain range of 0.01% to 1%, at a constant angular frequency of 10 rad/s. Measurements were taken at a gap size corresponding to a normal force of ≈0.1 N for all gels. Additionally, a series of strain sweep tests were performed under varying levels of compression. These tests involved sequentially increasing the normal force applied to the unconfined samples, with each strain sweep corresponding to a different compression level. The normal force started at ≈0.1 N, with minimal delays (on the order of a few seconds) between successive compressions. The Young’s modulus (E) was obtained from the shear modulus (G) via the following equation *E* = 2*G(1 +* υ), where Poisson’s ratio (υ) was assumed to be 0.5.

### Cell Culture

BM-MSCs purchased from PromoCell (#C-12794) and maintained below passage 5 were used for experiments. Cells were cultured in flasks followed by media change every 3 days until the monolayers reached 80% confluency. Cells were detached and seeded on different substrates at 2000 cells/cm^2^ in 10% FBS/DMEM. For the hypoxic experiments, cells were transferred to a hypoxic workstation (Ruskinn) set at 5% O_2_ on the third day of culture. For developing single cell MSCs to spheroids, 24-well AggreWell^TM^400 plates (STEMCELL Technologies, #34415) were used, treated with anti-adherence solution (STEMCELL Technologies, #07010). MSCs were seeded into AggreWell^TM^400 24-well plates at a density of 120,000 cells per well, corresponding to approximately 100 cells per spheroid.

After encapsulating the MSC spheroids within PEG-gels, the constructs were transferred into (Greiner) 12-well transwell inserts with 3.0 µm porous membranes. The transwells were maintained in DMEM for 3 days to allow initial conditioning. On day 3, the medium in the lower chamber was removed and replaced with THP-1 cells at a concentration of 50,000 cells/mL, suspended in a 1:1 mixture of DMEM and RPMI. The co-culture was maintained for an additional 3 days to facilitate BM niche remodelling. Finally, the established leukaemic niche was treated with daunorubicin and panobinostat for 2 days to induce niche damage.

HSCs (BM CD34^+^) purchased from STEMCELL Technologies (#70002) were used for experiments. After thawing, cells were then allowed to rest overnight in complete IMDM supplemented with BIT and 1% 200 mM L-glutamine, cytokine-enriched media Flt3L (50^ng/mL), SCF (20^ng/mL), and TPO (25^ng/mL) (all recombinant human, Peprotech). The leukaemic niche was established as previously described. Following chemotherapeutic treatment, the PEG-based gel constructs were carefully removed from the transwell inserts using a sterile spatula and transferred into fresh transwell inserts and 12-well plates. The lower chamber of each well was filled with 1.2□mL of HSCs suspended in IMDM + BIT at a concentration of 50,000 cells/mL, resulting in a total of 60,000 HSCs per well. Meanwhile, expanded MSCs intended for cellular therapy were harvested using trypsinisation and injected into the gel constructs at a density of 50,000 cells per gel. This approach maintained an approximate 1:1 MSC-to-HSC ratio, accounting for both the injected therapeutic MSCs and the 10,000 pre-existing MSCs within the gel (100 spheroids) that had been remodelled during leukaemic niche formation. All cultures were incubated in a 5% humidified CO_2_ atmosphere at 37 °C.

### Quantitative real-time PCR

After 21 days of culture, MSCs were harvested via the TRIzol™ Reagent (Thermo Fisher, #15596026). Samples were incubated for 5 minutes before addition of chloroform (200 μl chloroform/1 ml Trizol). Samples were incubated for 3 minutes while a 15-minute centrifugation at 12,000 xg at 4°C was followed. The colourless upper aqueous phase was transferred to a new Eppendorf tube followed by addition of isopropanol (500 μl isopropanol/1 ml Trizol) and a 24-hour incubation at -20 °C. The next day, samples were centrifuged at 12,000 g at 4 °C. The supernatant was removed, and pellets were resuspended in 75% ethanol. Samples were vortexed quickly and centrifuged for 5 minutes at 7500 × g at 4 °C. After the supernatant was discarded, the pellets were resuspended in RLT buffer. Further processing with the RNeasy® Micro Kit (Qiagen, #74004) was followed to maximise the purity of RNA samples, following the manufacturer’s instructions. The quantity and purity of RNA samples was assessed with NanoDrop (Thermo Fisher). Consequently, RNA samples were reverse transcribed using the QuantiTect Reverse Transcription Kit (Qiagen, #205311), following the manufacturer’s instructions. qPCR was performed using the QuantiNova® SYBR® Green PCR kit (Qiagen, #208052), following the manufacturer’s instructions, and the 7500 Real-Time PCR System (Applied Biosystems). Relative gene expression was calculated using the 2^-ΔΔCT method and normalised to the housekeeping control gene GAPDH.

### Metabolomics analysis

MSCs cultured on FN, LAM, and TCP for 7 or 14 days were washed with ice-cold PBS and lysed using an extraction buffer composed of distilled H₂O, methanol, and chloroform (1:3:1 ratio). The lysis process was carried out for 1 hour at 4 °C on a shaker. Following lysis, samples were transferred to Eppendorf tubes and centrifuged at 13,000 g for 5 minutes at 4 °C to separate cellular debris from the supernatant. The resulting extracts were stored at -80 °C until further processing at Glasgow Polyomics, following their standardised metabolomics protocols. Each condition was represented by three technical replicates derived from a single biological replicate.

Liquid chromatography-mass spectrometry (LC-MS) analysis was performed using an UltiMate 3000 RSLC system (Thermo Fisher) coupled with an Exactive (Orbitrap) mass spectrometer (Thermo Fisher). Chromatographic separation was achieved using a 150 × 4.6 mm ZIC-pHILIC column, operating at a flow rate of 300 μL/min. Raw mass spectrometry data were processed using a combination of bioinformatics tools: XCMS for peak picking, mzMatch for data filtering and grouping, and IDEOM for further filtering, post-processing, and identification. To validate metabolite identification, a panel of known standards was used, matching metabolites by accurate mass and retention time. Further metabolite identifications were inferred based on these validated standards. Heatmaps and principal component analysis (PCA) plots were generated using MetaboAnalyst software (version 6.0) for visualisation and statistical analysis.

### 13C6-Glucose metabolomic tracing

MSCs were cultured on LAM, FN, and TCP, and allowed to grow for 11 days. Each condition was represented by four technical replicates derived from a single biological replicate. Cells were washed and incubated in basal media comprising 25% normal glucose and 75% ^13^C - Glucose (Cambridge Isotopes Ltd) for a further 3 days. Extractions and LC-MS were performed as previously described. The LC-MS platform consisted of an Accela 600 HPLC system (Thermo Fisher) combined with an Exactive (Orbitrap) mass spectrometer. Two complementary columns were used: the zwitterionic ZIC-pHILLIC column (150□mm□×□4.6□mm; 3.5 μm, Merck) and the reversed phase ACE C18-AR column (150□mm□×□4.6□mm; 3.5 μm Hichrom) and in both cases, the sample volume was 10 μl at a flow rate of 300 μL/min. Eluted samples were then analysed by mass spectrometry. LCMS data of ^13^C-labelled extracts were processed to generate a combined PeakML file as described previously. Further analysis using mzMatch-ISO in R66 generated a PDF file containing chromatograms used to check peak shape and retention time. A tab-delineated file detailing peak height for each isotopologue was also generated to calculate percentage labelling. Cellular protein content was estimated using Coomassie Blue staining in quadruplicate and used to normalise metabolite abundance across samples.

### In-Cell Western

For in-cell western (ICW) analysis, the samples were fixed with the fixative solution for 15 minutes at 37 °C. Next, samples were permeabilised with the permeabilisation solution for 5 minutes at 4 °C on a shaker. 5% milk/PBS was used as the blocking solution for 1 hour at 37 °C, followed by a 30-minute incubation at RT on a shaker. The primary antibodies (Table S4) were incubated overnight in 1% milk/PBS at 4 °C on a shaker. The next day, the samples were washed with 0.1% Tween-20/PBS buffer 5 times, followed by incubation with the secondary antibodies IRDye 800CW (LI-COR, #926-32211, #926-32350) and CellTag^TM^ 700 Stain (LI-COR, #926-41090) in 1% milk/PBS covered in tin foil at RT on a shaker for 1 hour. Afterwards, samples were washed with 0.1% Tween-20/PBS buffer 5 times, followed by drying with an aspirator. Samples were scanned with a LI-COR Odyssey M imaging system, where signal detection at the 800 nm channel indicates the expression of the protein of interest, while the 700 nm channel indicates the total protein stain (indicative of cell number). Thereafter, the obtained normalised expression values 800nm/700nm were further normalised against the TCP control.

### Immunostaining

For immunostaining, the samples were fixed with the fixative solution for 15 minutes at 37 °C. Next, samples were permeabilised with the permeabilisation solution for 5 minutes at 4 °C on a shaker. For the staining of TERT, ice-cold methanol was used to fix the samples at -20 °C for 20 minutes. Then, samples were treated with 2M of HCL for 20 minutes at RT on a shaker, followed by a quick wash with 0.1 M of sodium borate (Na₂B₄O₇). Finally, samples were permeabilised for 15 minutes at RT on a shaker. 5% BSA/PBS was used as the blocking solution for 1 hour at RT on a shaker. The primary antibodies (Table S4) were incubated overnight in 1% BSA/PBS at 4 °C on a shaker. The next day, the samples were washed with 0.5% Tween-20/PBS buffer 5 times, followed by incubation with the Texas Red™ secondary antibodies (Vector Laboratories, #TI-2000-1.5, #TI-1000-1.5) and Oregon Green™ 488 Phalloidin (Thermo Fisher, #O7466), covered in tin foil at RT on a shaker for 1 hour. Afterwards, samples were washed with 0.1% Tween-20/PBS buffer 5 times, followed by NucBlue™ Live ReadyProbes™ Reagent (Hoechst/PBS) staining (Thermo Fisher, #R37605) at RT on a shaker for 30 minutes. The staining was removed, and samples were kept in PBS.

Protein expression on spheroids was quantified via the following formula: Expression = (Protein ROI - Protein Background ROI) / (DNA ROI - DNA Background ROI). To assess the viability of spheroids, gels were washed with PBS, followed by staining with LIVE/DEAD Viability Kit (Thermo Fisher, #L3224), with a final concentration of ethidium dimer being 4 µM and of calcein AM 2 µM in PBS, for 20 minutes at 37°C. Spheroid viability was quantified via the following formula: Viability = ((Live MFI) / (Live MFI + Dead MFI)) x 100. Samples were visualised via EVOS M7000 (Invitrogen) or ZEISS Axio Observer Z1 fluorescent microscopes. The spheroid size was calculated via the number of z-projections multiplied by the step size. ImageJ was used to analyse images from fluorescent microscopy.

### ELISA

The DuoSet^®^ ELISA kit (R&D Systems) was used to quantify IL-6 (#DY206), fibronectin (#DY1918), and pro-collagen Iα1 (#DY6220) levels in supernatant of co-cultured MSCs with THP-1s, following the manufacturer’s protocol. 96-well high-binding microplates were coated overnight at 4 °C with capture antibody diluted in PBS. The next day, plates were washed 3 times with wash buffer (0.05% Tween-20/PBS) and blocked with reagent diluent (1% BSA/PBS) for 1 hour at RT. After blocking, samples were prepared in triplicate, while the standards, in duplicate and added to the wells, followed by incubation for 2 hours at RT. Plates were then washed and incubated with the biotinylated detection antibody for 2 hours at RT. After 3 washes with 0.05% Tween-20/PBS, streptavidin HRP was added and incubated for 20 minutes covered in tin foil, at RT. Next, 3 washes with 0.05% Tween-20/PBS were followed, while the reaction was developed using substrate solution (TMB, tetramethylbenzidine) for 20 minutes. Lastly, the reaction was stopped by the addition of stop solution (2N H₂SO₄). Absorbance was measured at 450 nm with a correction wavelength of 570 nm using a Multiskan FC microplate reader.

### Zymography

Conditioned media from MSC spheroids was harvested and processed fresh for electrophoresis. To estimate molecular weight, SeeBlue™ Pre-stained Protein Standard (Thermo Fisher, #LC5625) was loaded alongside the samples. Samples were mixed with Tris-Glycine SDS Sample Buffer (2X) (Thermo Fisher, #LC2676) before loading on Novex™ Zymogram Gels (Thermo Fisher, #ZY00100BOX), run for 90 minutes at 125 V with 1X Novex™ Tris-Glycine SDS Running Buffer (Thermo Fisher, #LC2675). After electrophoresis, the gel was removed and incubated in 1X Novex™ Zymogram Renaturing Buffer (Thermo Fisher, #LC2670) for 30 minutes at RT on a shaker. After discarding the renaturing buffer, 1X Novex™ Zymogram Developing Buffer (Thermo Fisher, #LC2671) was added on top of the gel, which was left to incubate overnight at 37 °C. The next day, gels were stained in 0.5% Coomassie Brilliant Blue R-250 (Sigma, #1.12553) in 4% methanol, 10% acetic acid at RT for 1 hour on a shaker. The Coomassie Blue solution was replaced with destain solution (4% methanol and 10% acetic acid in water), which was replaced until bands were visible. Gels were scanned with a LI-COR Odyssey M imaging system. ImageJ was used to analyse images from zymography analysis. Band intensities were quantified after background correction, and values obtained from the media-only control were subtracted from the conditioned media samples to account for baseline proteolytic activity present in the culture media.

### Proliferation

AlamarBlue (BIO-RAD) was used to evaluate the metabolic activity of leukaemic cells. THP-1s were seeded into T-25 cm^2^ flasks and incubated under standard culture conditions.

Following treatment with panobinostat, daunorubicin, and cytarabine for 2 or 4 days under various concentrations, the alamarBlue reagent (resazurin) was added to each well at a final concentration of 10% (v/v) and incubated for 6 hours at 37 °C, 5% CO₂. After incubation, the absorbance was measured using a Multiskan FC microplate reader at 570 nm and 600 nm, corresponding to the absorbance maxima of reduced resorufin and oxidised resazurin, respectively. The percentage reduction of alamarBlue was calculated using the following equation:

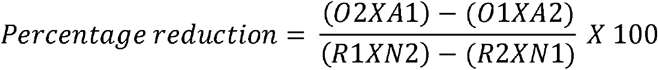

### Spheroid Proliferation

To assess the metabolic activity of spheroids in the PEG-gels, CellTiter-Glo^®^ 3D (Promega, #G9681) was used. Gels with spheroids grown for 2 or 8 days were washed with PBS, followed by the addition of the reagent and left for 5 minutes to incubate on a shaker at RT, covered with foil. Afterwards, the gels were removed from the shaker and left to incubate for another 25 minutes at RT before recording the chemiluminescence on a Modulus^TM^ II microplate reader.

### Flow cytometry

MSCs were washed with PBS and detached using TrypLE Express (Thermo Fisher) to minimise disruption of surface markers and preserve cell integrity. The detached cells were subsequently supplemented with 10% FBS/DMEM, followed by harvesting and spinning at 300 g for 4 minutes. MSC or HSC cell suspensions were split in Eppendorf tubes and spun down at 300 g for 5 minutes, while 3 pooled samples from all the replicates were created. For the viability assessments, a pooled sample was fixed with fixative solution for 30 minutes at 37 °C to identify the fluorescence of dead cells, while a second one was kept as a control for the live cells. The Zombie Violet™ Fixable Viability Kit (BioLegend, #423113) was diluted in PBS, and 100 μl were used to stain each cell pellet for 30 minutes at RT. Next, the samples were centrifuged, followed by resuspension with 100 μl of antibody master mix diluted in flow buffer for 1 hour at 4 °C (Table S5). Samples were spun down, and the antibody solution was discarded, followed by resuspension in 400 μl of flow buffer before flow cytometry analysis. The last pooled sample remained unstained as a negative control to determine the autofluorescence level of the cells and indicate the gating strategy.

To compensate for the fluorescence spillover to other channels, 1 μl of every fluorophore was added in 1 drop of the UltraComp eBeads^TM^ (Thermo Fisher, #01-2222-41) accordingly. Similarly, to the samples, the tubes with the compensation beads were incubated for 1 hour at 4 °C. After centrifugation, the supernatant was discarded and replaced with 400 μl of flow buffer. Regarding the staining of MSCs for ROS, the Cellular ROS Assay Kit (Red) (Abcam, #ab186027) was used to stain each pellet with 100 μl for 1 hour at 37 °C. After centrifugation, the supernatant was discarded and replaced with 400 μl of flow buffer. Similarly, unstained samples were used to determine autofluorescence levels. Cell populations were quantified using the number of gated cell-associated events recorded during flow cytometry analysis. Data analysis was performed with the FCS Express 7 software.

### RNA-sequencing and analysis

Total RNA was extracted from control, leukaemic, leukaemic + chemotherapy, leukaemic + chemotherapy + LAM-MSCs, and leukaemic + chemotherapy TCP-MSCs, as previously described. Each condition was represented by four technical replicates derived from a single biological replicate. RNA sequencing (RNA-Seq) was performed by Azenta Life Sciences following their standardised protocols. RNA quality and integrity were assessed using the Agilent 2100 Bioanalyzer System. mRNA sequencing was conducted using poly(A) selection, and library preparation was followed by sequencing on an Illumina platform with a paired- end (PE) 2×150 strategy, generating 150 bp paired-end reads, with an average sequencing depth of 20 million reads per sample. To ensure data quality, sequence reads were trimmed to remove adapter sequences and low-quality nucleotides using Trimmomatic v0.36. The cleaned reads were then aligned to the Homo sapiens GRCh38 reference genome (ENSEMBL) using the STAR aligner v2.5.2b. Unique gene hit counts were quantified using featureCounts from the Subread package v1.5.2.

Following count extraction, differential gene expression (DEG) analysis was performed using DESeq2, identifying significantly upregulated and downregulated genes. Log_2_ fold change values from significant DEGs were utilised to generate heatmaps of genes of interest. Additionally, this data was analysed using Ingenuity Pathway Analysis (IPA, Qiagen). To identify shared and condition-specific transcriptional responses, significant DEG lists from each comparison were cross-compared and visualised using Venn diagrams. Principal Component Analysis (PCA) was performed using iDEP (integrated Differential Expression and Pathway analysis) to explore sample clustering based on gene expression profiles. Transcripts Per Million (TPM) values were log-transformed (log(*TPM* + 1)) before PCA to stabilise variance and reduce the influence of highly expressed genes. Gene set enrichment analysis (GSEA) was performed using the GSEA software (version 4.4.0).

Gene nomenclature and broad functional information for genes highlighted in the transcriptomic analyses were cross-checked using the NCBI Gene database (National Center for Biotechnology Information, U.S. National Library of Medicine). Functional annotations were subsequently curated according to the biological processes and disease contexts examined in this study.

### Long-Term Culture-Initiating Cell (LTC-IC) Assay

LTC-IC assay was performed using CD34⁺ HSCs isolated from leukaemic niche conditions after 5 days of culture. Mouse stromal fibroblast feeder layers were established using two cell lines: M2-10B4 (overexpressing IL-3 and G-CSF), maintained in RPMI with 10% FBS, and Sl/Sl (overexpressing IL-3 and SCF), maintained in DMEM with 10% FBS. Both cell types were cultured for 1 week until reaching ∼80% confluence, then seeded at a 1:1 ratio into collagen-coated 6-well plates (StemCell Technologies, UK). Once the feeder layer reached ∼80% confluence, cells were mitotically inactivated using mitomycin C (20 μg/mL). Subsequently, CD34⁺ HSCs were seeded onto the feeder layers in hydrocortisone-supplemented HLTM medium, according to the manufacturer’s instructions (StemCell Technologies, UK).

### CFU Assay

After 5 weeks of LTC-IC culture, cells were harvested for CFU assessment. HLTM medium containing non-adherent cells was gently resuspended and transferred into a 15 mL Falcon tube. The suspension was diluted with IMDM supplemented with 2% FBS and centrifuged at 300 × g for 8 minutes. Following centrifugation, the supernatant was carefully aspirated, leaving approximately 100 µL of medium to avoid disturbing the cell pellet. Cells were counted, and 3,000 cells were resuspended in 3.3 mL of MethoCult medium (StemCell Technologies) in Bijou tubes. Tubes were vortexed briefly and left at RT for 5 minutes to allow any air bubbles to dissipate. Each replicate was plated in duplicate into non-tissue culture-treated 6-well plates using a 1 mL syringe (without a needle) to ensure even distribution. Sterile PBS was added between the wells to minimise MethoCult evaporation during incubation. Plates were incubated at 37 °C in 5% CO₂ for 14 days. Colony formation was assessed using a light microscope, and colony types were identified and counted. Two independent researchers performed the counts, and the results were averaged to ensure consistency and accuracy.

### Statistical Analysis

GraphPad Prism 8^TM^ software was used to perform statistical analysis. The Shapiro-Wilk test was used to determine if the data is normally distributed. For n=2 groups, Student’s t-test was used, while for n > 2 groups, One-way ANOVA with Tukey’s multiple comparisons test was used to identify significance accordingly. Considering data visualisation, super plots were used to display individual technical replicate measurements alongside the mean of each biological replicate. This approach enhances transparency by visualising both within-experiment and between-experiment variability.

## Resource availability

### Lead contact

Requests for further information and resources should be directed to and will be fulfilled by the lead contact.

## Materials availability

This study did not generate any new, unique reagents.

## Data and code availability

All raw data are available through the University of Glasgow Enlighten Repository at https://doi.org/10.5525/gla.researchdata.2343. Materials used in this study are described in Methods and are commercially available unless otherwise stated. No custom code is required to reproduce the analyses reported.

## Acknowledgements

The authors thank Dr. Vineetha Jayawarna for assistance with XPS and characterisation of PEA coatings with fibronectin and laminin. I.A.T. was supported by a Medical Research Council (MRC) iCASE studentship (MR/R01566X/1). This work was supported by the Engineering and Physical Sciences Research Council (EPSRC) Programme Grant StemNiche (EP/X036049/1; M.J.D. and M.S.S.), the EPSRC Hub MAINSTREAM (APP18790; M.J.D. and M.S.S.), and the European Research Council (ERC) Advanced Grant DEVISE (101054728; M.S.S.). IBEC is a recipient of a Severo Ochoa Centre of Excellence award from MICINN and is a member of the CERCA Programme of the Generalitat de Catalunya.

## Author contributions

Conceptualization: I.A.T., Y.A., M.J.D., P.M.T., M.V.; Methodology: I.A.T., M.S.S.; Investigation: I.A.T.; Formal analysis: I.A.T., M.J.D., P.M.T.; Funding acquisition: M.J.D., M.S.S.; Writing - original draft: I.A.T., M.J.D.; Writing - review & editing: I.A.T., Y.A., P.M.T., M.V., M.S.S., M.J.D.

## Supplemental information

Document S1. Figures S1–S18 and Tables S1–S5

---

**Figure S1.**
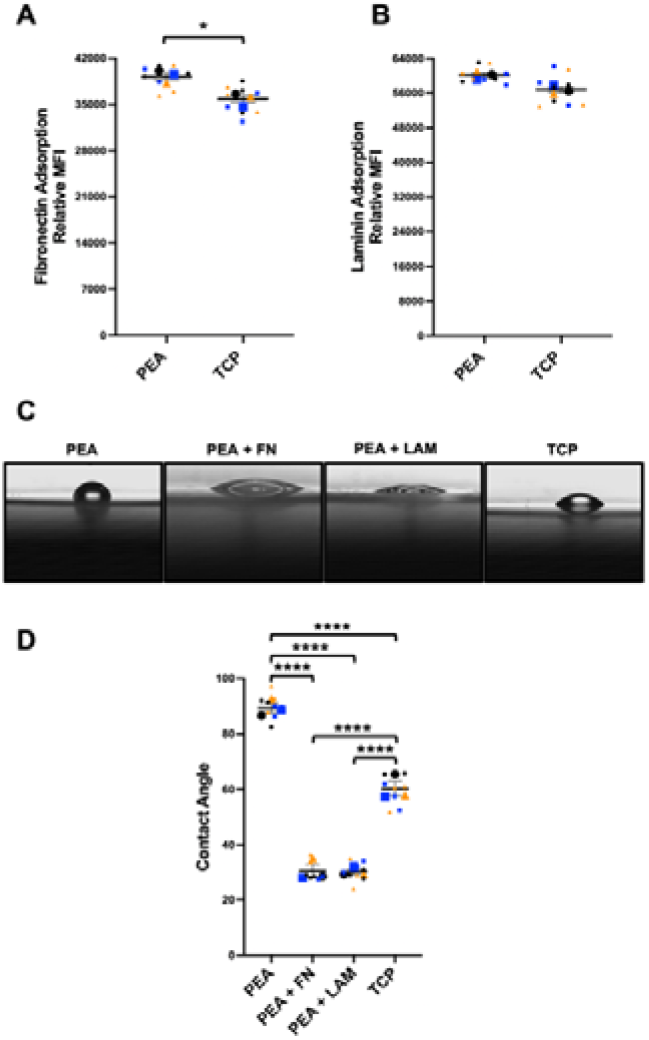
Characterisation of PEA-coated surfaces and contact angle measurements to show successful coating. Quantification of fluorescence intensity of total FN (**A**) and LAM (**B**) adsorption. Quantification of contact angle. PEA surfaces were more hydrophobic (>80°) than TCP (∼60°; p < 0.0001). Adsorption of FN or LAM reduced hydrophobicity significantly (∼30°; p < 0.0001 vs. PEA and TCP) (**C**-**D**). The different coloured symbols on graphs represent the mean of every condition for every experiment. Smaller symbols represent the technical replicates of every experiment, accordingly. Statistical analysis was performed via Student’s t-test or One-way ANOVA with Tukey’s multiple comparisons. *: p ≤ 0.05, **: p ≤ 0.01, ***: p ≤ 0.001, ****: p ≤ 0.0001. Data is presented as (mean ± SEM) from 3 biological replicates, (N=3).

**Figure S2.**
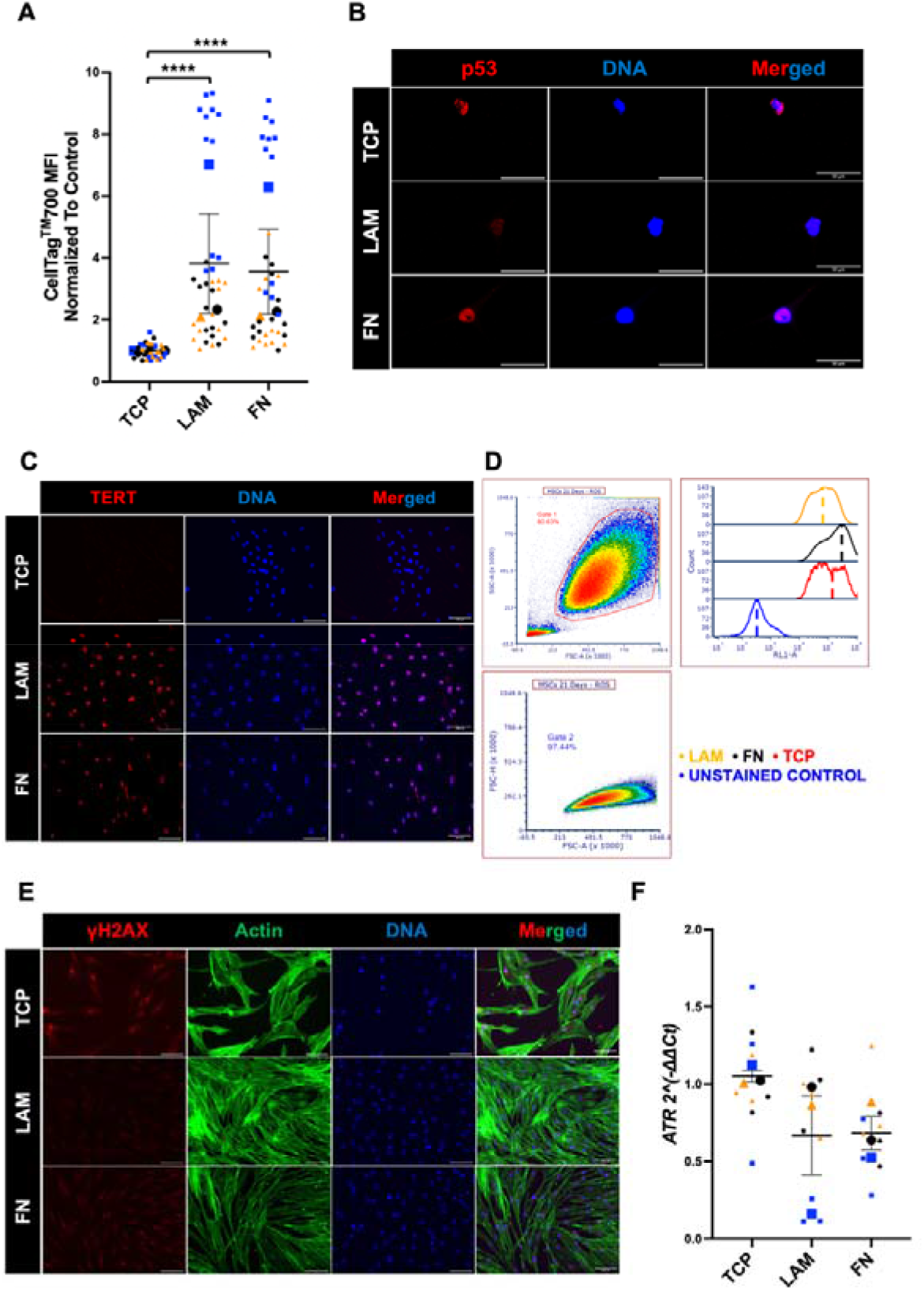
Investigating the effects of LAM, FN, and TCP in MSC senescence. Quantification of total protein stain CellTag^TM^700 of MSCs (**A**). MSC staining for p53 (**B**) and TERT (**C**). Gating strategy and MFI histograms for ROS. Histograms indicate changes in fluorescence intensity in ROS production. A right shift of the curve indicates an upregulation in ROS production. Dashed lines signify the middle of the curve. Black curves correspond to FN while red and orange to TCP and LAM, accordingly. Blue curve refers to unstained control (**D**). MSC staining for γH2AX (**E**). Gene expression of ATR (**F**). The different coloured symbols on super plots represent the mean of every condition for every experiment. Smaller symbols represent the technical replicates of every experiment, accordingly. Blue, green, and red stains represent the DNA, F-actin, and protein of interest, respectively. Scale Bar = 100 µm. Representative images out of 3 biological replicates. Statistical analysis was performed via One-way ANOVA with Tukey’s multiple comparisons. *: p ≤ 0.05, **: p ≤ 0.01, ***: p ≤ 0.001, ****: p ≤ 0.0001. Data is presented as (mean ± SEM) from 3 biological replicates, (N=3).

**Figure S3.**
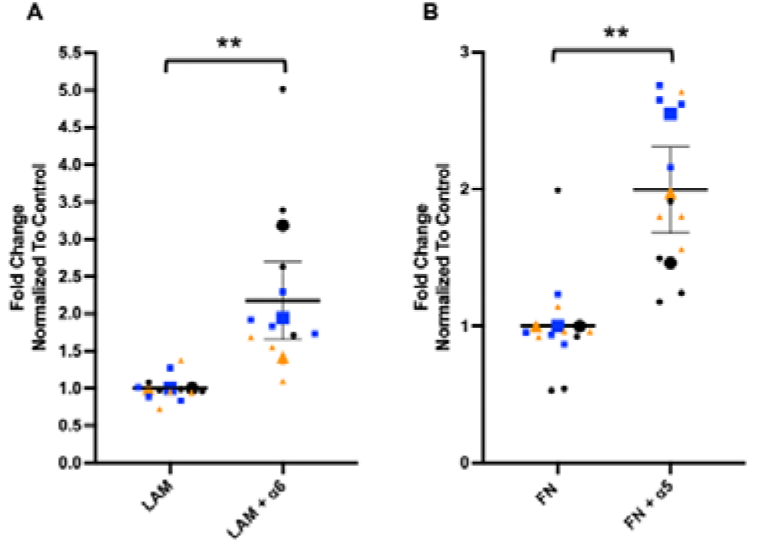
Assessment of α6 and α5 roles in the regulation of senescence at day 21. Expression of p53 on LAM (**A**) and FN (**B**). The different coloured symbols on super plots represent the mean of every condition for every experiment. Smaller symbols represent the technical replicates of every experiment, accordingly. Statistical analysis was performed via Student’s t-test. *: p ≤ 0.05, **: p ≤ 0.01, ***: p ≤ 0.001, ****: p ≤ 0.0001. Data is presented as (mean ± SEM) from 3 biological replicates, (N=3).

**Figure S4.**
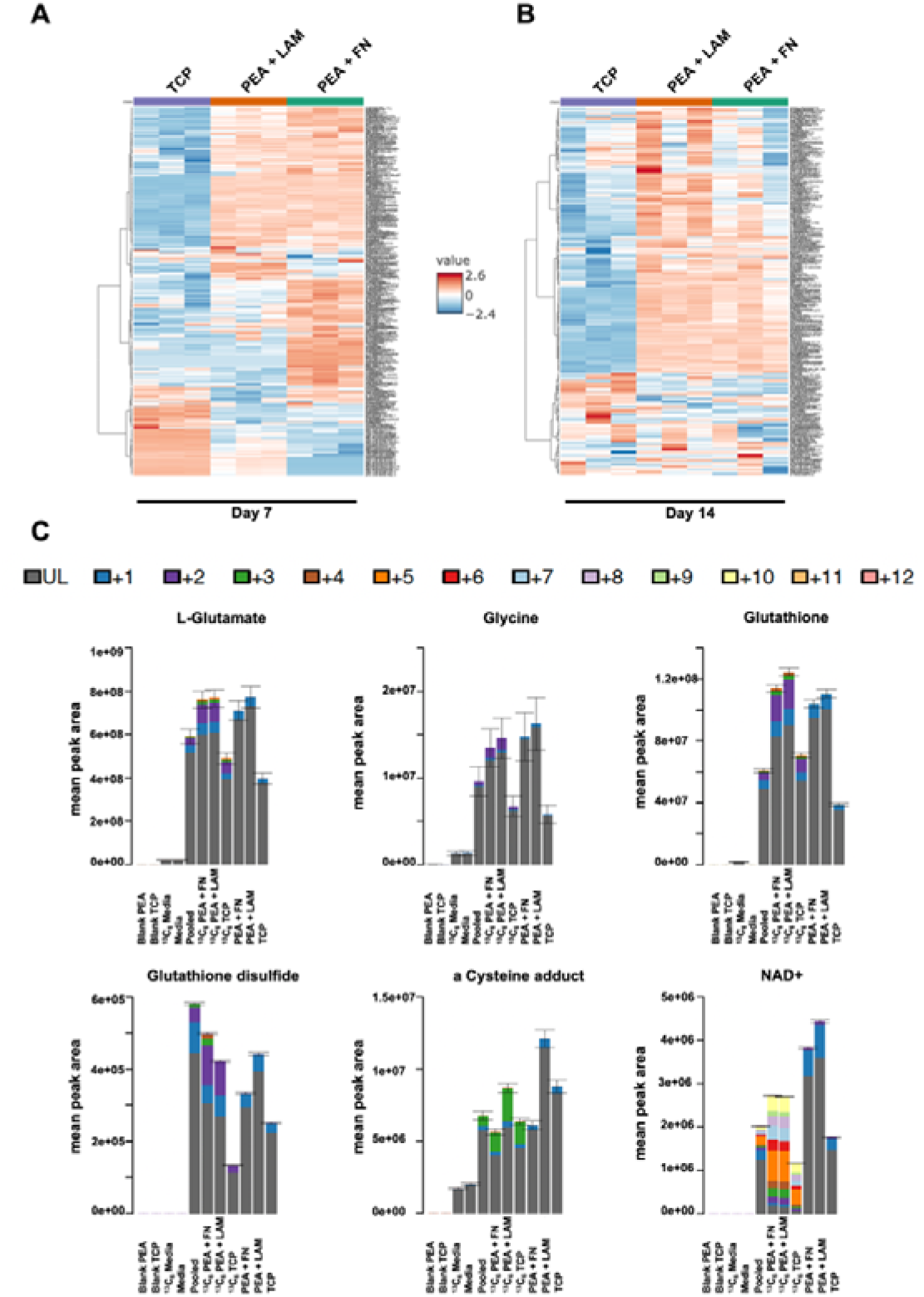
Assessment of metabolomic differences in MSCs at day 7 and day 14, and isotopologue distributions of selected metabolites following stable isotope tracing at day 14. Heatmaps displaying total metabolomic expression across conditions, with log_10_-transformed peak intensities clustered using Ward’s linkage and Euclidean distance for each technical replicate. Colour scale bar: red represents upregulation, while blue indicates downregulation (A, B). Isotopologue distributions of L-glutamate, glycine, GSH, GSSG, cysteine adducts, and NAD^+^ (C). Control unlabelled samples were also assessed to evaluate the incorporation of environmental ^13^C -glucose in the samples. Blank samples (extraction solvent of TCP or PEA) confirm the absence of metabolites from the surfaces. ^13^C labelled and control unlabelled media controls validate the presence and metabolism of metabolites in and from the media, respectively. Pooled samples were used as baseline reference for comparison of metabolite intensities to reduce inter-sample variation. Colours represent metabolite isotopologues according to the number of incorporated ^13^C atoms derived from ^13^C -glucose: Black: unlabelled, Blue: +1, Purple: +2, Green: +3, Brown: +4, Orange: +5, Red: +6, Light Blue: +7, Violet: +8, Light Green: +9, Yellow: +10, Light Orange: +11, Pink: +12. Analysis was performed on four technical replicates derived from one biological sample (n=4).

**Figure S5.**
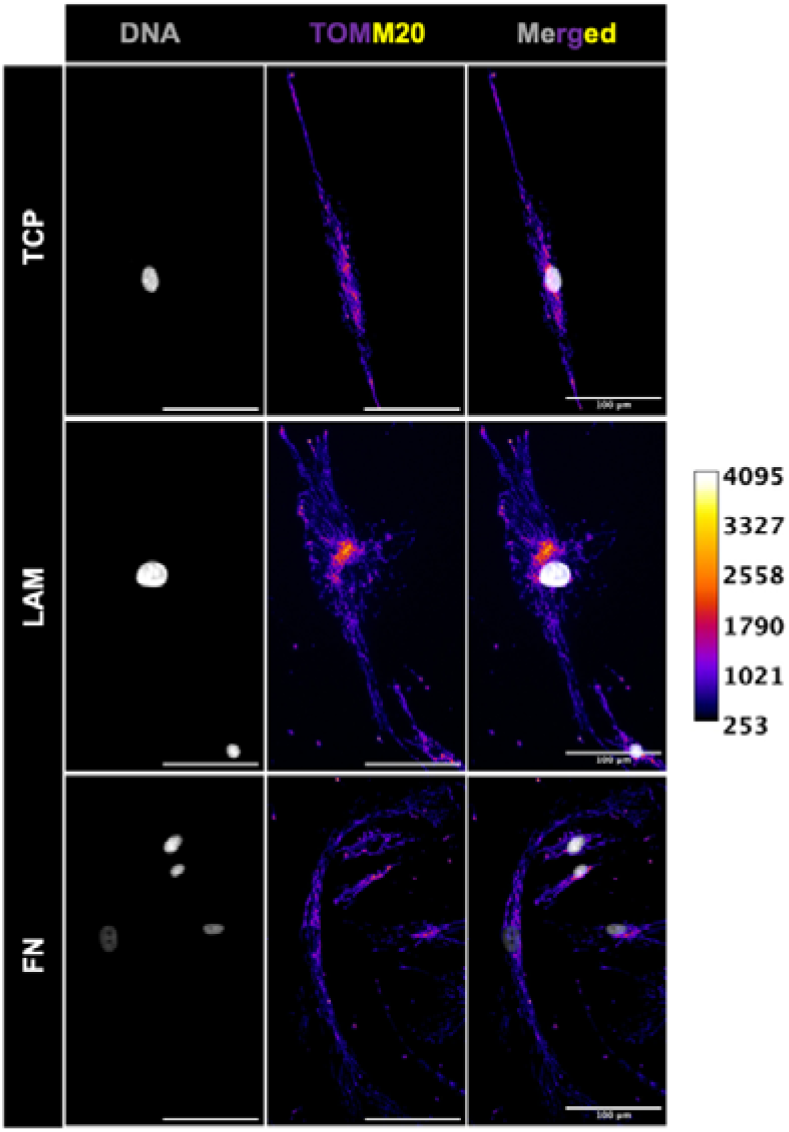
Assessment of mitochondrial localisation at day 3. Identification of mitochondrial clustering and perinuclear enrichment via quantitative assessment of fluorescence intensity: Fire look-up table visualisation: colour intensity represents relative TOMM20 fluorescence signal, with warmer colours (yellow–white) indicating higher mitochondrial signal intensity and cooler colours (purple–blue) indicating lower signal intensity, as indicated by the calibration bar. Scale Bar = 100 µm. Representative images out of 3 biological replicates.

**Figure S6.**
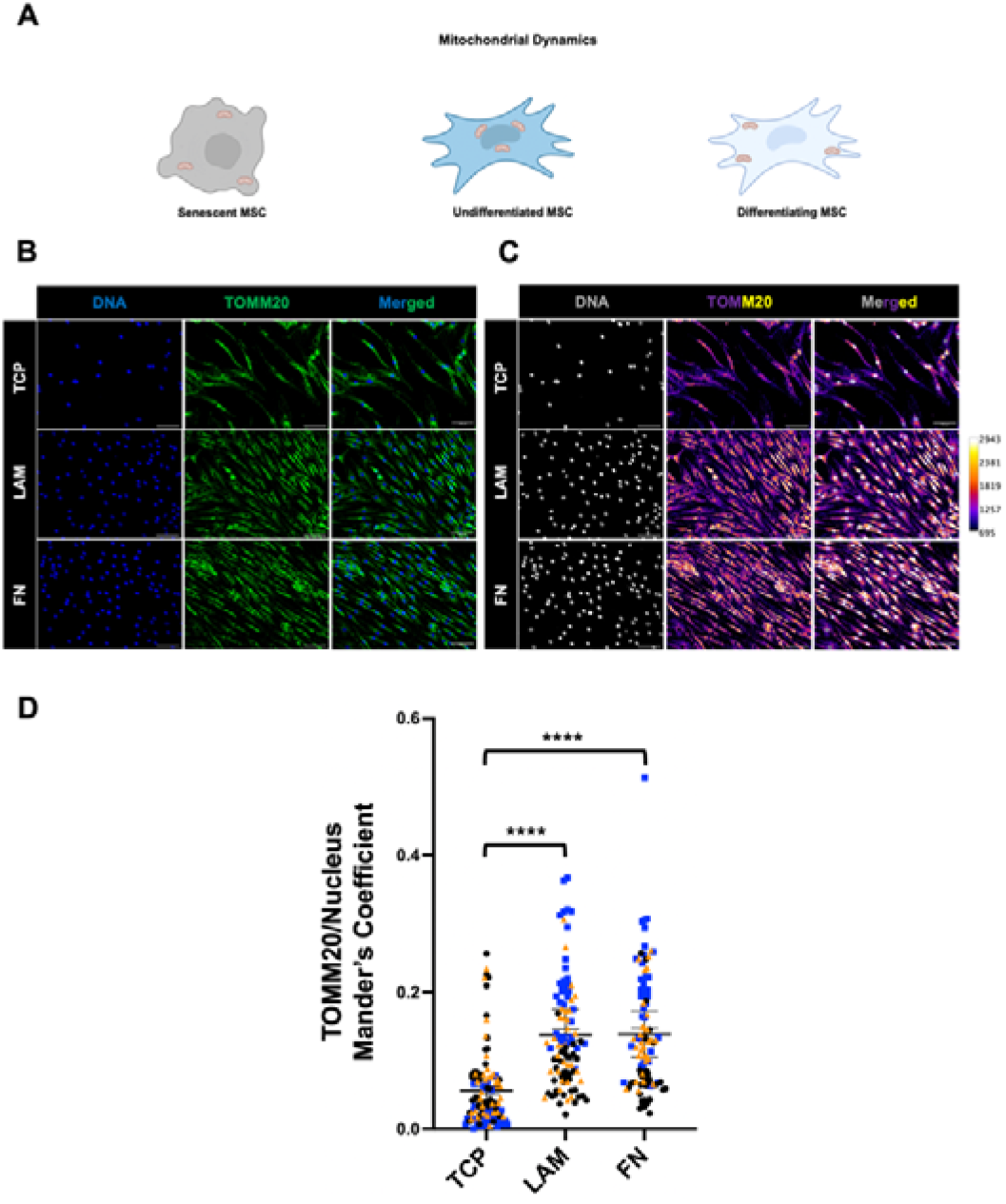
Assessment of mitochondrial localisation at day 21. Schematic representation of mitochondrial dynamics in MSCs at different states (**A**). MSC staining for TOMM20 (**B**). Identification of mitochondrial clustering and perinuclear enrichment via quantitative assessment of fluorescence intensity: Fire look-up table visualisation: colour intensity represents relative TOMM20 fluorescence signal, with warmer colours (yellow–white) indicating higher mitochondrial signal intensity and cooler colours (purple–blue) indicating lower signal intensity, as indicated by the calibration bar (**C**). Measurement of TOMM20 localisation at day 21 (**D**). The different coloured symbols on super plots represent the mean of every condition for every experiment. Smaller symbols represent the technical replicates of every experiment, accordingly. Blue and green stains represent the DNA, and protein of interest, respectively. Scale Bar = 100 µm. Representative images out of 3 biological replicates. Statistical analysis was performed via One-way ANOVA with Tukey’s multiple comparisons. *: p ≤ 0.05, **: p ≤ 0.01, ***: p ≤ 0.001, ****: p ≤ 0.0001. Data is presented as (mean ± SEM) from 3 biological replicates, (N=3).

**Figure S7.**
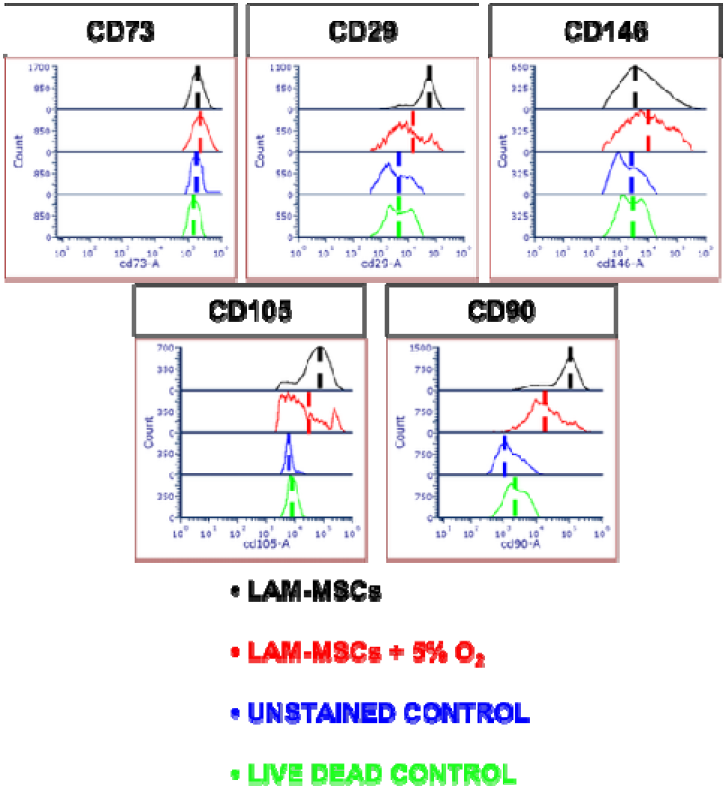
Histograms for ISCT flow cytometry analysis. Quantification of median fluorescence intensity, representative histograms out of 3 biological replicates. Histograms indicate changes in fluorescence intensity in the according markers. A right shift of the curve indicates an upregulation in protein expression. Dashed lines signify the middle of the curve. Black curves correspond to LAM-MSCs while red to LAM-MSCs cultured under 5% O_2_. Blue and green curves refer to unstained and live-dead controls, respectively. Data is presented as heatmaps from 3 biological replicates, (N=3).

**Figure S8.**
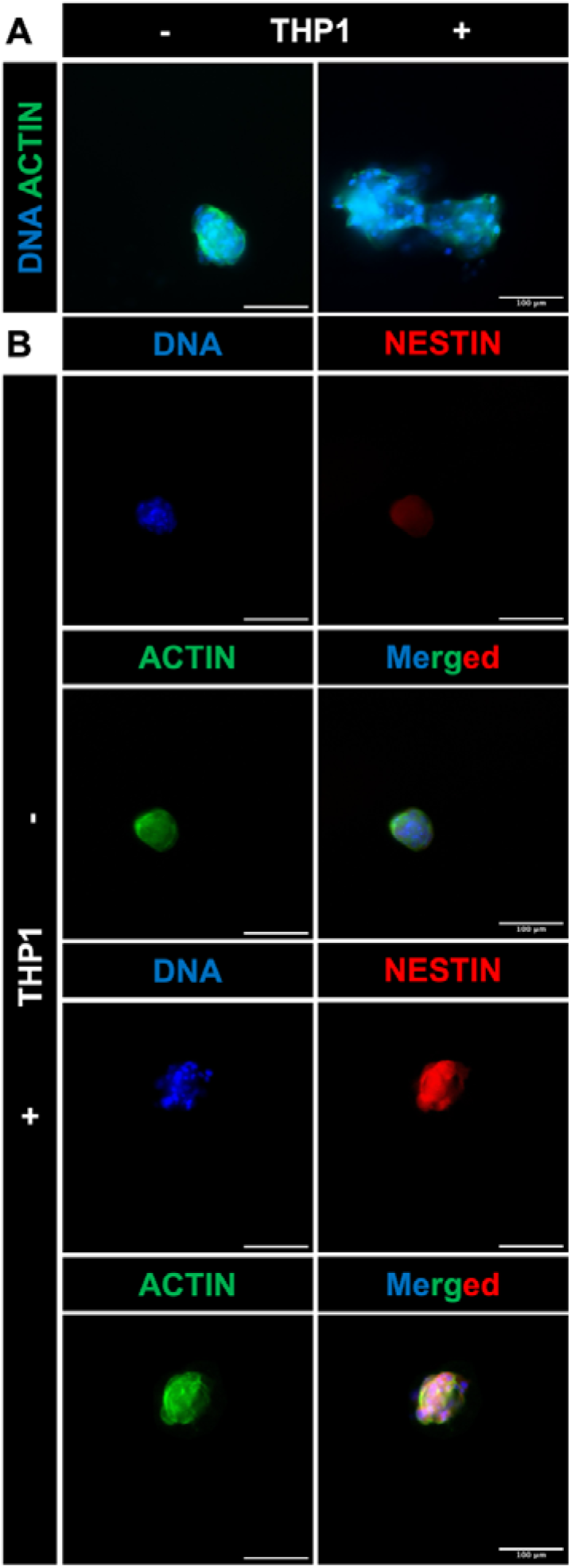
Staining for DNA, F-actin, and nestin on remodelled spheroids. Representative merged images of MSC spheroids cultured in the absence or presence of THP-1 cells (**A**). Spheroid staining for nestin (**B**). Green, blue, and red stains represent the F-actin, DNA, and protein of interest, respectively. Scale Bar = 100 µm. Representative images out of 3 biological replicates.

**Figure S9.**
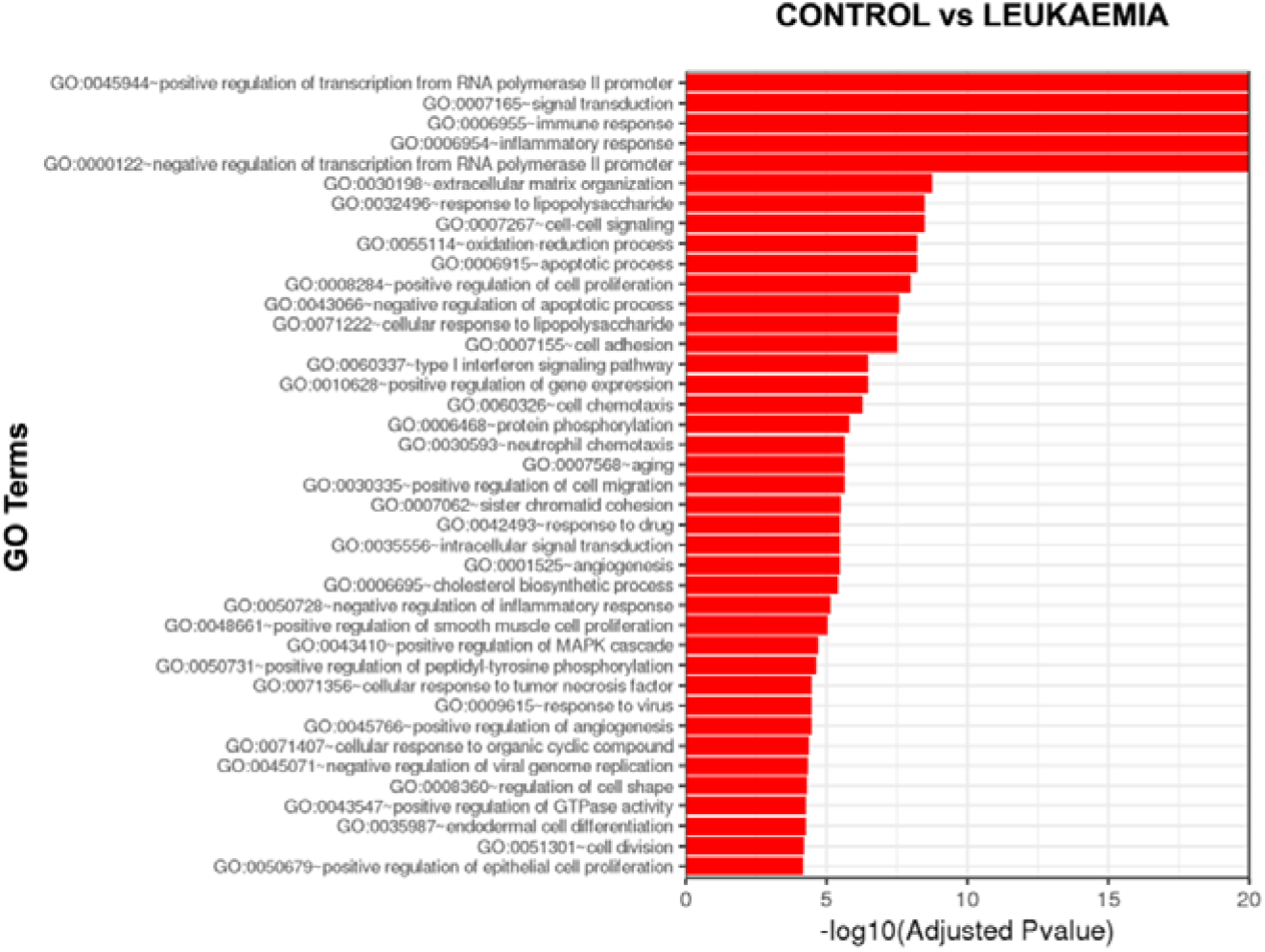
Gene ontology enrichment analysis of differentially expressed genes between control and leukaemic stroma. GO analysis. Significantly differentially expressed genes were clustered by their gene ontology and the enrichment of gene ontology terms was tested using Fisher exact test (GeneSCF v1.1-p2). Figure shows gene ontology terms, if any, that are significantly enriched with an adjusted P-value less than 0.05 in the differentially expressed gene sets (up to 40 terms). Analysis was performed on four technical replicates derived from one biological sample (n=4).

**Figure S10.**
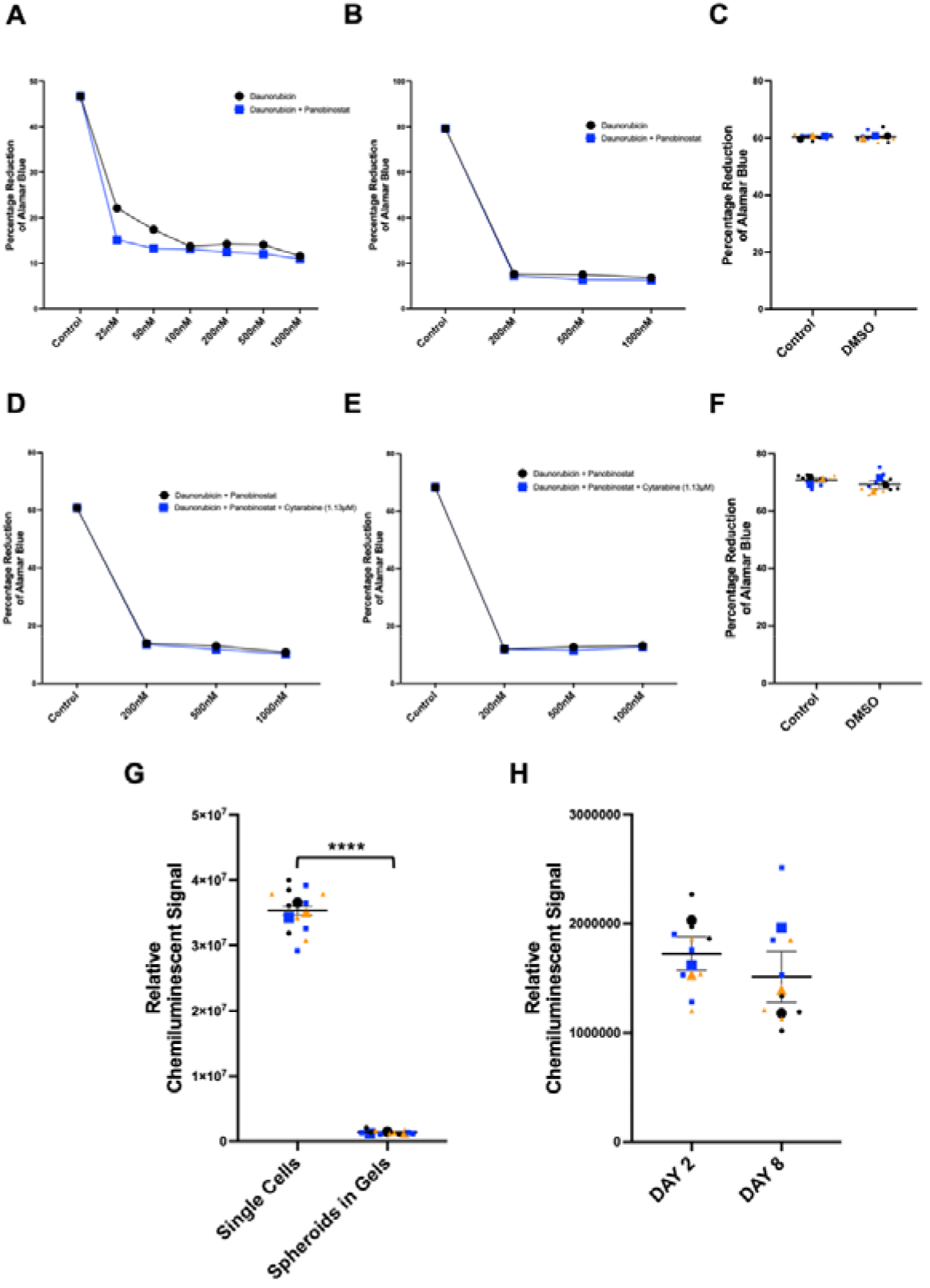
Investigation of chemotherapy effects on THP-1s and MSCs. Assessment of daunorubicin and panobinostat treatment effects on THP-1s for 2 days (**A**) and 4 days (**B**). DMSO control treatments on day 2 (**C**). Assessment of daunorubicin, panobinostat, and cytarabine treatment effects on THP-1s for 2 days (**D**) and 4 days (**E**). DMSO control treatments on day 4 (**F**). Comparison of ATP-associated luminescence between MSC monolayers and spheroids (**G**). Assessment of spheroid ATP-associated luminescence at days 2 and 8 (**H**). The different coloured symbols on graphs represent the mean of every condition for every experiment. Smaller symbols represent the technical replicates of every experiment, accordingly. Statistical analysis was performed via Student’s t-test or One-way ANOVA with Tukey’s multiple comparisons. *: p ≤ 0.05, **: p ≤ 0.01, ***: p ≤ 0.001, ****: p ≤ 0.0001. Data is presented as (mean ± SEM) from 3 biological replicates, (N=3).

**Figure S11.**
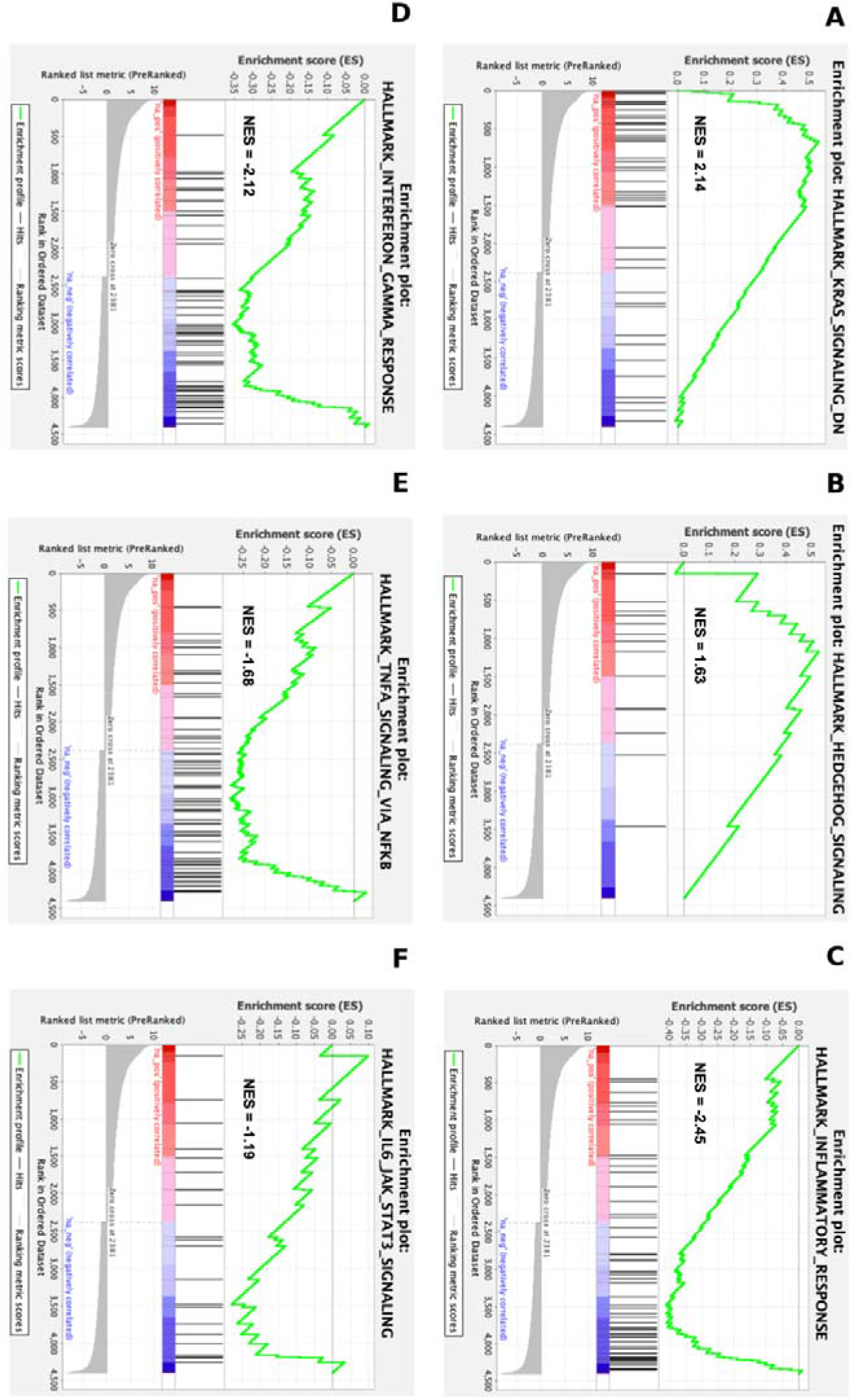
An expanded GSEA version of chemotherapy-treated leukaemia-remodelled MSC spheroids. Panels (**A**–**F)** show enlarged GSEA enrichment plots corresponding to Figure 6(**J**–**O)**, respectively, comparing MSCs from leukaemic conditions with MSCs following chemotherapy treatment (leukaemia + chemotherapy). Analysis was performed on four technical replicates derived from one biological sample (n=4).

**Figure S12.**
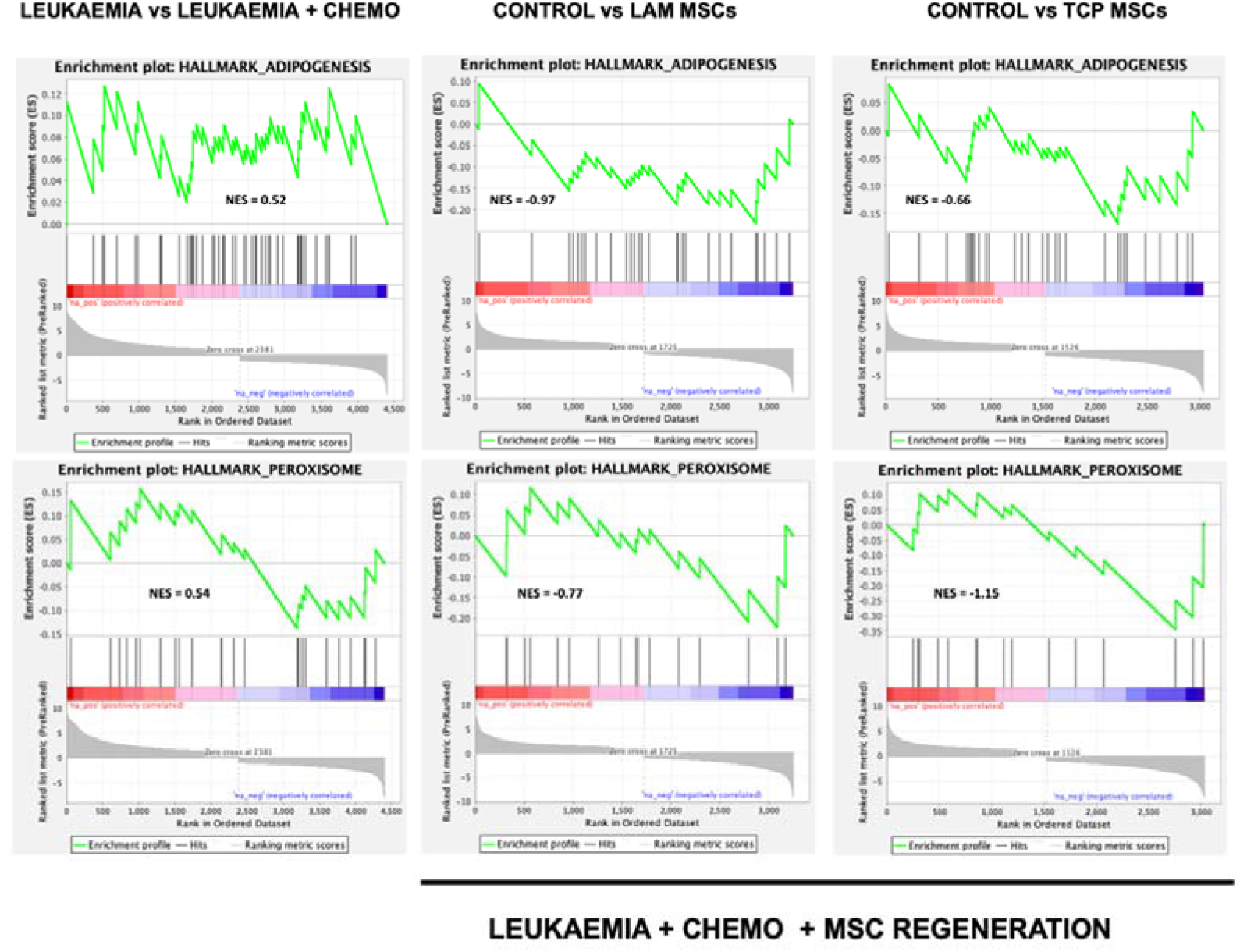
MSC-based cellular therapies promote regeneration of the leukaemic niche following chemotherapy exposure. GSEA was performed to evaluate transcriptional changes in MSCs following chemotherapy and subsequent regeneration with two MSC-based strategies: LAM-MSCs and TCP-MSCs. Enrichment plots display the expression of hallmark pathways related to adipogenesis and peroxisome signalling, both of which are associated with MSC functional restoration and niche support. Comparison of leukaemia vs. leukaemia□+□chemo shows a slight tendency towards adipogenesis and stress post-chemotherapy. In contrast, treatment with LAM-MSCs or TCP-MSCs results in significant downregulation of adipogenic and peroxisome gene sets relative to chemo-only conditions, as indicated by negative normalised enrichment scores (NES). These findings suggest that MSC therapies modulate the niche toward a less adipogenic, less oxidatively stressed state, potentially restoring homeostatic stromal function disrupted by leukaemic remodelling and chemotherapy. Analysis was performed on four technical replicates derived from one biological sample (n=4).

**Figure S13.**
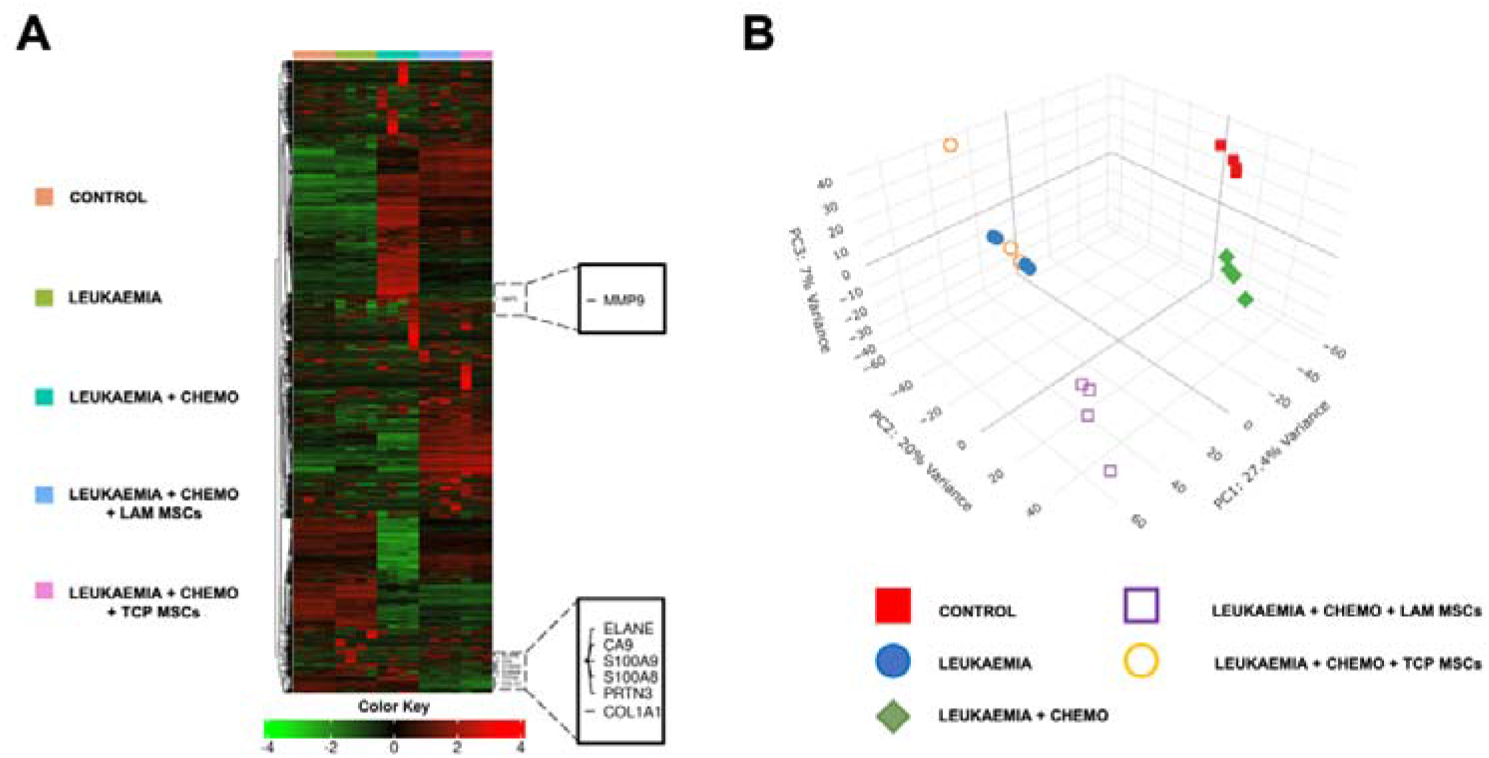
Transcriptomic profiling reveals distinct gene expression signatures across control, leukaemic, treated, and regenerated niche conditions. Hierarchical clustering heatmap of the top 2000 most variable genes across samples. The heatmap was generated using log₂-transformed transcripts per million (TPM) values. Gene expression profiles were analysed using Pearson correlation and average linkage hierarchical clustering. Each row represents a gene, and each column represents a sample. The colour scale represents relative expression levels, with red indicating higher and green indicating lower expression (Z-score scale from –4 to +4). Colour-coded bars above the heatmap denote sample groupings. This analysis highlights distinct gene expression signatures across experimental conditions (A). 3D PCA plot comparing the clustering of transcriptomic profiles across all experimental conditions. PC1, PC2, and PC3 represent the top three principal components, indicating 27.4%, 20%, and 7% of the total variance, respectively. The PC1 vs. PC2 comparison shows the primary separation between the control and leukaemic niches with those treated with chemotherapy and regenerated via cellular therapy, signifying that treatment shows the greatest variance. The PC3 vs. PC1 comparison further distinguishes the control from the leukaemic niche and the therapies, highlighting differences in the leukaemic remodelling. While both MSC therapies alter the transcriptomic profile following chemotherapy, only LAM MSC-treated samples exhibit a shift towards the control cluster along PC2, suggesting a more extensive restoration of the healthy stromal transcriptional programme. In contrast, TCP MSC-treated samples remain closely aligned with the leukaemic transcriptional profile, indicating a comparatively limited transcriptomic reversion towards the healthy state. Red square: control, blue circle: leukaemia, green rhombus: leukaemia + chemotherapy, purple square: LAM-MSCs, orange circle: TCP-MSCs (B). Analysis was performed on four technical replicates derived from one biological sample (n=4).

**Figure S14.**
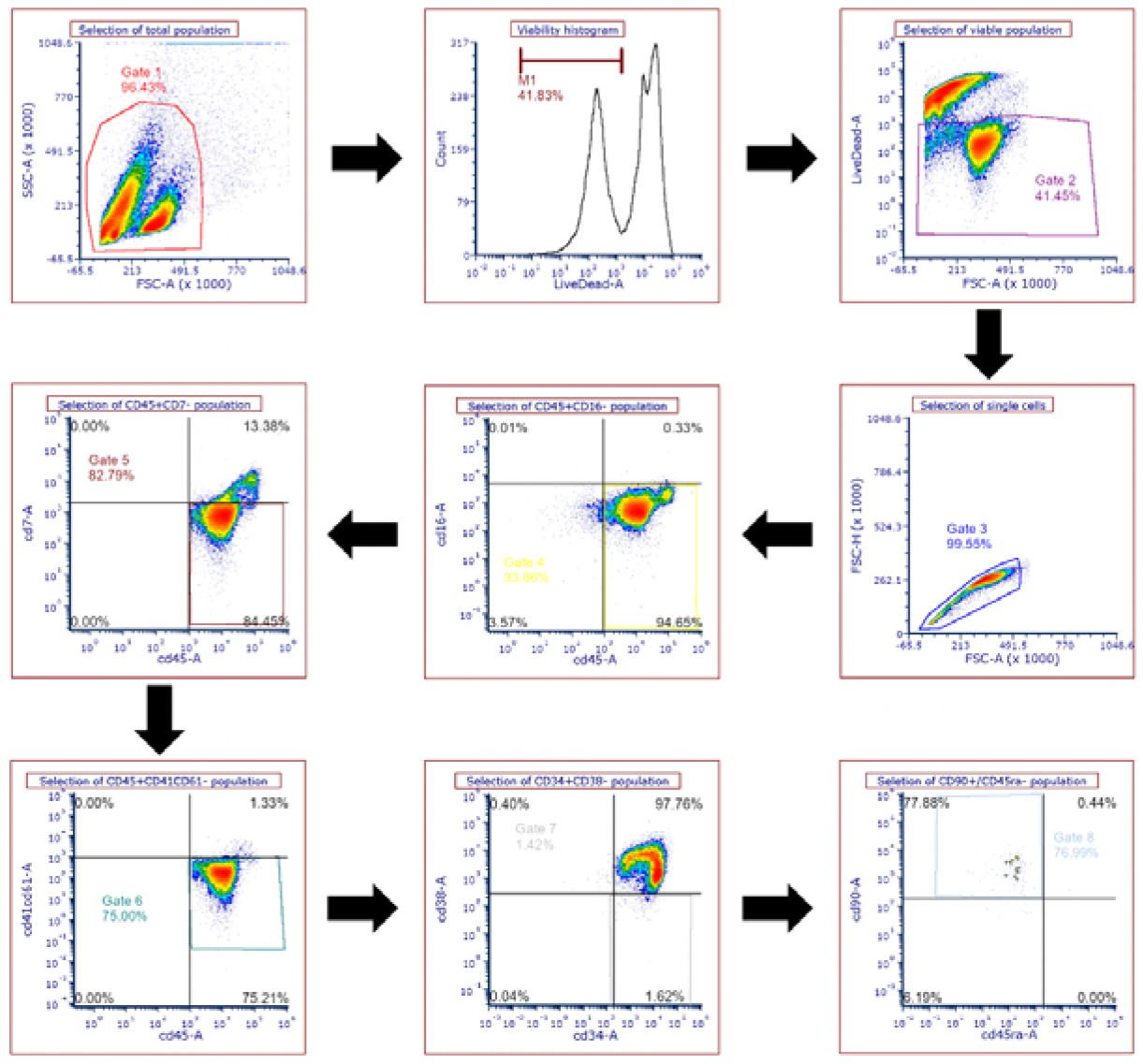
Gating strategy of HSC flow cytometry. Following single-cell selection, dead cells were excluded (dead-cell fluorescence peak at 10⁴·⁵) before subsequent assessment of marker expression. CD45⁺ cells were then selected, followed by exclusion of cells expressing CD16, CD7, or CD41/CD61 during subsequent gating. Within the remaining CD45⁺ population, CD34⁺CD38 cells were identified as an HSC/progenitor-enriched population. Finally, CD90⁺CD45ra cells within the CD45⁺CD34⁺CD38 population were identified as cells exhibiting an LT-HSC-enriched immunophenotype.

**Figure S15.**
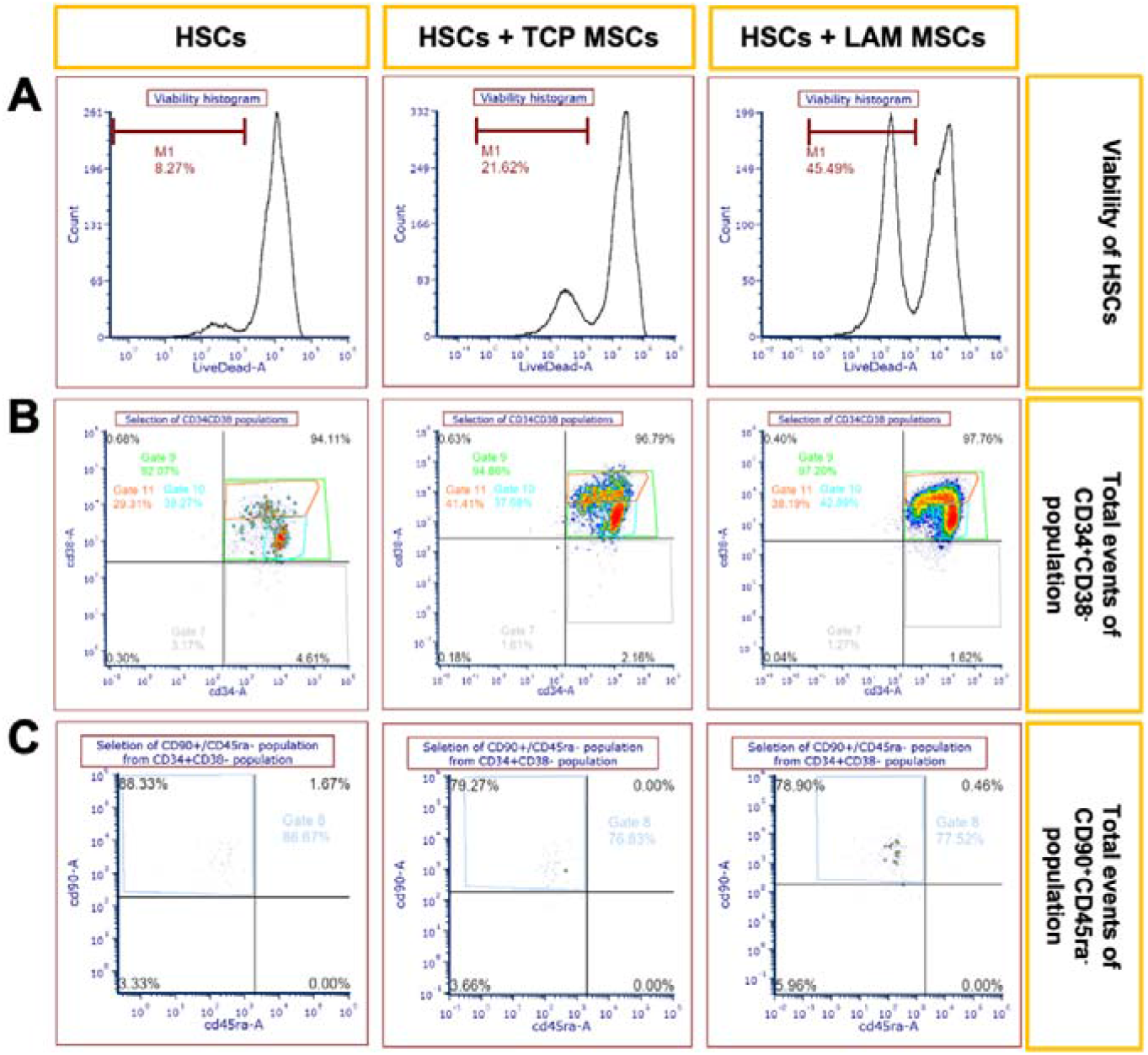
Flow cytometry analysis of HSC viability, haematopoietic progenitor, and stem cells following co-culture with MSCs. Representative viability histograms of HSCs (**A**). Representative dot plots showing the gating strategy used to identify haematopoietic progenitor cells (CD34^+^CD38^-^ population) (**B**). Representative dot plots showing the gating strategy used to identify stem cells (CD34^+^CD38^-^ CD90⁺CD45ra^-^ population) (**C**).

**Figure S16.**
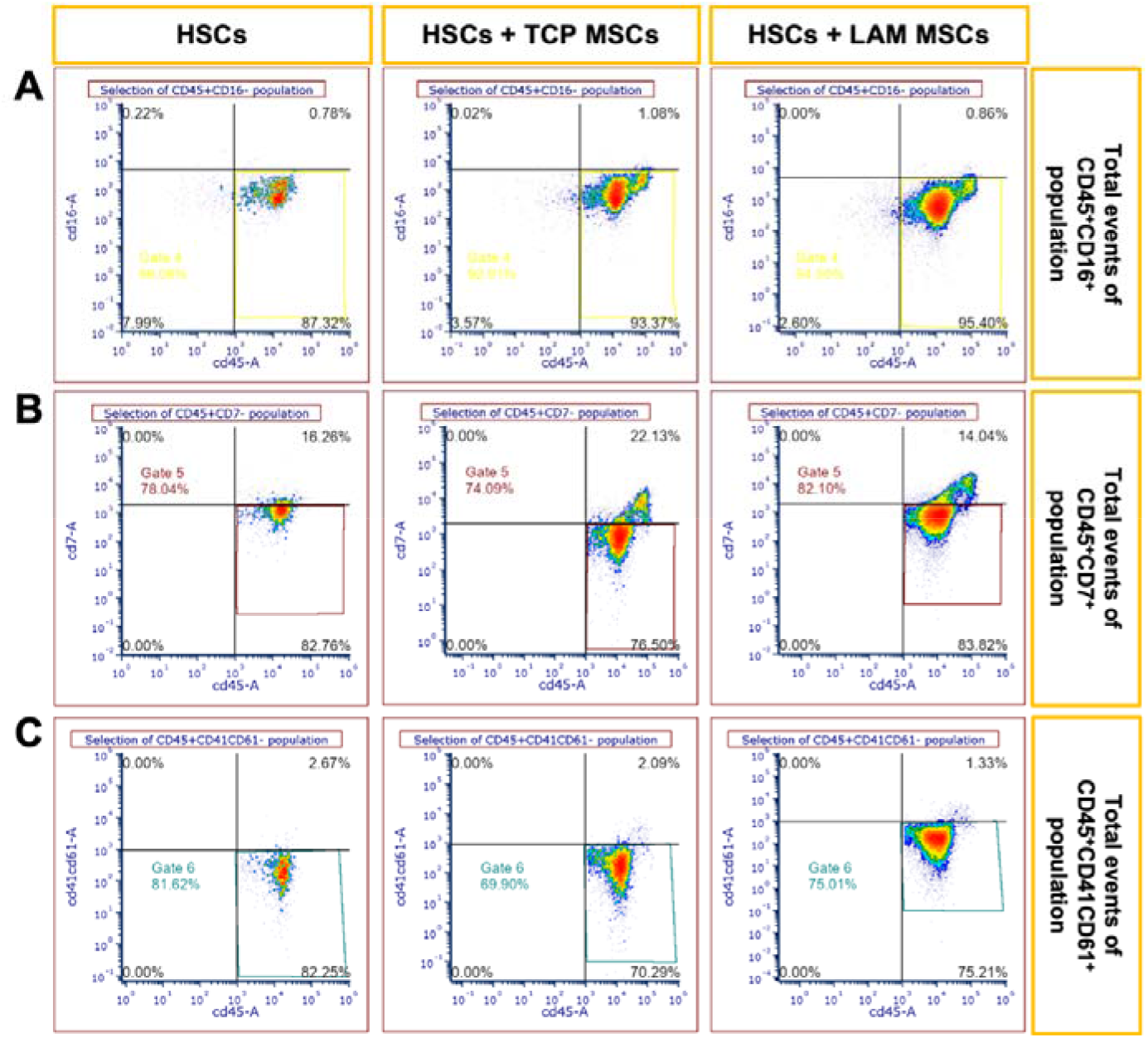
Flow cytometry analysis of NK-cell, T-cell, and megakaryocyte differentiation following co-culture with MSCs. Representative dot plots showing the gating strategy used to identify NK-cells (CD45^+^CD16^+^ population) (**A**). Representative dot plots showing the gating strategy used to identify T-cells (CD45^+^CD7⁺ population) (**B**). Representative dot plots showing the gating strategy used to identify megakaryocyte cells (CD45^+^CD41CD61⁺ population) (**C**).

**Figure S17.**
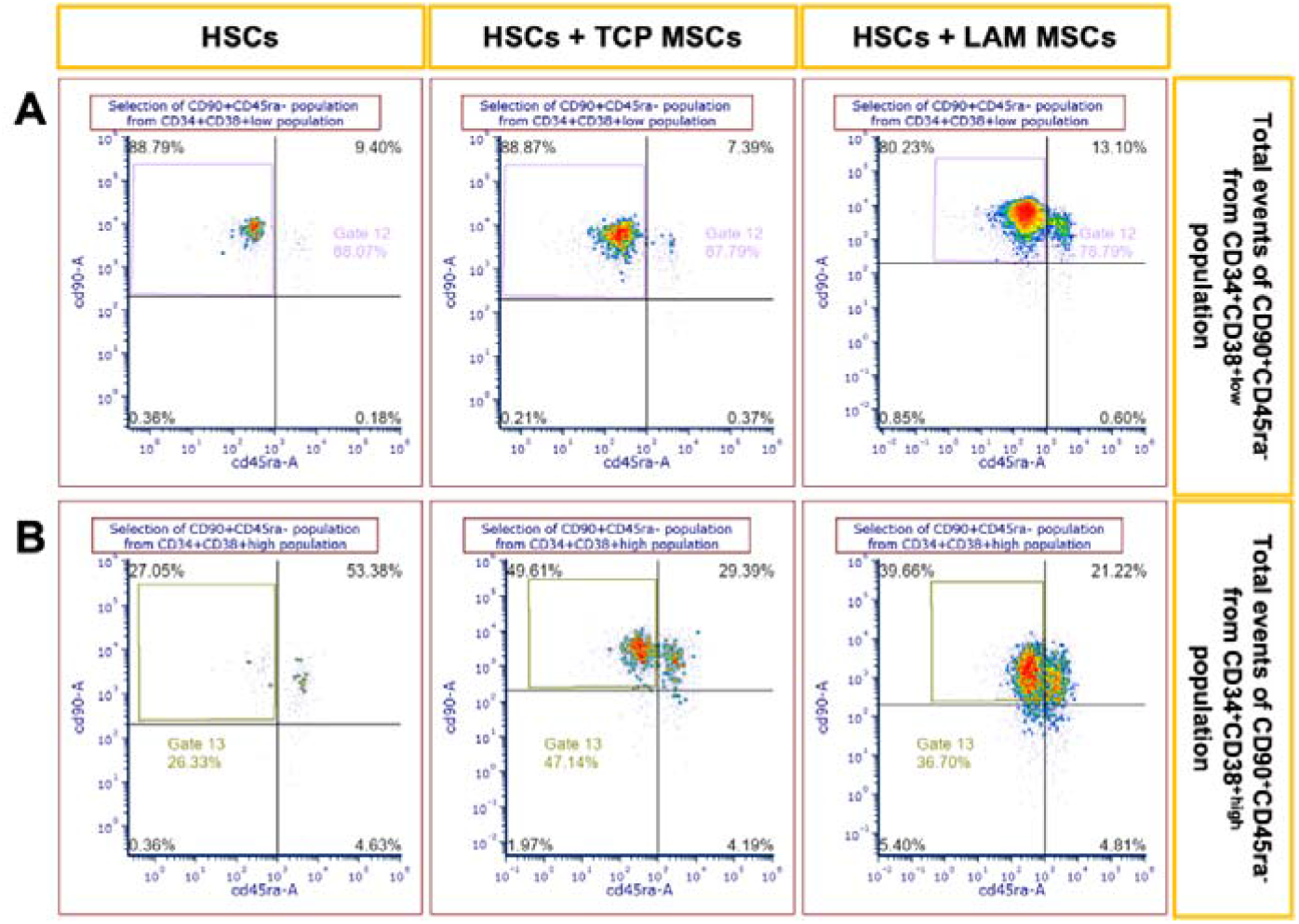
Flow cytometry identification of primitive haematopoietic progenitor subpopulations following co-culture with MSCs. Representative dot plots showing the identification of CD90⁺CD45ra^-^ cells within the CD34⁺CD38^⁺low^ population (**A**). Representative dot plots showing the identification of CD90⁺CD45ra^-^ cells within the CD34⁺CD38^⁺high^ population (**B**).

**Figure S18.**
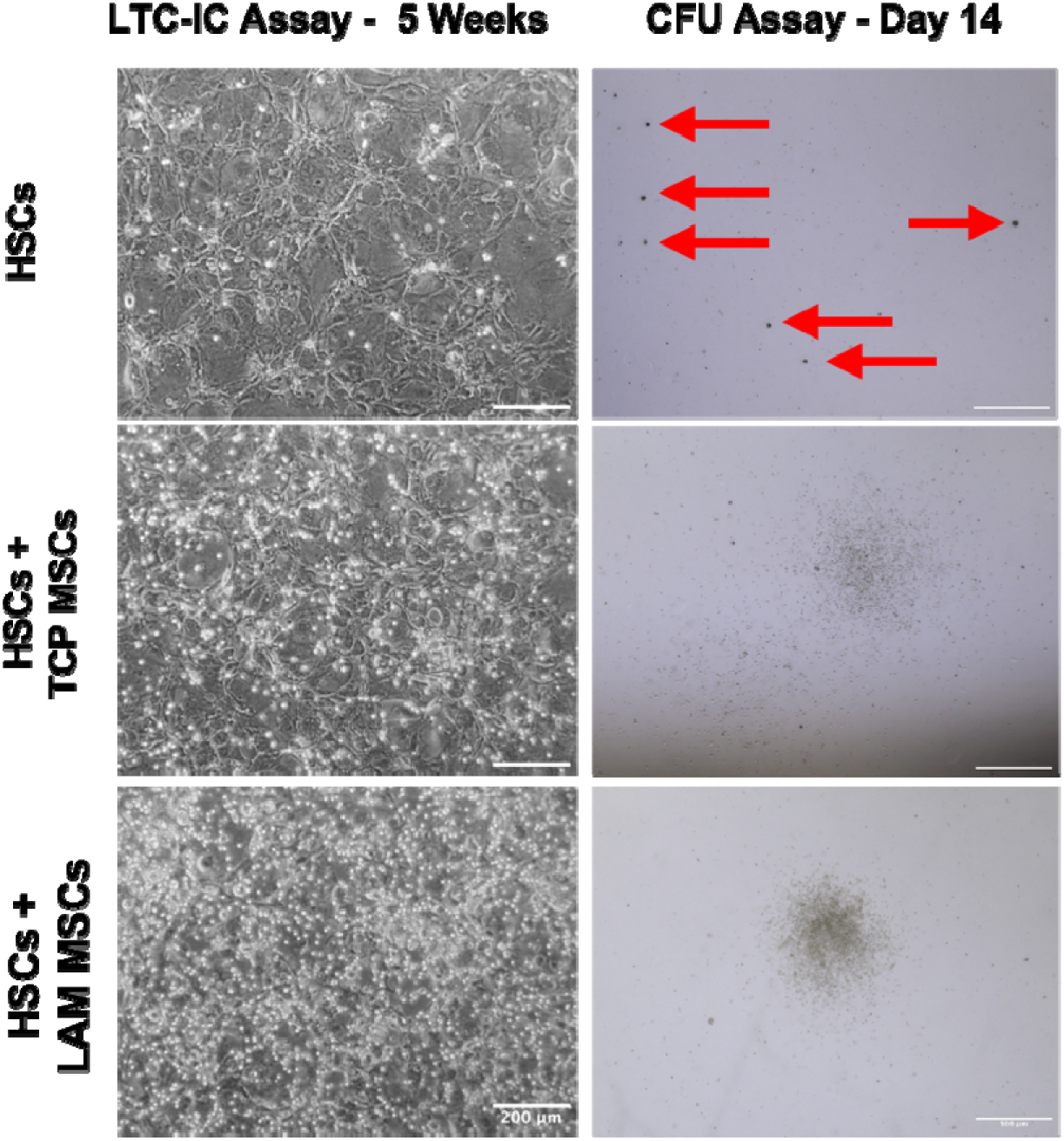
Long-term HSC maintenance followed by assessment of colony-forming capacity. Representative brightfield images of HSCs on feeder layers post 5-week LTC-IC culture. Scale bars: 200 µm. Representative brightfield images of colonies formed after 14 days of CFU assay post 5-week LTC-IC culture. Scale bars: 500 µm. HSC monocultures exhibited a predominance of small, dispersed erythroid colonies (indicated by red arrows), while co-culture with TCP-MSCs and LAM-MSCs resulted in denser colony formation with a more balanced distribution between erythroid and myeloid outputs.

**Table S1.** Leukaemia-associated niche remodelling gene set.

| Gene Name | Associated Function |
|---|---|
| 24-Dehydrocholesterol Reductase (DHCR24) | Cholesterol biosynthesis.<br>CAFs undergo lipidomic reprogramming towards fatty acid synthesis to promote colorectal cancer progression. |
| Aldo-Keto Reductase Family 1 Member C2 (AKR1C2) | Oxidative-electrophilic-xenobiotic stress response. |
| BCL2 Interacting Protein 3 (BNIP3) | Hypoxia response promoting survival via autophagy. |
| C-X-C Motif Chemokine Ligand 1 (CXCL1) | Chemoresistance in breast cancer. |
| C-X-C Motif Chemokine Ligand 2 (CXCL2) | Promotion of AML survival. |
| C-X-C Motif Chemokine Ligand 3 (CXCL3) | CAF-secreted CXCL3 induced chemoresistance in renal cell carcinoma. |
| C-X-C Motif Chemokine Ligand 6 (CXCL6) | Radio-resistance in hepatocellular carcinoma.<br>Immunotherapy resistance in cholangiocarcinoma. |
| C-X-C Motif Chemokine Ligand 8 (CXCL8) | Chemoresistance in AML. |
| Carbonic Anhydrase 9 (CA9) | Hypoxia – Acidosis Response. |
| CCAAT/Enhancer Binding Protein Delta (CEBPD) | CEBP family members are stress-responsive in stromal cells. |
| Cellular Communication Network Factor 1 (CCN1) | CCN1 (matricellular protein) in CAFs helps create a pro-metastatic, proangiogenic, fibrotic tumour microenvironment in melanoma. |
| Decidual Protein Induced by Progesterone 1 (DEPP1) | Hypoxia/ROS Response – Autophagy Induction.<br>Associated with greater CAF infiltration in gastric cancer. |
| DNA Damage Inducible Transcript 4 (DDIT4) | DNA damage – ER stress response. |
| Dual Specificity Phosphatase 1 (DUSP1) | Negative regulator of osteoclast formation.<br>Resolution of inflammation. |
| Early Growth Response 1 (EGR1) | Oxidative stress response. |
| Elastase, Neutrophil Expressed (ELANE) | Enhanced expression in Shwachman-Diamond Syndrome (SDS)-MSCs. |
| Fibroblast Growth Factor 7 (FGF7) | CAF-secreted FGF7 promotes ovarian cancer progression. |
| H1.0 Linker Histone (H1F0) | Stromal Senescence.<br>Chromatin Compaction.<br>Epigenetic Regulation. |
| Heme Oxygenase 1 (HMOX1) | Stress-Responsive.<br>Cytoprotective.<br>Expressed in TME promoting disease progression. |
| Immediate Early Response 3 (IER3) | Inflammatory stress response. |
| Interferon Induced Transmembrane Protein 1 (IFITM1) | Interferon Response.<br>Interferons derived from solid tumours promote CAF expansion and reprogramming. |
| Interleukin 1 Alpha (IL1A) | Promotes leukaemic cell proliferation and tumour progression. |
| Interleukin 1 Beta (IL1B) | AML progression and chemoresistance. |
| Interleukin 11 (IL11) | Enhances AML proliferation. |
| Interleukin 24 (IL24) | Tumour suppressing response.<br>Facilitates apoptosis of leukaemic cells. |
| Interleukin 32 (IL32) | Tumour initiation, proliferation, and maintenance.<br>Implicated in AML progression and chemoresistance. |
| Interleukin 6 (IL-6) | Chemoresistance in AML. |
| JunB Proto-Oncogene, AP-1 Transcription Factor Subunit (JUNB) | ER stress protective factor. |
| Leukaemia Inhibitory Factor (LIF) | Anti-leukaemic effect in myeloid malignancies. |
| Long Intergenic Non-Protein Coding RNA 1119 (LINC01119) | Stromal cells deliver LINC01119 to ovarian cancer cells via exosomes, facilitating disease progression.<br>Stromal cells induce its expression in breast cancer cells, facilitating disease progression. |
| Marker of Proliferation Ki-67 (MKI67) | Proliferation Marker. |
| NADH:Ubiquinone Oxidoreductase Subunit A4 Like 2 (NDUFA4L2) | Hypoxia Response. |
| Nicotinamide N-Methyltransferase (NNMT) | ER stress response. |
| Nuclear Receptor Subfamily 4 Group A Member 1 (NR4A1) | Oxidative stress protective factor. |
| Nuclear Receptor Subfamily 4 Group A Member 2 (NR4A2) | Oxidative stress response. |
| Pentraxin 3 (PTX3) | ECM remodelling.<br>Upregulation in breast cancer CAFs facilitates disease progression. |
| Prostaglandin-Endoperoxide Synthase 2 (PTGS2) | Oxidative stress response |
| Proteinase 3 (PRTN3) | Enhanced expression in SDS-MSCs. |
| S100 Calcium Binding Protein A8 (S100A8) | Enhanced expression in SDS-MSCs. Inflammatory alarmin causing genotoxic stress to HSPCs.<br>MSC-derived S100A8 promotes leukaemic cell survival and chemoresistance.<br>Fibroblasts and CAFs can be reprogrammed from tumoral S100A8/9 to produce inflammation and promote chemoresistance. |
| S100 Calcium Binding Protein A9 (S100A9) | Enhanced expression in SDS-MSCs. Inflammatory alarmin causing genotoxic stress to HSPCs.<br>Fibroblasts can be reprogrammed from tumoral S100A8/9 to produce inflammation.<br>S100A9 induces senescence in MSCs in MDS. |
| Squalene Epoxidase (SQLE) | Cholesterol biosynthesis.<br>CAFs undergo lipidomic reprogramming towards fatty acid synthesis to promote colorectal cancer progression. |
| Stanniocalcin 1 (STC1) | Hypoxia – Acidosis – Oxidative stress response |
| Thioredoxin Interacting Protein (TXNIP) | ER stress response.<br>Oxidative stress and inflammation regulator. |
| Tissue Factor Pathway Inhibitor 2 (TFPI2) | ECM remodelling.<br>Elevated in breast cancer CAFs. |
| Tribbles Pseudokinase 3 (TRIB3) | ER stress response. |
| Tumour Necrosis Factor Alpha–Induced Protein 6 (TNFAIP6) | Chemoresistance in melanoma and osteosarcoma. |
| Tumour Necrosis Factor Receptor Superfamily Member 11B (TNFRSF11B) | Decoy receptor that inhibits osteoclastogenesis. |
| Tumour Necrosis Factor Receptor Superfamily Member 25 (TNFRSF25) | Survival Response.<br>Promotion of fibrosis in solid tumours. |
| Vascular Cell Adhesion Molecule 1 (VCAM-1) | CAF-derived VCAM-1 promotes tumour cell proliferation and invasion. |

**Table S2.** Chemotherapy-induced gene expression changes.

| **Gene Name** | **Associated Function** |
|---|---|
| Amphiregulin (AREG) | Anti-inflammatory response.<br>Cytoprotective – tissue repair activity. |
| Caveolae Associated Protein 3 (CAVIN-3) | Membrane organization. |
| Cell Migration Inducing Hyaluronidase 1 (CEMIP) | ECM – tissue remodelling. |
| Chromodomain Helicase DNA Binding Protein 3 (CHD3) | Subunit of the Nucleosome Remodelling and Deacetylase (NuRD) complex.<br>DNA damage response/repair.<br>Chromatin remodelling. |
| Collagen Triple Helix Repeat Containing 1 (CTHRC1) | ECM – tissue remodelling. |
| Collagen Type I Alpha 1 Chain (COL1A1) | Deposition implicated in myelofibrosis. |
| Collagen Type I Alpha 2 Chain (COL1A2) | Fibrotic effector in systemic sclerosis fibroblasts. |
| Collagen Type III Alpha 1 Chain (COL3A1) | Implicated in reticulin deposition in myelofibrosis. |
| Collagen Type V Alpha 2 Chain (COL5A2) | Involved in collagen fibrillogenesis in wounds.<br>CAF-derived mediator of tumour progression. |
| Collagen Type X Alpha 1 Chain (COL10A1) | CAF-COL10A1 promotes tumour EMT and metastasis. |
| Collagen Type XIII Alpha 1 Chain (COL13A1) | Implicated with a matrix fibroblast population in pulmonary fibrosis. |
| Collagen Type XVI Alpha 1 Chain (COL16A1) | Produced by myofibroblasts.<br>Upregulated during the progression from liver fibrosis to hepatocellular carcinoma. |
| Collagen Type XXVII Alpha 1 Chain (COL27A1) | Upregulated during the progression from liver fibrosis to hepatocellular carcinoma. |
| Creatine Kinase B-type (CKB) | Cytoprotective under metabolic stress by supporting ATP homeostasis.<br>Cytoprotective against hypoxia/reoxygenation stress.<br>Cytoprotective role against oxidative stress. |
| Cytoglobin (CYGB) | Hypoxia-responsive.<br>Antioxidant, anti-fibrotic effector. |
| Epithelial Membrane Protein 3 (EMP3) | Membrane organization.<br>Cell adhesion. |
| Glutathione Peroxidase 3 (GPX3) | Antioxidant effector. |
| Glycophorin C (GYPC) | Membrane organization. |
| Heat Shock Protein Family B (Small) Member 7 (HSPB7) | Cytoprotective – polyQ aggregation/misfolded protein suppressor. |
| Inhibitor of DNA Binding 3, HLH Protein (ID3) | Anti-fibrotic regulator. |
| Lysophosphatidic Acid Receptor 1 (LPAR1) | Mediates cytoskeletal changes, cell migration, proliferation, and survival.<br>Mediates tissue fibrosis.<br>CAF-LPAR1 promotes tumour proliferation and invasion. |
| Matrix Metallopeptidase 1 (MMP1) | Collagenase – collagen degradation. |
| Metallothionein 1X (MT1X) | Antioxidant effector. |
| NAD(P)H Quinone Dehydrogenase 1 (NQO1) | Oxidative stress response - Antioxidant effector. |
| Olfactomedin Like 2B (OLFML2B) | ECM-associated factor. |
| Paired Related Homeobox 1 (PRRX1) | Transcription factor that programs stromal fibroblasts into tumour-promoting myofibroblast-like CAFs.<br>CAF-PRRX1 promotes EMT and chemoresistance in tumours.<br>Controls fibroblast activation in wound healing, fibrosis, and cancer stroma. |
| Platelet Derived Growth Factor Receptor Beta (PDGFRB) | PDGF/PDGFR axis implicated in tumour–stroma/CAF signalling and chemoresistance in solid tumours. |
| S100 Calcium Binding Protein A4 (S100A4) *also known as* Fibroblast-Specific Protein 1 (FSP1) | CAF marker, also identified in AML-BM. |
| Serpin Family H Member 1 (SERPINH1) | ER stress effector. |
| Solute Carrier Family 12 Member 8 (SLC12A8) | Metabolic stress response. |
| Syndecan 4 (SDC4) | Cell-surface heparan sulfate proteoglycan involved in cell–matrix adhesion, migration.<br>SDC4 expression is modulated by anticancer therapy.<br>Identified in tumour stroma of colorectal cancer with poor prognosis.<br>Tumour stromal SDC4 has been associated with disease progression in solid tumours. |
| Thioredoxin Reductase 1 (TXNRD1) | Antioxidant effector. |
| Transcription Factor 25 (TCF25) *also known as* Nuclear localized protein-1 (Nulp1) | bHLH transcription factor regulating morphogenesis, cell fate, and differentiation.<br>Transcriptional repressor.<br>Cytoprotective regulator that attenuates stress-induced transcriptional remodelling.<br>Nutrient sensor - metabolic adaptation under glucose starvation. |
| WNT1 Inducible Signalling Pathway Protein 2 (WISP2) | Promotes proliferation. |

**Table S3.** Regeneration-associated gene set.

| Gene Name | Associated Function |
|---|---|
| Aldo-Keto Reductase Family 1 Member C1 (AKR1C1) | Cytoprotective against oxidative stress–induced ferroptosis. |
| Bone Morphogenetic Protein 2 (BMP2) | Osteoinductive marker. |
| C1q and Tumour Necrosis Factor Related Protein 1 (C1QTNF1) | Inflammation-associated adipokine. |
| C-X-C Motif Chemokine Ligand 12 (CXCL12) | Pro-inflammatory mediator via promoting the recruitment and retention of immune cells.<br>Pro-fibrotic activity on stromal cells. |
| C-X-C Motif Chemokine Ligand 16 (CXCL16) | Inflammatory-associated marker.<br>Immune cell recruitment.<br>Identified in CAFs. |
| C-X-C Motif Chemokine Ligand 5 (CXCL5) | Pro-inflammatory cytokine. |
| Family with Sequence Similarity 20 Member A (FAM20A) | Associated with osteogenic potential and mineralization. |
| Integrin-Binding Sialoprotein (IBSP) | Osteogenic marker. |
| Insulin-Like Growth Factor 2 (IGF2) | Osteoinductive enhancer. |
| Interleukin 33 (IL33) | CAF-derived IL-33 is an inflammatory alarmin that promotes tumour metastasis. |
| Interleukin 36 Beta (IL36B) | Pro-inflammatory cytokine. |
| Interleukin 6 Receptor (IL6R) | IL6R expression on stromal cells in tumours is associated with poor outcome. |
| Mesenteric Estrogen Dependent Adipogenesis (MEDAG) | Adipogenesis-associated factor. |
| Metallothionein 1F (MT1F) | MT1 isoform - cytoprotective against oxidative stress, inflammation, and DNA damage. |
| Metallothionein 1G (MT1G) | MT1 isoform - cytoprotective against oxidative stress, inflammation, and DNA damage. |
| Metallothionein 1H (MT1H) | MT1 isoform - cytoprotective against oxidative stress, inflammation, and DNA damage. |
| Metallothionein 1M (MT1M) | MT1 isoform - cytoprotective against oxidative stress, inflammation, and DNA damage. |
| Solute Carrier Family 14 Member 1 (SLC14A1) | Higher transcription identified in undifferentiated MSCs. |
| Solute Carrier Family 7 Member 11 (SLC7A11) | Antioxidant effector. |
| Superoxide Dismutase 3 (SOD3) | Antioxidant effector. |
| Secreted Phosphoprotein 1 (SPP1) | Osteogenic marker. |
| Tumour Necrosis Factor Receptor Superfamily Member 1B (TNFRSF1B) | Inflammation-associated effector. |
| Tumour Necrosis Factor Superfamily Member 12 (TNFSF12) | Pro-inflammatory cytokine mediating inflammatory stromal activation.<br>Expressed from myofibroblasts. |
| Tumour Necrosis Factor Superfamily Member 15 (TNFSF15) | Expression identified in prostate CAFs. |

**Table S4.**
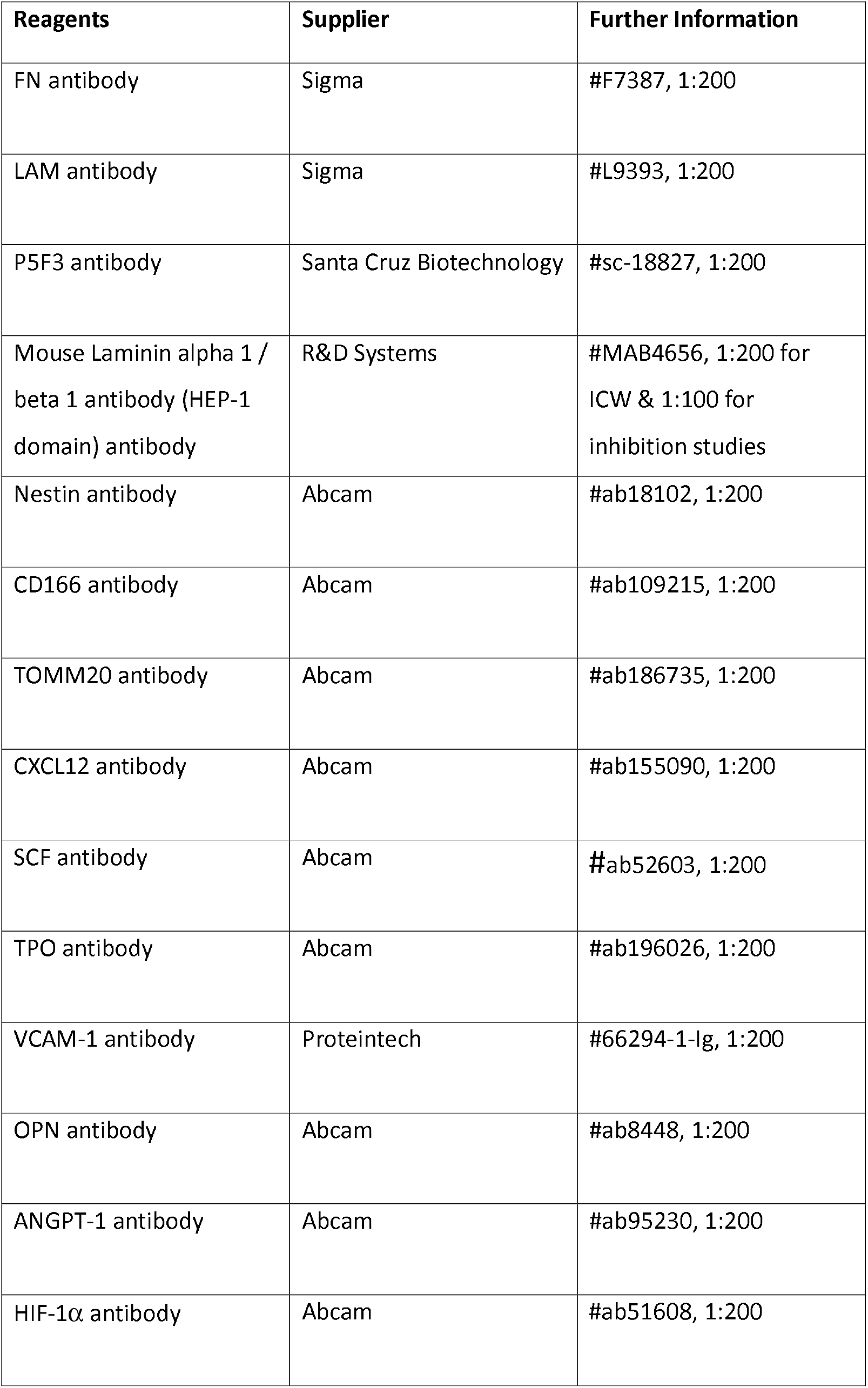
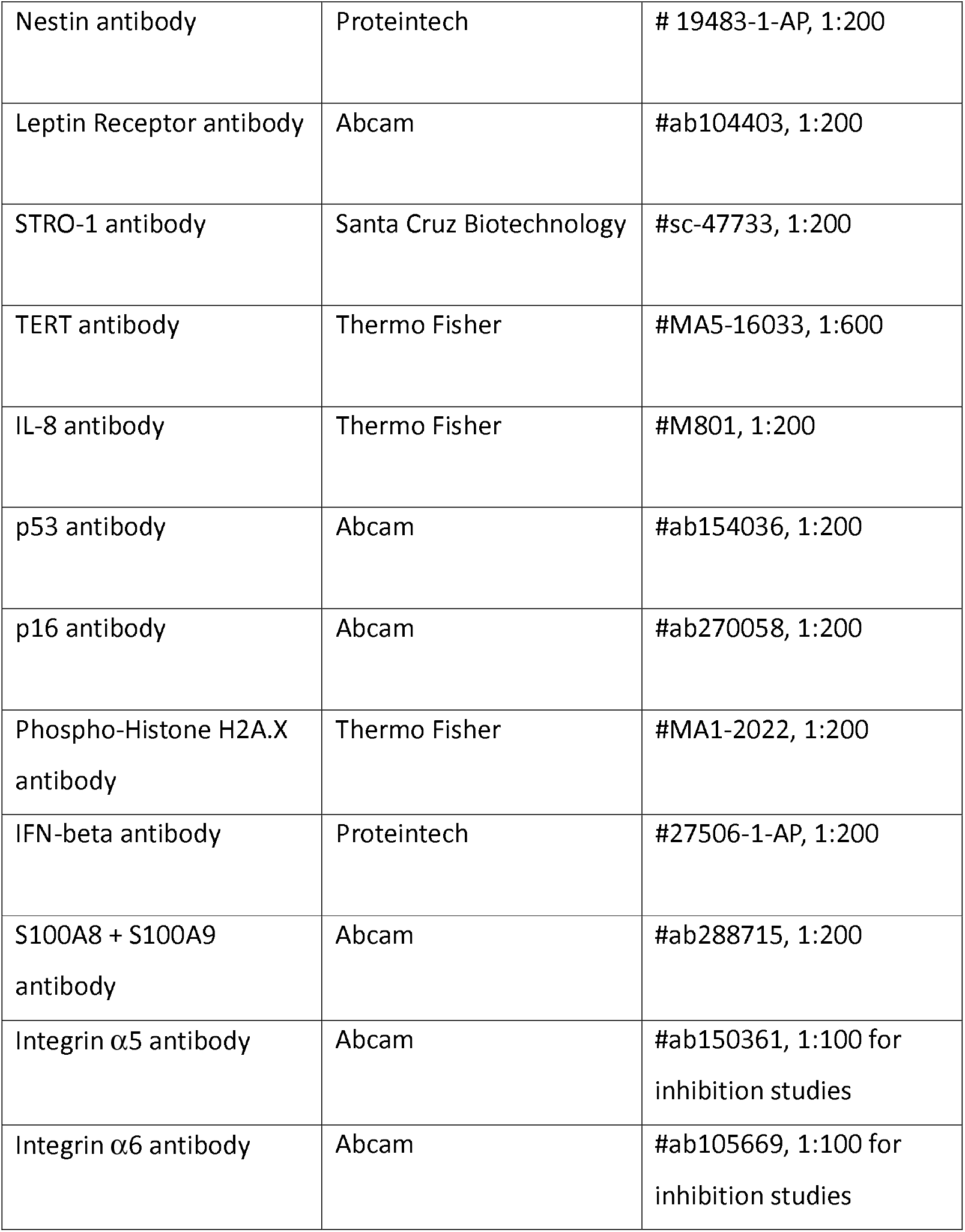
List of general antibodies used.

| Reagents | Supplier | Further Information |
|---|---|---|
| FN antibody | Sigma | #F7387, 1:200 |
| LAM antibody | Sigma | #L9393, 1:200 |
| P5F3 antibody | Santa Cruz Biotechnology | #sc-18827, 1:200 |
| Mouse Laminin alpha 1 / beta 1 antibody (HEP-1 domain) antibody | R&D Systems | #MAB4656, 1:200 for ICW & 1:100 for inhibition studies |
| Nestin antibody | Abcam | #ab18102, 1:200 |
| CD166 antibody | Abcam | #ab109215, 1:200 |
| TOMM20 antibody | Abcam | #ab186735, 1:200 |
| CXCL12 antibody | Abcam | #ab155090, 1:200 |
| SCF antibody | Abcam | #ab52603, 1:200 |
| TPO antibody | Abcam | #ab196026, 1:200 |
| VCAM-1 antibody | Proteintech | #66294-1-Ig, 1:200 |
| OPN antibody | Abcam | #ab8448, 1:200 |
| ANGPT-1 antibody | Abcam | #ab95230, 1:200 |
| HIF-1α antibody | Abcam | #ab51608, 1:200 |
| Nestin antibody | Proteintech | # 19483-1-AP, 1:200 |
| Leptin Receptor antibody | Abcam | #ab104403, 1:200 |
| STRO-1 antibody | Santa Cruz Biotechnology | #sc-47733, 1:200 |
| TERT antibody | Thermo Fisher | #MA5-16033, 1:600 |
| IL-8 antibody | Thermo Fisher | #M801, 1:200 |
| p53 antibody | Abcam | #ab154036, 1:200 |
| p16 antibody | Abcam | #ab270058, 1:200 |
| Phospho-Histone H2A.X antibody | Thermo Fisher | #MA1-2022, 1:200 |
| IFN-beta antibody | Proteintech | #27506-1-AP, 1:200 |
| S100A8 + S100A9 antibody | Abcam | #ab288715, 1:200 |
| Integrin α5 antibody | Abcam | #ab150361, 1:100 for inhibition studies |
| Integrin α6 antibody | Abcam | #ab105669, 1:100 for inhibition studies |

**Table S5.** List of flow cytometry antibodies.

| Reagents | Supplier | Further Information |
|---|---|---|
| CD7 Monoclonal Antibody (4H9), PerCP-eFluor™ 710, eBioscience™ | Thermo Fisher | #46-0078-42, 1 µl / 100 µl of flow buffer |
| CD271 (NGF Receptor)-Super Bright™ 600 | Thermo Fisher | #63-9400-42, 1 µl / 100 µl of flow buffer |
| CD29 (Integrin beta 1), Super Bright™ 702 | Thermo Fisher | #67-0299-42, 1 µl / 100 µl of flow buffer |
| CD41/CD61-Alexa Fluor™ 488 | Thermo Fisher | #MA5-44123, 1 µl / 100 µl of flow buffer |
| CD16 Monoclonal Antibody (eBioCB16 (CB16)), eFluor™ 506, eBioscience™ | Thermo Fisher | #69-0168-42, 1 µl / 100 µl of flow buffer |
| CD45RA Monoclonal Antibody (HI100), Super Bright™ 600, eBioscience™ | Thermo Fisher | #63-0458-42, 1 µl / 100 µl of flow buffer |
| CD90 (Thy-1) Monoclonal Antibody (eBio5E10 (5E10)), Alexa Fluor™ 700, eBioscience™ | Thermo Fisher | #56-0909-42, 1 µl / 100 µl of flow buffer |
| CD73 Monoclonal Antibody (AD2), PerCP-eFluor™ 710, eBioscience™ | Thermo Fisher | #46-0739-42, 1 µl / 100 µl of flow buffer |
| CD105 (Endoglin) Monoclonal Antibody (SN6), APC-eFluor™ 780, eBioscience™ | Thermo Fisher | #47-1057-42, 1 µl / 100 µl of flow buffer |
| CD146 Monoclonal Antibody (P1H12), PE-Cyanine7, eBioscience™ | Thermo Fisher | #25-1469-42, 1 µl / 100 µl of flow buffer |
| CD38 Monoclonal Antibody (HB7), PE-Cyanine7, eBioscience™ | Thermo Fisher | #25-0388-42, 1 µl / 100 µl of flow buffer |
| CD14 Monoclonal Antibody (61D3), FITC, eBioscience™ | Thermo Fisher | #11-0149-42, 1 µl / 100 µl of flow buffer |
| CD19 Monoclonal Antibody (HIB19), FITC, eBioscience™ | Thermo Fisher | #11-0199-42, 1 µl / 100 µl of flow buffer |
| CD43 Monoclonal Antibody (eBio84-3C1 (84-3C1)), FITC, eBioscience™ | Thermo Fisher | #11-0439-42, 1 µl / 100 µl of flow buffer |
| CD45 Monoclonal Antibody (HI30), FITC, eBioscience™ | Thermo Fisher | #11-0459-42, 1 µl / 100 µl of flow buffer |
| HLA-DR Monoclonal Antibody (L243), FITC, eBioscience™ | Thermo Fisher | #11-9952-42, 1 µl / 100 µl of flow buffer |
| Super Bright Complete Staining Buffer, eBioscience™ | Thermo Fisher | #SB-4401-42, 1 µl / 100 µl of flow buffer |
| CD34-APC | Miltenyi Biotec | #130-113-738, 1 µl / 100 µl of flow buffer |
| CD45-APC-Vio® 770 | Miltenyi Biotec | #130-110-635, 1 µl / 100 µl of flow buffer |

